# Fibroblasts as an in vitro model of circadian genetic and genomic studies: A temporal analysis

**DOI:** 10.1101/2023.05.19.541494

**Authors:** Marcelo Francia, Merel Bot, Toni Boltz, Juan F. De la Hoz, Marco Boks, René Kahn, Roel Ophoff

## Abstract

Bipolar disorder (BD) is a heritable disorder characterized by shifts in mood that manifest in manic or depressive episodes. Clinical studies have identified abnormalities of the circadian system in BD patients as a hallmark of underlying pathophysiology. Fibroblasts are a well-established *in vitro* model for measuring circadian patterns. We set out to examine the underlying genetic architecture of circadian rhythm in fibroblasts, with the goal to assess its contribution to the polygenic nature of BD disease risk. We collected, from primary cell lines of 6 healthy individuals, temporal genomic features over a 48 hour period from transcriptomic data (RNA-seq) and open chromatin data (ATAC-seq). The RNA-seq data showed that only a limited number of genes, primarily the known core clock genes such as *ARNTL*, *CRY1*, *PER3*, *NR1D2* and *TEF* display circadian patterns of expression consistently across cell cultures. The ATAC-seq data identified that distinct transcription factor families, like those with the basic helix-loop-helix motif, were associated with regions that were increasing in accessibility over time. Whereas known glucocorticoid receptor target motifs were identified in those regions that were decreasing in accessibility. Further evaluation of these regions using stratified linkage disequilibrium score regression (sLDSC) analysis failed to identify a significant presence of them in the known genetic architecture of BD, and other psychiatric disorders or neurobehavioral traits in which the circadian rhythm is affected. In this study, we characterize the biological pathways that are activated in this *in vitro* circadian model, evaluating the relevance of these processes in the context of the genetic architecture of BD and other disorders, highlighting its limitations and future applications for circadian genomic studies.

## BACKGROUND

It is estimated that the lifetime worldwide prevalence of bipolar disorder (BD) is 1% (Moreira et al. 2017), with an estimated heritability of 60-85% (Song et al. 2015); (Bienvenu, Davydow, and Kendler 2011). Genome-wide association studies (GWAS) of BD are showing a highly polygenic genetic architecture of disease susceptibility with common genetic variants explaining 20% of the heritability (Stahl et al. 2019); (Mullins et al. 2021). BD is primarily characterized by shifts in mood, which result in manic or depressive episodes. Clinical studies have associated abnormalities of the circadian system in Bipolar disorder type 1 (BD1) patients as a hallmark component of its pathophysiology, with disturbed sleep quality being identified as an early symptom of manic episodes (Leibenluft et al. 1996). Furthermore, dysregulation of sleep and wake cycles during manic episodes include sleep abnormalities such as decrease in total sleep time, delta sleep, and REM latency (Levenson and Frank 2011). These abnormalities have also extended to other circadian regulated systems such as cortisol levels. Both differences at morning levels of cortisol within BD subjects when compared to controls (Girshkin et al. 2014), as well as higher cortisol levels prior to a manic episode (van den Berg et al. 2020) have been reported. Despite these findings, the precise mechanisms of altered circadian rhythms in BD remain unclear.

The circadian rhythms synchronize physiological processes with the environment, creating and maintaining an internal 24 hour cycle. The main controller of the circadian cycle in mammals is the suprachiasmatic nucleus (SCN), a brain region located in the basal hypothalamus. It receives environmental cues, also called zeitgebers, such as light information from the retina which is relayed using synaptic and hormonal signaling (Le Minh et al. 2001) to the rest of the central nervous and peripheral systems. At the molecular level, the circadian machinery within every cell (Schibler and Sassone-Corsi 2002) consists of multiple transcriptional feedback loops, where core circadian genes *BMAL1* and *CLOCK* induce the expression of their own repressors, *PER1, PER2, PER3* and *CRY1, CRY2*. These genes modulate different layers of gene expression, from modifying the chromatin landscape to make certain regions of the genome more or less accessible (Menet, Pescatore, and Rosbash 2014), to post-transcriptional modifications altering the function of the associated proteins at specific times during the day (Robles, Humphrey, and Mann 2017). Although disruptions in the circadian rhythms have been associated with neuropsychiatric traits, specifically in mood disorders (Walker et al. 2020), the direct interactions between them, as well as the contributions from genomic loci, are to be elucidated.

The localization of the SCN makes direct interaction and collection in humans impossible, with researchers instead using peripheral fibroblast cells to study the molecular and genetic components of this system (Yamazaki and Takahashi 2005). These cells receive cortisol as a circadian signal from the SCN, through the hypothalamic-pituitary-adrenal axis (HPA). In order to study circadian rhythms using cell cultures, the cells need to be synchronized. One approach for this is treating the cells with dexamethasone, which elicits rhythm synchronization between the cells in a culture (Yamazaki and Takahashi 2005). Dexamethasone binds to the glucocorticoid receptor, acting on the same pathways through which cortisol regulates circadian rhythms *in vivo*. This synchronization method has been used in conjunction with luciferase bioluminescence reporter assays to study the molecular dynamics of selected circadian genes *in vitro (Nakahata et al. 2006)*. Studies using these systems have been applied to both sleep disorders and BD. Although researchers were able to find differences in the period of expression of circadian genes in sleep disorders (Hida et al. 2017), similar studies using cells derived from BD1 patients were unable to detect significant (S. Yang et al. 2009) or replicable(McCarthy et al. 2013) differences.

Here we examine the genomic components of circadian related genetic regulation and general biology that this *in vitro* fibroblast model captures, and assess whether these features relate to the genetic architecture of BD susceptibility. For this, we collected longitudinal temporal sequencing data of both gene expression and accessible chromatin regions. The temporal gene expression was used to identify genes that display circadian oscillations and are under glucocorticoid control, as well as genes with distinct temporal patterns representing other biological pathways. The temporal accessible chromatin data was used to identify regions of the genome and associated transcription factor motifs that are implicated in the temporal regulation of gene expression. Finally, we examined whether the genomic regions showing temporal transcriptomic and epigenomic circadian profiles in primary cultures of fibroblasts were enriched in genetic association signals of BD or other related psychiatric and sleep-related phenotypes.

## RESULTS

### Temporal RNA-seq captures genes with distinct longitudinal expression patterns

Outside of the subset of genes that compose the core circadian transcriptional feedback loop, most rhythmic genes are tissue specific (R. Zhang et al. 2014). Within fibroblasts, we aimed to identify the overall longitudinal patterns of all the genes that are temporally regulated and classify them based on their temporal features. For this purpose, we collected RNA-seq data every 4 hours for a 48-hour period, from cell cultures of 6 human primary fibroblasts that were derived from a skin biopsy of subjects with no psychiatric disorders. To select these subjects, we confirmed that their cell lines displayed measurable circadian oscillations via a bioluminescence assay (Supplementary figure 2). After quality control, the temporal RNA-seq dataset consisted of n=11,004 genes. We used a cubic spline regression model to identify genes that had a significant effect of time in their expression ((Wang, Ke, and Brown 2003); (Qin and Guo 2006); (Madden et al. 2017)). This approach identified n=2,767 (∼25%) genes with significant evidence (False discovery rate (FDR) < 0.05) for temporal changes of gene expression levels. To cluster these genes according to distinct temporal patterns we applied the Weighted Gene Co-Expression Network Analysis (WGCNA)(Langfelder and Horvath 2008), which identifies genes with highly correlated expression levels. WGCNA produced 11 modules with eigengene values that captured the principal time patterns present in the expression of these genes (i.e: temporal modules; Supplementary figure 3). Gene ontology (GO) analysis of WGCNA modules with MetaScape (Zhou et al. 2019) highlighted specific cell processes associated with distinct temporal patterns among 11 of these modules. Figure 1 depicts the eigengene values of 4 temporal modules with significant enrichment of GO terms (FDR adjusted by Benjamini-Hochberg method). Genes in the turquoise module, which show a linear decrease in expression over time, had GO terms for supramolecular fiber organization (p= 1e-15) and mRNA splicing via spliceosome (p= 2.5e-13). In compasion, genes in the blue module, which show a linear increase in expression, had a GO term for cellular response to hormone stimulus (p= 1.3e-9). The genes in the black module, which show an increase in expression that plateaus by the 16 hour time point (28 hours after dexamethasone treatment), were enriched for chromatin organization(p= 1e-12) and transcription elongation by RNA polymerase II (p= 7.9e-8) GO terms. The genes in the brown module, which show an expression pattern opposite of the black module, had a GO term for intracellular protein transport (p= 1e-18). Lastly, the purple module, which has genes with a peak expression at the 12 hour time point (24 hours after dexamethasone treatment), had GO terms for cell division (p= 1e-67) and mitotic cell cycle (p= 1e-60). The complete results of GO analysis for all the WGCNA modules are available in Supplementary Table 1.

**Figure 1.**
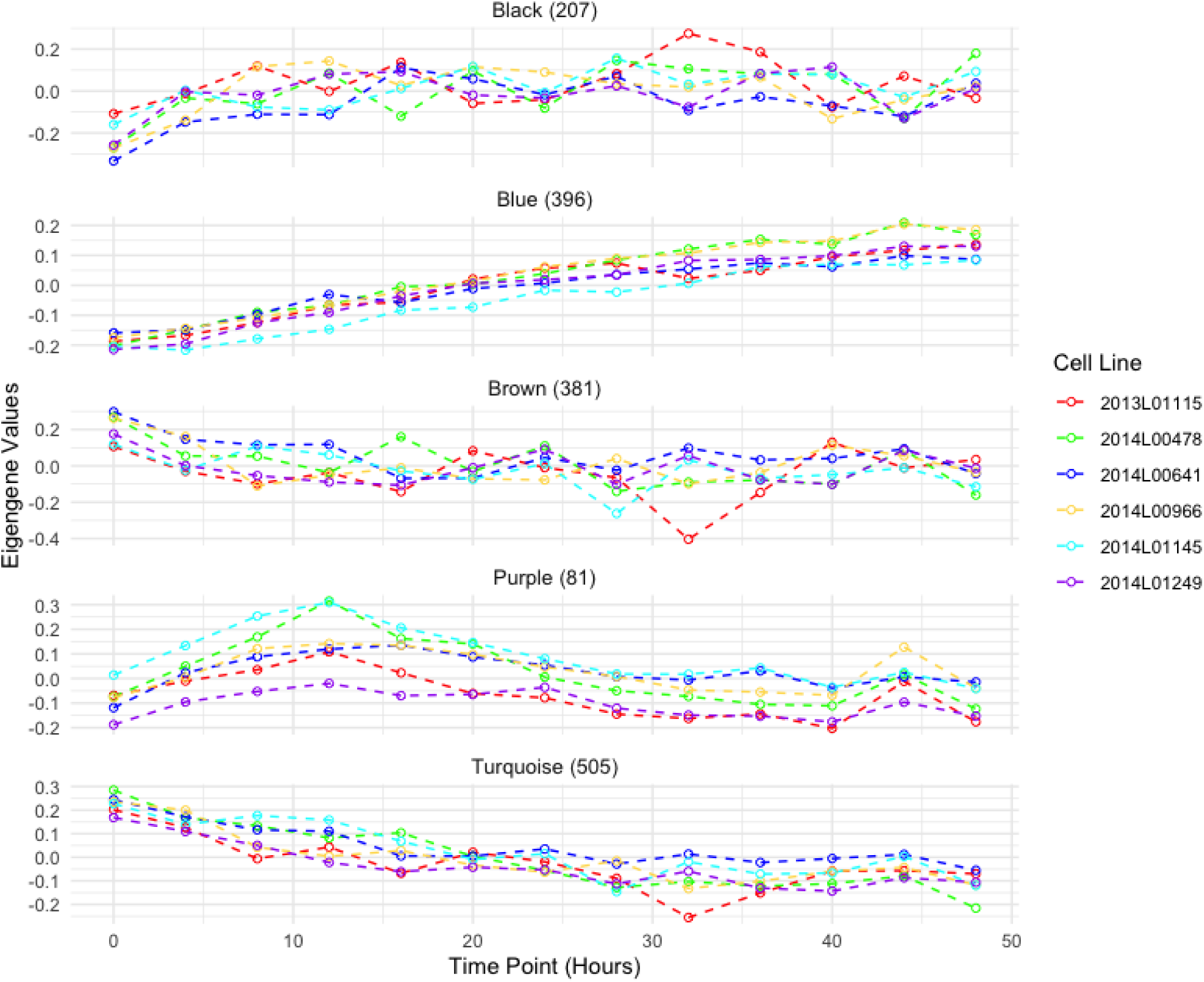
Eigengene values for RNA-seq modules obtained from WGCNA. **Description:** Eigengene modules from WGCNA of the longitudinal temporal expression patterns of RNA-seq data collected every 4 hours for a 48 hour period. Each color represents a fibroblast cell culture from a different individual. Module names were assigned by WGCNA. The number of genes assigned per module is indicated next to the module name.

WGCNA results did not yield a module of co-expressed genes with eigengene values representative of oscillating 24 hrs cycles resembling a circadian rhythm, nor were circadian related functional enrichment of GO terms found in any of the modules. Next we focused on closely inspecting known circadian genes for skin fibroblasts, identified by a previous *in vivo* array-based gene-expression circadian study on human skin cells (Del Olmo et al. 2022). Out of the 1,439 circadian genes reported in that study, we identified 267 genes in our dataset with significant changes in expression over time (Figure 2A). Using the circadian detection tools JTK Cycle (Hughes, Hogenesch, and Kornacker 2010), LS (Glynn, Chen, and Mushegian 2006), ARSER (R. Yang and Su 2010), Metacycle (Wu et al. 2016) and RAIN (Thaben and Westermark 2014), we aimed to detect significant oscillations within these putative circadian genes in the complete temporal RNA-seq dataset. Among these methods, only JTK and ARSER identified significant periodic expression patterns (after Benjamini-Hochberg correction of 0.05) for the circadian gene *NR1D2*, and further only ARSER identified significant oscillations for 73 genes. However, the predicted period differed between the methods. For example, JTK predicted a period of 27.6 hours for *NR1D2*, while ARSER predicted 24.7 hours (Supplementary table 2 and supplementary files). Therefore instead of using these circadian detection tools, we applied smoothing-splines mixed effect models using the R package "sme" (Berk 2018) to model the temporal features of these circadian genes (Figure 2B and Supplementary Figure 4). These models showed that for some of these circadian genes, such as *CRY2* and *NFIL3*, the circadian expression pattern is only present in some of the cell cultures, whereas for genes such as *NR1D2* and *TEF*, the circadian pattern is ubiquitous across cell cultures from different individuals. The fitted models for these circadian genes were then used in time warping analysis to group genes with known expression dynamics (Figure 2C and Supplementary figure 4). From these expression patterns, we corroborate that *NR1D2* expression follows its inhibition effect on *ARNTL* and *CRY1 (Rijo-Ferreira and Takahashi 2019).* Similarly, *PER3* expression follows its inhibition effect with *ARNTL*. Despite observing similar expression patterns in *PER2* and *PER3*, these were not consistent across individuals (Supplementary figure 4). While the expected inhibition relationship between *CRY1* and *ARNTL* was not present (Supplementary figure 5), this pattern of expression was also reported in the circadian dataset that was used as reference (Del Olmo et al. 2022).

**Figure 2.**
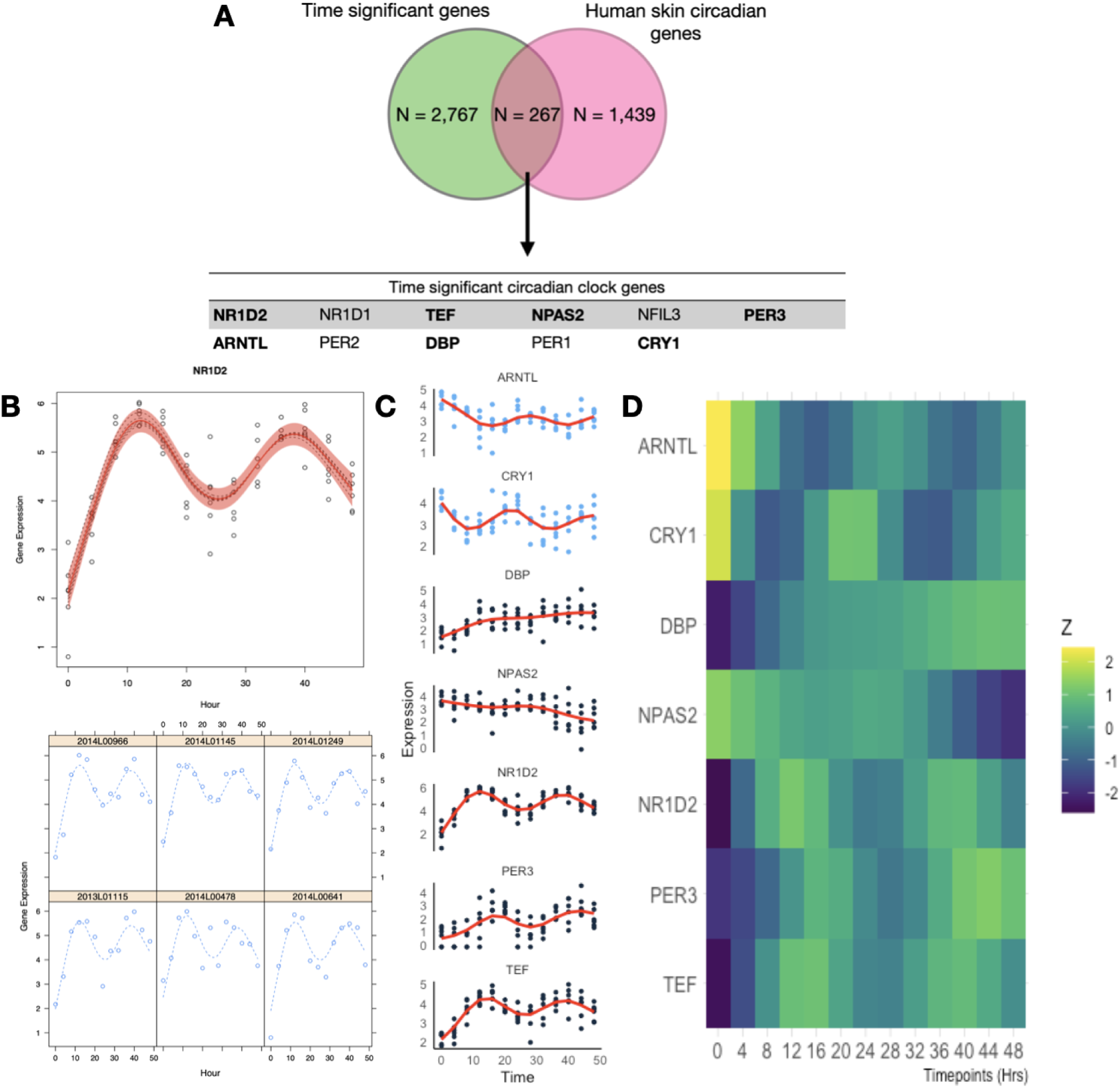
Expression patterns and mixed non-linear modeling of circadian genes. **Description A**. Overlap of the genes that were found to have a significant effect of time in their expression as well as being previously identified as circadian within the skin tissue (Del Olmo et al. 2022). In bold are those genes that displayed expression patterns consistent with circadian rhythms. **B.** Example of smoothing-splines mixed-effect model of the genes that displayed circadian oscillations and their known redundant gene partners. The red area indicates the 95 percent confidence interval. Gene expression values are presented as lCPM (log of counts per million) **C.** Dynamic time warp clustering results of the 9 circadian clock genes identified in this *in vitro* model. Due to the differences in magnitude of gene expression across circadian genes, expression values were z-scored. Clustering results for all 267 genes can be found in the supplementary material.

### Temporal open chromatin levels measured by ATAC-seq highlights potential regulatory regions and transcription factor binding sites

To identify regions of the genome associated with the regulation and downstream effects of circadian genes, we collected ATAC-seq data following the same temporal design as with the RNA-seq dataset. Quality control metrics such as fraction of reads in peaks and transcription starting site for these samples is available in the supplementary material. After removing a cell line that did not pass quality control, we merged all overlapping regions of open chromatin, also known as peaks, across samples and time points as described previously (Keele et al. 2020), to define a common set of ATAC-seq signals (n = 126,057). We then used cubic spline regression models to identify peaks that have a significant change in accessibility over time. This approach yielded n=7,568 (6%) time significant peaks, which were functionally annotated using ChipSeeker (Yu, Wang, and He 2015), a software that annotates peaks with the nearest gene and genomic regions (Supplementary figure 8B). Peaks with significant changes in accessibility over time showed a similar genomic distribution as the full dataset (Supplemetary figure 8A). Following the approach for the RNA-seq data, we applied WGCNA to cluster peaks with similar temporal patterns of accessibility changes (Supplementary figure 7).

WGCNA identified 4 different modules for the temporal patterns of chromatin accessibility, however the main pattern that characterizes these modules is an overall increase or decrease in accessibility. One module captured all the regions that were decreasing in accessibility (Figure 3A), comprising 4,435 peaks. The other 3 modules showed regions increasing in accessibility. Individual motif enrichment analysis conducted with HOMER (Heinz et al. 2010, Yan et al. 2020), showed similar enrichment across these modules, therefore we combined them into a single cluster of regions increasing in accessibility, in total 3,133 peaks. Regions that were decreasing in accessibility over time (Figure 3A) had motif sequences for Fos (p= 1e-1047), Fra1 (p= 1e-1041), ATF3 (p= 1e-1034), BATF (p= 1e-1002), Fra2 (p= 1e-996), AP-1 (p= 1e-976), Jun-AP1 (p= 1e-681), Bach2 (p= 1e-330) and JunB (p= 1e-1001). Most of these transcription factors are part of the AP-1 transcription complex. Regions that were increasing in accessibility over time (Figure 3B) had motif sequences for BHLHA15 (p= 1e-201), TCF4 (p= 1e-180), NeuroG2 (p= 1e-160), Twist2 (p= 1e-160), Pitx1 (p= 1e-186), Atoh1 (p= 1e-162), Tcf21(p= 1e-147), Olig2 (p= 1e-130), ZBTB18 (p= 1e-139) and NeuroD1 (p= 1e-123).

**Figure 3.**
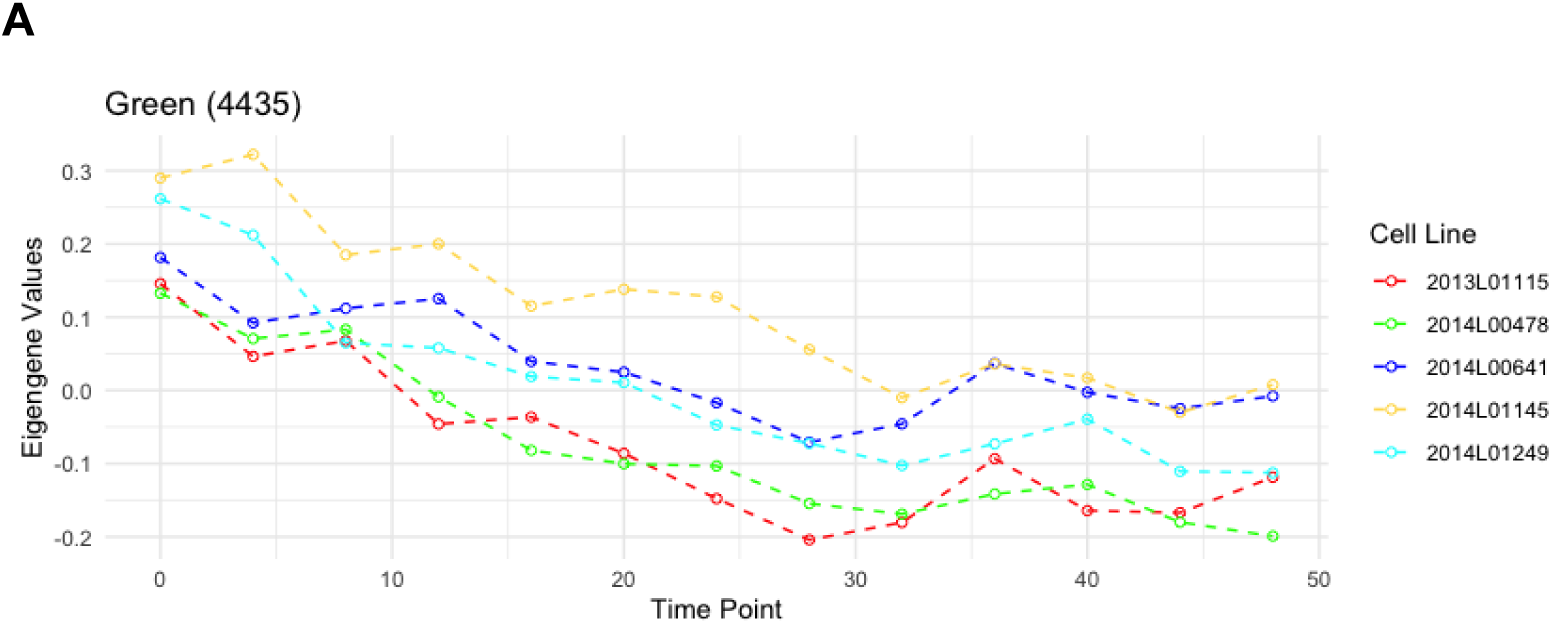
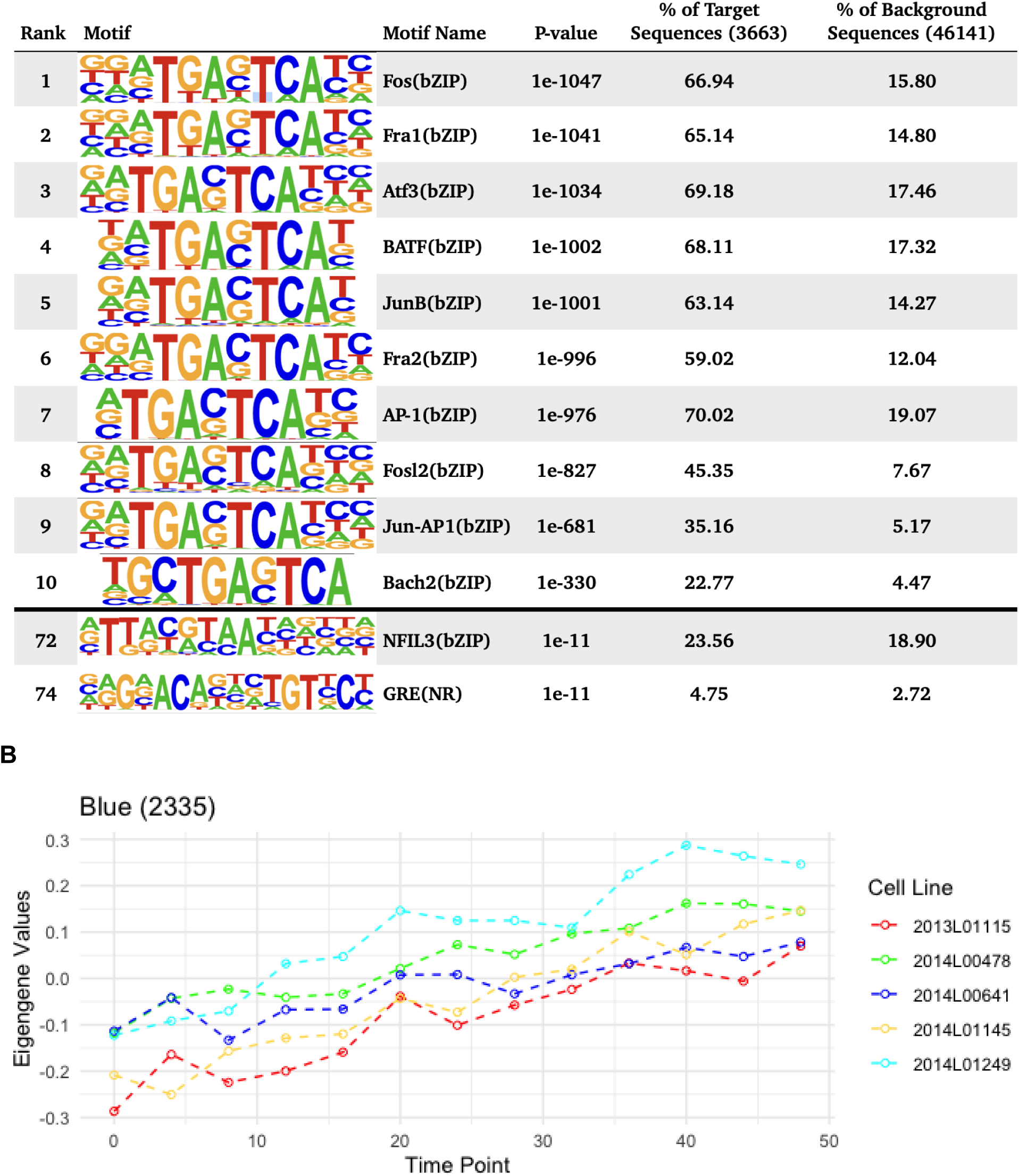
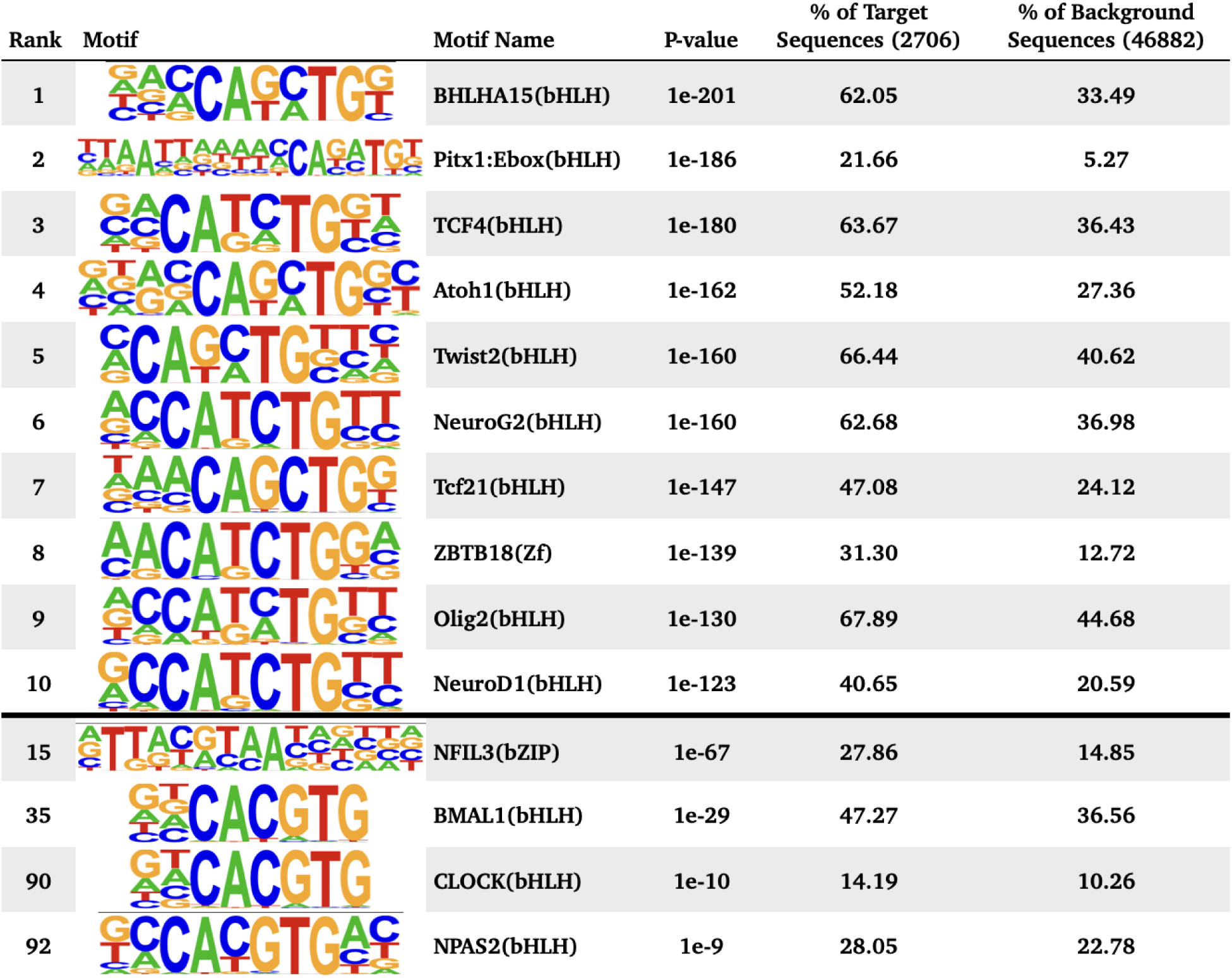
Motif enrichment analysis of time significant peak regions A. **description:** Motif enrichment analysis results after combining all open chromatin regions that followed a similar change in accessibility over time, including the top 10 motifs as well as motifs associated with circadian genes and glucocorticoid response. P-values were confirmed as significant after the Benjamini adjustment cutoff of 1% FDR. **A**. Eigengene values for the Green WGCNA module and motif enrichment analysis of the associated decreasing in accessibility regions. **B**. Eigengene values for the Blue module and motif enrichment analysis of the associated peak regions, including HOMER results for all peak regions decreasing in accessibility.

These are dimerizing transcription factors that have the basic helix-loop-helix protein structural motif. In both types of regions HOMER identified the binding sequence of the glucocorticoid response element (GRE), although the rank for the GRE motif in regions that were decreasing in accessibility was higher. For the known circadian transcription factors, HOMER identified significant enrichment of the binding sequences for BMAL1 (p= 1e-29), NPAS2 (p= 1e-9), CLOCK (p= 1e-10), particularly within regions that had increasing accessibility over time. For the regions with decreasing accessibility over time, HOMER identified enrichment of NFIL3 (p= 1e-11).

### Stratified Linkage Disequilibrium Score Regression (sLDSC) analysis

Functional annotation of the ATAC-seq dataset showed that approximately one third of the peak regions identified are located in distal intergenic regions, with unknown functions. Furthermore, it also showed that these regions displaying transient changes in chromatin state are located across the entire genome. To examine whether these open chromatin regions highlighted in our study are enriched for genetic susceptibility of BD and other neuropsychiatric traits, we used sLDSC (stratified linkage disequilibrium analysis)(Finucane et al. 2015) to calculate the partitioned heritability of these features. For this approach we used published Psychiatric Genomics Consortium (PGC) summary statistics for BD (Mullins et al. 2021), ADHD (Attention-Deficit/HyperactivityDisorder) (Demontis et al. 2019), schizophrenia (Trubetskoy et al. 2022), PTSD (Post-traumatic stress disorder) (Nievergelt et al. 2019), MDD (Major depression disorder) (Howard et al. 2019), insomnia (Watanabe et al. 2022), and the circadian trait of morningness (Jones et al. 2016). We used the temporally significant ATAC-seq regions with 1 kilobases (kb) and 10 kb genomic windows in both downstream and upstream directions for each region. These ATAC-seq defined annotations were tested jointly with the baseline annotations included with sLDSC (Finucane et al. 2015). Figure 4 shows the enrichment for the traits tested from the ATAC-seq regions annotations as well as the baseline annotations (Full enrichment results are provided in the Supplementary Material). Among these, only the ATAC-seq regions that were decreasing in accessibility had a nominally significant (p value = 0.00463) less than expected presence for ADHD, and this effect was not present when the regions are extended by either 1kb or 10kb. In comparison, baseline annotations such as conserved regions in mammals showed a significant enrichment for all the traits (ADHD p value = 7.88e-11, schizophrenia p value = 1.67e-23, BD p value = 1.14e-8, MDD p value = 5.92e-22, insomnia p value = 2.65e-15, morningness p value = 3.05e-29), except PTSD (p value = 0.073). We did not identify significant enrichment of ATAC-seq regions in the other psychiatric and behavioral traits tested, indicating that these genomic regions with temporal trends in cromatin accessibility do not play a major role to their genetic architecture.

**Figure 4.**
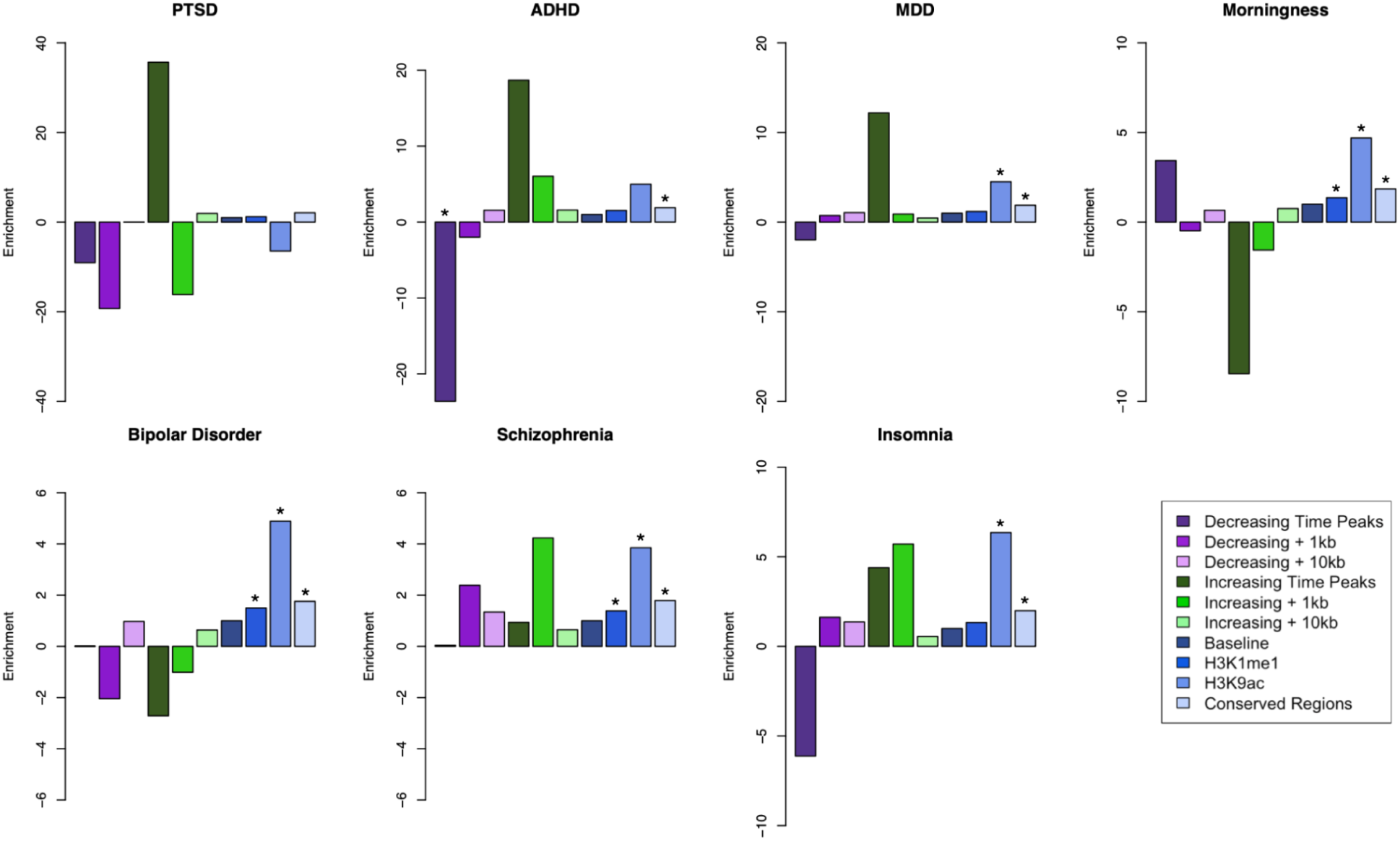
sLDSC enrichment results for psychiatric disorders and a circadian trait. **Description:** Results of partitioned sLDSC across 3 psychiatric disorders and morningness trait across different genomic annotations. Shown are the enrichment for both the temporal ATAC-seq regions and extended genome windows; as well as annotations part of the baseline model of sLDSC, such as baseline for all the annotations, histone markers H3K9ac, H3K4me1, and conserved regions in mammals. PTSD: Post-traumatic stress disorder; MDD: Major depressive disorder; ADHD: Attention-Deficit / Hyperactivity Disorder. * indicates enrichment p-value below 0.01.

## DISCUSSION

Cell cultures of peripheral tissues have been employed as models of *in vitro* circadian clock systems to study their molecular components (Balsalobre, Marcacci, and Schibler 2000) and the disorders in which they are disrupted (Kripke et al. 2009). For this particular model that uses primary cultures of fibroblasts derived from skin biopsies, we aimed to characterize the circadian features that are present at gene expression and chromatin accessibility levels, with the goal to identify the circadian genes that are engaged by this system as well as their associated regulatory genomic regions. From the longitudinal RNA-seq data we identified consistent circadian patterns of expression in a limited amount of genes such as *ARNTL, CRY1, PER3, NR1D2* (Rev-erbBeta) and TEF, but observed noticeable differences in the expression patterns between cell cultures. When compared to a recent *in vivo* circadian human skin dataset (Del Olmo et al. 2022), we identified 267 out of the 1,439 circadian genes previously identified in this tissue to have a significant effect of time in their expression. This limited overlap could indicate that this *in vitro* model for studying circadian rhythms is constrained to the circadian genes that are directly activated by a glucocorticoid-like stimulus. Glucocorticoid response elements have been identified for circadian genes such as *PER1, PER2, PER3, CRY1, CRY2, Rev-erbAlpha (NR1D1), Rev-erbBeta (NR1D2), DBP, NPAS2* and *BMAL1* (So et al. 2009). Consistent with these results we identified circadian rhythmicity in most of those genes that had previously been identified to have a glucocorticoid response element (GRE), with the exception of *NR1D1*. While we do identify expression levels from *NR1D1*, the lack of a significant circadian oscillation in comparison to the strong results from *NR1D2* could be consistent with their known redundant function for circadian rhythms (Liu et al. 2008).

We found robust circadian expression patterns for *NR1D2*. However, differences in the lenght of the predicted periods across tools indicates that estimating period duration from longitudinal RNA-seq data is not a straightforward problem. This could be due to the the small number of subjects used, leading to insufficient power. Furthermore, while ARSER identified 73 genes with significant periodic expression patterns, this software is known to have a high false positive rate in high resolution data (Wu et al. 2016). Interestingly, while the expression pattern of *CRY1* follows the expected inhibition by *NR1D2* (Chiou et al. 2016), it does not reflect the expected inhibitory action on *ARNTL*, nor the similar phase pattern with its heterodimer partner *PER3*. For the relationship with *PER3*, *CRY1* has been previously reported to have a known phase delay with the *PER* genes (*PER1,2,3*)(Fustin et al. 2009), which could be attributed to the multiple binding sites that *CRY1* has for different circadian modulators, resulting in stimulus and tissue specific temporal dynamics. Based on the time patterns, changes in the expression of *CRY1* appear to precede the expression of *ARNTL* by 4 to 8 hours. Similar expression patterns between these genes were reported in a previous *in vivo* study of human skin cells (Del Olmo et al. 2022), indicating that this model was able to replicate some of the circadian temporal dynamics seen within tissue.

ATAC-seq data can provide an untargeted yet comprehensive view of chromatin accessibility changes over time. Within this fibroblast *in vitro* model, we mainly identified genomic regions with linear increases and decreases of chromatin accessibility. Although we did not found circadian temporal patterns in chromatin accessibility, previous studies conducted *in vivo* have reported such patterns (Koike et al. 2012). Specifically, proteins such as CLOCK and BMAL1 have been found to associate and interact with chromatin remodeling and chromatin modifying enzymes (Zhu and Belden 2020), as well as act as pioneer factors by directly modifying chromatin accessibility (Menet, Pescatore, and Rosbash 2014). The absence of anticipated oscillations in chromatin within our dataset, as opposed to our observations in genes, could be due to multiple reasons. One possibility is that the mechanisms governing oscillatory chromatin changes may be exclusive to *in vivo* conditions. Under a physiological setting, cells within a tissue are exposed to multiple stimuli that act as Zeitgebers, such as sunlight (exposure), metabolic signals, temperature, and hormones like cortisol (Roenneberg and Merrow 2016). The exposure to these signals is under a rhythmic control, with levels cycling throughout the day (Chauhan et al. 2023). By using a single exposure to dexamethasone in this model, we are missing the cyclic aspect of cortisol response present *in vivo*, as well as other effects that could be due to the coupling of Zeitgebers. However, our ATAC-seq dataset does replicate previous findings on the broader role of glucocorticoids in the chromatin landscape. Within regions with decreasing accessibility post-dexamethasone, the motif enrichment analyses identified the motifs for the GRE as well as for members of the AP-1 transcriptional complex. These regions may have initially opened due to the dexamethasone treatment (for synchronization of the cells), but are closing without further continued exposure of dexamethasone. Regions with increasing accessibility may have initially closed due to the dexamethasone treatment, and this is consistent with motif profiles that were unrelated to direct glucocorticoi receptor (GR) targets (Bothe, Buschow, and Meijsing 2021). The dataset, however, lacks a Chip-seq analysis for GR occupancy, and we are limited to confirm if the identified regions are indeed due to GR activity. The strong glucocorticoid effects observed in our data underscore the need for further exploration of circadian influences on chromatin regulation in fibroblast cell culture models. Other methods for synchronizing these cells and studying the circadian rhythms, such as a switch to serum free media (Yamazaki and Takahashi 2005), Forskolin treatment (Yagita and Okamura 2000), or temperature cycles (Saini et al. 2012), could result in different types of chromatin regulation and gene expression dynamics. This study raises questions about the context-dependent nature of chromatin remodeling events and emphasizes the need to evaluate different synchronization methods to ascertain their implications for circadian rhythms.

The ATAC-seq data showed genome wide transient changes in chromatin conformation, with most of these changes occurring within regions of unknown functions. To evaluate the relevance of these genomic regions for BD and other psychiatric traits, we used partitioned sLDSC regression. This tool identifies genetic susceptibility enrichment for a particular trait across the whole genome and within specific genomic annotations. The partitioned sLDSC analysis mainly showed a significant deflation for the chromatin regions that were decreasing in accessibility over time with ADHD. Although not significant, it mirrored the enrichment for the regions that were increasing in accessibility. When expanding the genomic regions by either 1kb or 10kb both the magnitude and the significance of the enrichment are lost, indicating that this effect could be highly localized for these regions. For the other traits that we examined, the enrichment from the ATAC-seq regions were also attenuated when the genomic region was extended. These results show that these regions with linear changes in chromatin accessibility identified here may not play a relevant role for these neuropsychiatric traits. However the attenuation observed when expanding the genomic window suggests that any relevant signal may be specific to those genomic positions.

The lack of an overlap between the temporal regulatory regions identified in this study and the known genetic architecture of BD could indicate four different interpretations. First, although peripheral tissues capture the genome of an individual, they don’t recapitulate brain molecular physiology, the main tissue implicated in the pathophysiology of BD. Second, it could be that the disruptions in the circadian rhythm are not under strong genetic control and are actually influenced by other downstream processes, such as post-translational modifications and differences at the protein level. Third, the specific circadian pathways that are engaged in this *in vitro* model by dexamethasone (i.e: glucocorticoids) are not part of the genetic architecture of BD. This however does not exclude other circadian pathways that could be engaged by a different synchronizing stimulus, such as serum (Balsalobre, Damiola, and Schibler 1998), (Iyer et al. 1999). Furthermore, there are other stimuli that also act in different ways with the circadian system, such as temperature. Whereas dexamethasone acts through binding of the glucocorticoid receptor, temperature affects heat-shock proteins (Saini et al. 2012). Lastly, the disrupted circadian phenotype is an episodic state in BD patients, not a constant trait. This could indicate that rather than the regular circadian system being affected by BD, it is the ability to deal with circadian stressors and disruptors that is implicated in BD disease susceptibility.

The circadian analysis of gene expression and chromatin accessibility data faced limitations that could be attributed to variability among cell lines from different subjects. Human skin cell studies, both *in vivo (Del Olmo et al. 2022)* and *in vitro (Brown et al. 2005)*, have demonstrated that genetic differences contribute to variations in circadian gene expression’s phase and amplitude. The variability observed in this dataset was therefore not unexpected, but remains a factor to be considered for this kind of studies. Another potential source of variability was the data collection scheme, involving 13 separate cell cultures for each individual cell line. Distinctions in cell cycle state and growth rates among these cultures might have influenced the data. Previous research has shown that cell cycle and circadian rhythms are coupled processes((Nagoshi et al. 2004), (Farshadi, van der Horst, and Chaves 2020)), and that these rhythms can be impacted by cell density (Noguchi, Wang, and Welsh 2013). Our approach, utilizing a 5% FBS culture that minimizes cell growth, aimed to control for both of these factors. Our lab’s prior work also confirmed that the cell density used for our study allows for the production of rhythmic circadian cycles in these cells(Supplementary figure 2). Although various factors known to influence circadian rhythms could have contributed to the variability in this dataset, certain circadian genes appeared resilient, consistently producing rhythmic cycles across cultures and individual cell lines. This could highlight specific genes’ resilience to various sources of variation in this kind of studies.

With the knowledge of the specific features that this *in vitro* model is able to capture of the circadian system, we advise care when interpreting the results of such experiments, as they may be heavily influenced by genetics, cell culture factors, and, crucially, the circadian cycle synchronization method. This can, inadvertently, lead to a narrowing in the scope of studies of circadian rhythms in the context of neuropsychiatric traits. While the biology that this model captures after circadian synchronization induced by dexamethasone treatment does not seem to be directly involved in the known genetic architecture of BD, this model can still be applied to scientific questions that cannot be explored directly in human subjects. For instance, this model could be employed to characterize the specific biological pathways that are engaged during circadian distress. Notably, the dysregulated circadian phenotype in BD patients is characterized by episodic events rather than a static state, emphasizing the dynamic nature of the subjects. Responses to circadian distress could be directly compared between fibroblast cell lines derived from BD patients and healthy subjects. Additionally, the accessibility that this *in vitro* model provides could be used to study the effect of lithium, the most commonly used prescribed drug treatment for BD, during such circadian distress.

## ACKNOWLEDGMENTS

We would like to acknowledge the advice and expertise of Dr. Christopher Colwell for this project, and to Dr. Annet van Bergen for their work in collecting the tissue samples used in this study. We thank the donors for their willingness to provide a skin biopsy for the generation of the primary cell cultures. This study was supported by funding from the National Institutes of Health (NIH), research grants R01 MH090553, R01 MH115676, the NARSAD Distinguished Investigator Grant, (to RAO) and NS048004 T32 Training grant in Neurobehavioral genetics.

## MATERIALS AND METHODS

### Cell lines and Culture

Fibroblasts were isolated by taking skin biopsies from the nether region from subjects without known psychiatric disorders. Fibroblast cultures were established following standard procedures (Villegas and McPhaul 2005) and stored as frozen aliquots in liquid nitrogen. 6 fibroblast cell lines matched for sex, age and passage number were thawed out and grown to confluence in T75 culture flasks in standard culture media (DMEM containing 10% fetal bovine serum (FBS) and 1x Penicillin-Streptomycin).

Upon reaching confluence, 5x10^4 cells were plated per line into 13 different 6 well plates (1 well per line per plate). All 6 lines were collected in the same experiment for the RNA-seq experiment. Due to the labor-intensive nature of the ATAC protocol and the need to process cells fresh, the 6 lines were split into 2 batches, so 3 lines per batch were processed.

### Assessment of Circadian Expression *in vitro*

In order to collect RNA or cells every 4 hours for 48 hours, cells were split into two batches, which were reset 12 hours apart (see supplementary figure 9). Cells were reset 12 hours before the first collection to exclude the acute effects of dexamethasone and variation in synchronization conditions (Brown et al. 2005). 5 days after being plated the cells from batch one were synchronized by treatment with 100 nM Dexamethasone for 30 min. Cells were then washed with PBS and switched to collection media (DMEM containing 5% FBS and 1x Penicillin-Streptomycin). Lower concentration of FBS was used in this media to stop the cells from growing during the experiment, in order to keep all time points at approximately the same culture density. 12 hours later cells from batch 2 were synchronized and switched to collection media and the RNA/cell collection was started (from batch one).

### RNA and Cell collection

For the collection of RNA, cells were lysed using 350uL RLT lysis buffer from the Qiagen RNeasy mini kit. Lysed cells were then scraped off the plate, transferred to a Qiaschredder (Qiagen 79656) and centrifuged for 2 min at max speed to further homogenize. Cell lysates were kept in -80 until extraction.

For the collection of cells for the ATAC protocol, cells were dissociated using 500uL of prewarmed TrypLE (ThermoFisher 12604013) and left for 5 min at 37℃. TrypLE was inactivated using 500uL of DMEM. Cells were then counted using the Logos Biosystems LUNA-FL automated cell counter, and 50×10^4 cells were used as input for tagmentation. Tagmented DNA for library preparation was collected following the previously described protocol (Buenrostro et al. 2015).

### RNA extraction

RNA from cell lysates was extracted using the Qiagen RNeasy mini kit (Qiagen 74106). Cell lysates were extracted in a randomized order to prevent batch effects in downstream analysis. In order to collect total RNA including small RNAs, the standard extraction protocol (Purification of Total RNA from Animal Cells using Spin Technology) was adjusted by making the following changes: (i) adding 1.5 volumes of 100% ethanol, instead of 70%, after the lysis step (step 4 in handbook protocol) and (ii) adding 700 mL of buffer RWT (Qiagen 1067933) instead of the provided RW1 (step 6 in handbook protocol).

### RNA and ATAC sequencing

For the RNA sequencing, library preps were made using the Lexogen QuantSeq 3’ mRNA-Seq Library Prep Kit and sequenced with 65-base single end reads, and sequenced at a targeted depth of 3.8M reads per sample, which is well above the recommended minimum 1M reads per sample read depth for these types of libraries. ATAC seq libraries were generated following the previously described protocol (Buenrostro et al. 2015) and sequenced with 75-base double end reads, and sequenced at a targeted depth of 61M reads per sample. Library preparation and sequencing was performed at the UCLA Neuroscience Genomics Core (https://www.semel.ucla.edu/ungc). All samples were sequenced on a Illumina HiSeq 4000 sequencer.

### RNA-seq data processing and analysis

Fastqc (Andrews, 2010) software was used to assess the quality of the read files. Low quality reads were trimmed using TrimGalore and Cutadapt.

Alignment of reads was performed with the STAR(Dobin et al. 2013) software and to human gene ensembl version GrCh38. STAR was indexed to the genome using the –runMode genomeGenerate function. For aligning, STAR was run with the parameters –outFilterType BySJout –outFilterMultimapNmax 20 –alignSJoverhangMin 8 –alignSJDBoverhangMin 1 –outFilterMismatchNmax 999 --outFilterMismatchNoverLmax 0.1 --alignIntronMin 20 --alignIntronMax 1000000 --alignMatesGapMax 1000000. Samtools was used to index the aligned files from STAR. Read counts were associated with genes using featureCounts software with the NCBI GRCh38 gene annotation file.

Analysis of the RNA-seq data used the R packages limma, Glimma and edgeR, as previously described (Law et al., 2018). Genes with low read counts were removed and reads were normalized by CPM. GeneIDs were converted to Gene Symbols using the package Homo.sapiens.

WGCNA (Langfelder and Horvath 2008) software was used to classify genes with similar temporal expression patterns. WGCNA was run using a power value of 12 obtained from diagnostic plots and with the "signed" argument. MetaScape was used for Gene Ontology analysis of the resulting gene sets from WGCNA.

Following the method described in (Mei et al. 2021), MetaCycle (Wu et al. 2016) was used to run circadian detection tools such as ARSER, JTK (Hughes, Hogenesch, and Kornacker 2010), LS and metacycle. In order to integrate results from the different individuals, the function meta3d was used from the Metacycle R package. RAIN (Thaben and Westermark 2014) was run separately.

According to the EdgeR user guide, cubic splines were generated using the splines package in R, with the ns function and 5 degrees of freedom. Resulting p-values were corrected using a false discovery rate of 0.05. Significant genes were then compared to a previously published dataset of circadian human skin gene expression, resulting in 267 genes that were classified according to their time series using the “dtwclust” R package. The resulting clusters are available in the supplementary material. This analysis was also performed with only the known circadian clock genes that had consistent expression patterns across the cell-lines.

### ATAC-seq data processing and analysis

Sequenced open chromatin data from the ATAC-seq assay followed the standard ENCODE Pipeline for the identification of open chromatin regions (OCRs) of the genome. The steps included using fastqc to evaluate the quality of the sequenced library. Followed by trimming of low quality reads with Trimgalore and Cutadapt. Alignment of the raw reads data to human gene Ensembl version GRCh38 was performed using bowtie2 (Langmead and Salzberg 2012) with a 2kb insert size and allowing up to 4 alignments. Reads within black-listed regions alongside PCR duplicates were removed with samtools. MACS2 (Y. Zhang et al. 2008) software was used to identify OCRs with parameters -g hs -q 0.01 –nomodel –shift -100 –extsize 200 –keep-dup all -B. PCR. Quality control metrics for the ATAC-seq dataset such as peak counts, PCR bottlenecking coefficients, fraction of reads in peaks and enrichment of transcription starting site are provided in the supplementary material.

To compare the ATAC-seq signal across timepoints and subjects, we created a consensus bed file using the bedtools(Quinlan and Hall 2010) function merge function, combining all the overlapping peak regions across timepoints and subjects into a single file. The Featurecounts software was then used to assign read counts to those regions. Read counts were normalized by RPM.

WGCNA (Langfelder and Horvath 2008) software was used to classify peaks with similar temporal accessibility patterns. WGCNA was run using a power value of 12 obtained from diagnostic plots and with the "signed" argument, identifying 4 modules after merging.

HOMER (Heinz et al. 2010) findMotifsGenome.pllp program was used to identify enriched transcription factor motifs first individually in the peaks that belonged to the largest WGCNA modules, as well as in the resulting set of grouping all the modules that displayed a similar increasing or decreasing pattern of accessibility.

### Stratified linkage disequilibrium score regression analysis

Following the procedure described in (Ori et al. 2019), we applied an extension to sLDSR, a statistical method that partitions SNP-based heritability(h2) from GWAS summary statistics (Finucane et al. 2015). We ran sLDSR (ldsc.py –h2), using an ancestry-match 1000 Genomes Project phase 3 release reference panel, for each annotation of interest while accounting for the full baseline model, as recommended by the developers ((Finucane et al. 2015); (Gazal et al. 2017)), and an extra annotation of all the ATAC-seq detected in our in vitro model (n=3126 for peaks that were decreasing in accessibility, n=4415 for peaks increasing in accessibility), as well as extension of these regions by 1kb and 10kb genomic windows in both directions.

