## Supplementary Figure for "Fibroblasts as an in vitro model of circadian genetic and genomic studies: A temporal analysis"

**Supplementary Files Information:**

All Circadian Genes Heatmap Clusters: Directory containing heat map plots of the 12 clusters obtained from the previously identified circadian genes in this dataset

ATACQC_Scores.csv: CSV containing quality control metrics for the ATAC-seq data. First column is sample name, second if the Fractions of reads in Peaks score, third is the Non Redundant Fraction, columns 4 and 5 are the PCR bottlenecking coefficient 1 and 2.

Full_Metascape_Output.zip: Zip file containing full Metascape gene ontology analysis for the WGCNA modules identified from the RNA-seq data

GO_Results: Gene ontology results from Metascape. Last column indicates the WGCNA module.

MetaCycle Results: Directory containing results from meta2d, JTK, ARSER, LS and RAIN analysis.

SLDSC_Enrichment_Results: Directory containing enrichment results from the Stratified linkage disequilibrium score analysis for ADHD, Bipolar disorder, Schizophrenia, Insomnia, MDD, Morningness and PTSD.

TSS QC Summary: PDF file including coverage curved of nucleosome-free and nucleosome signals for each ATAC-seq sample. Transcription Start Site (TSS) Enrichment Score is indicated in bold next to the sample name.

WGCNA_ATAC_Modules_ID.csv: CSV file containing WGCNA module membership results for the ATAC-seq data.

WGCNA_RNA_Modules_ID.csv: CSV file containing WGCNA module membership results for the RNA-seq data.

### Supplementary Figure 1 Principal component analysis of RNA-seq temporal dataset

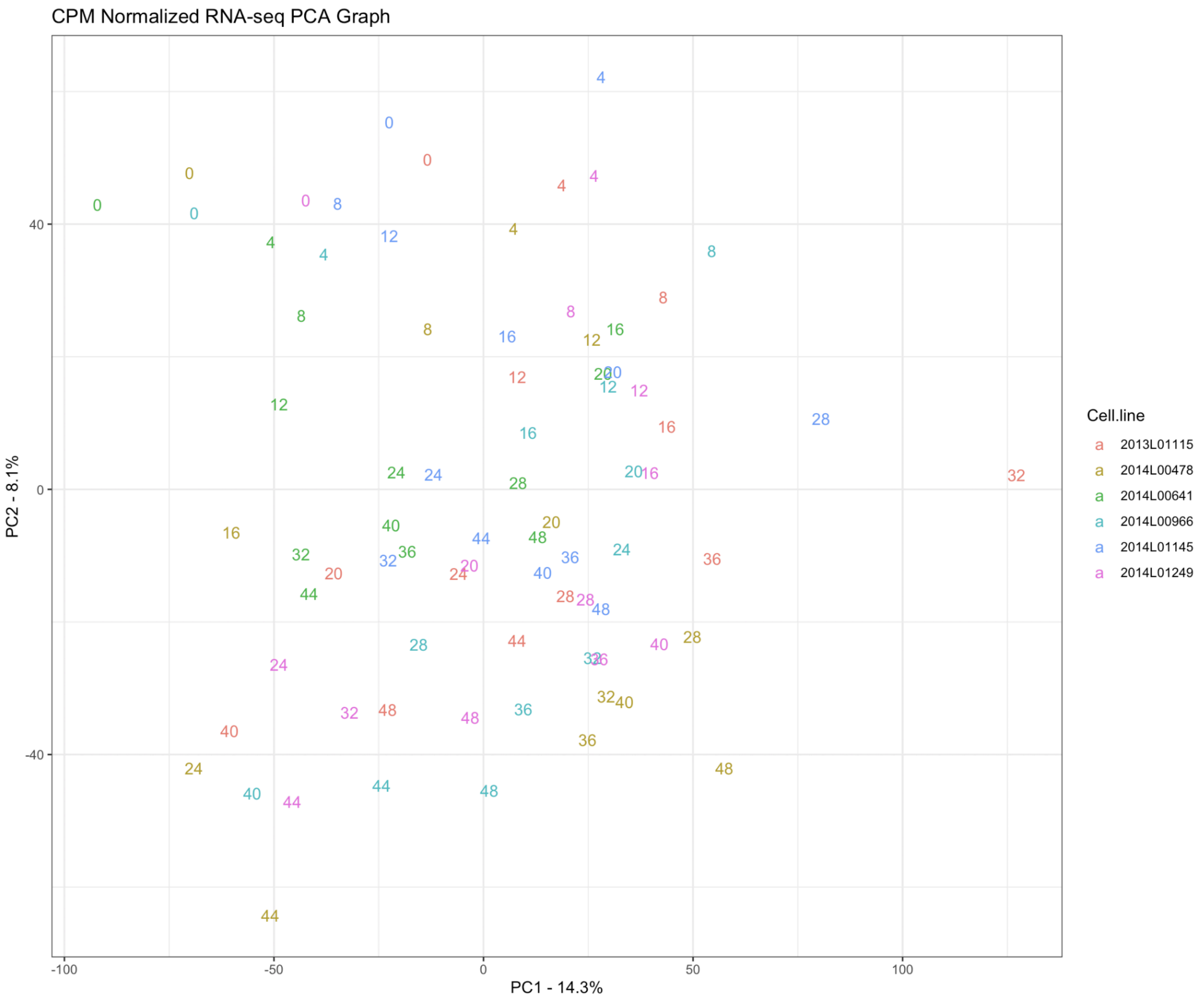


**Supplementary Figure 1 Description:** Principal component component representation of RNA-seq data after CPM(counts per million) normalization. Colors indicate each individual cell line used in this study. Numbers in the plot indicate the corresponding time point for that cell line.

**Supplementary Figure 2 Circadian-bioluminescence transduction experiment results**

**A**

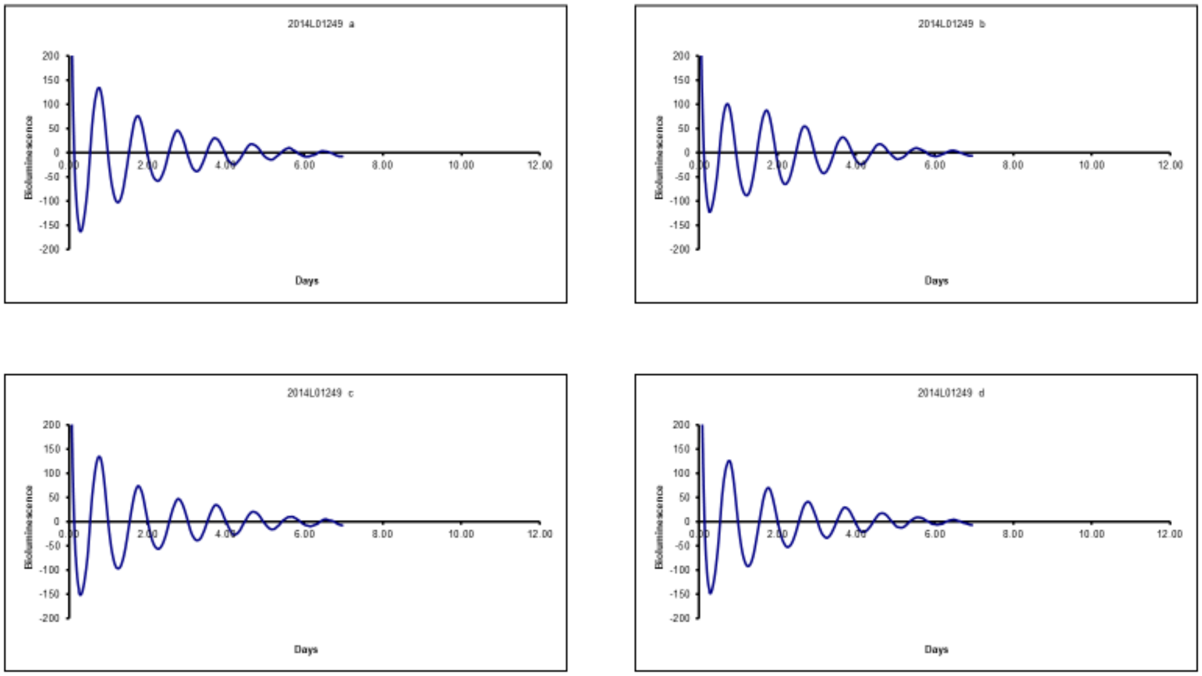


**B**

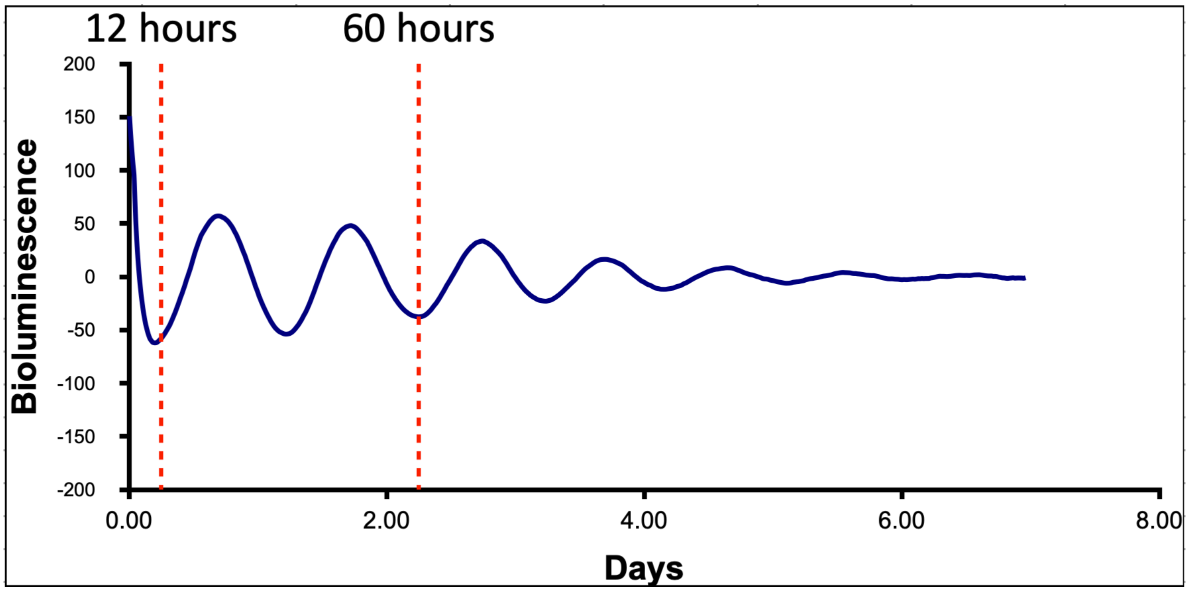


**Supplementary Figure 2 Description**: Example of circadian bioluminescence assay performed using transduced primary fibroblast cell lines with a Bmail1:luc construct, as described previously by Brown, et al. 2005 (The Period Length of Fibroblast Circadian

Gene Expression Varies Widely among Human Individuals). A. Bioluminescence measurements of cell cultures following synchronization with dexamethasone. B. Red lines indicate the time period during which RNA-seq and ATAC-seq data was collected from the cell lines.

**Supplementary Figure 3 WGCNA modules obtained from the RNA-seq temporal dataset**

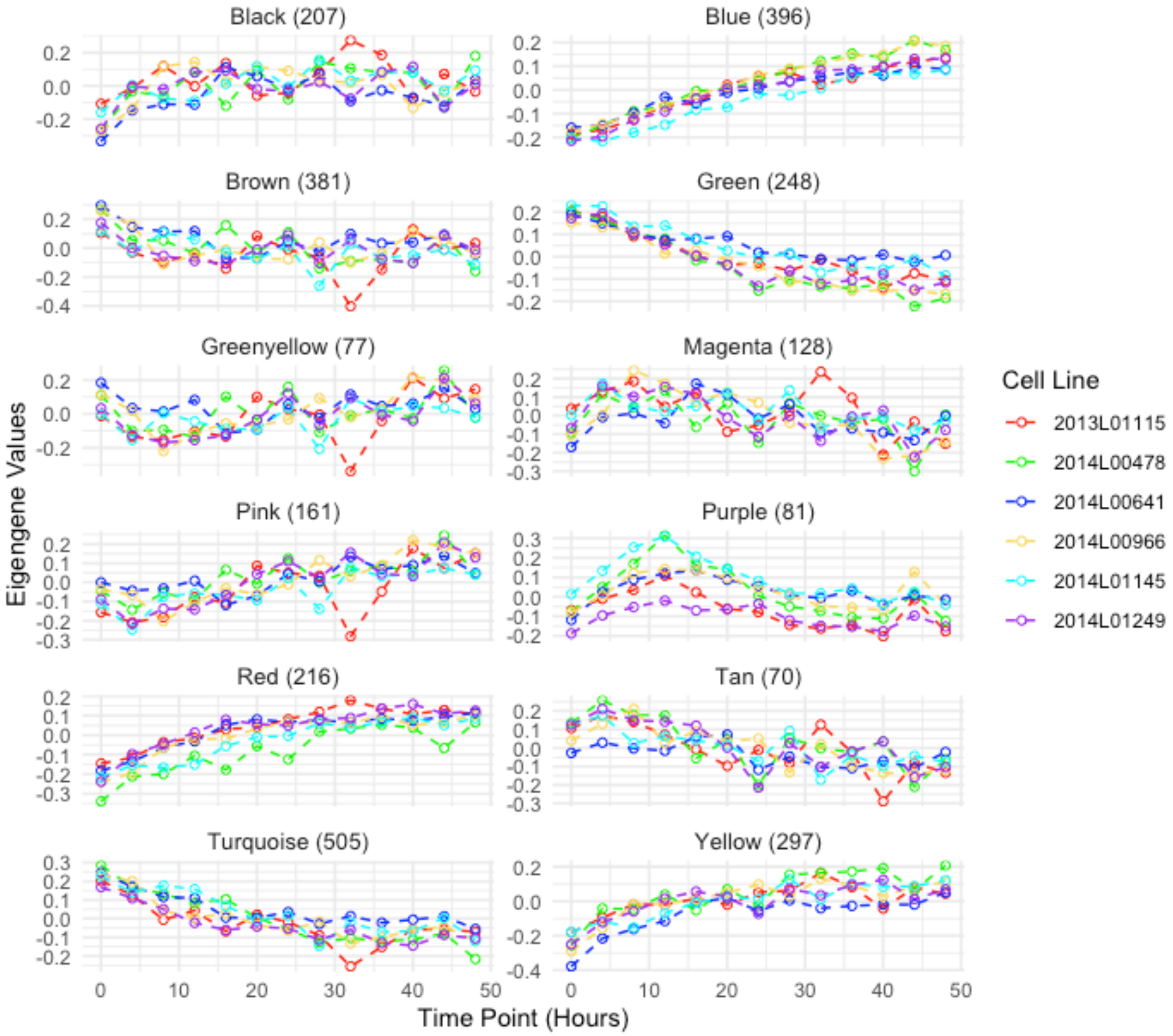


**Supplementary Figure 3 description**: WGCNA modules obtained from n=2,767 genes identified as having a significant effect of time in their expression through cubic splines modeling. Expression values are represented here as eigengene values. Names of the modules were assigned by WGCNA. The number of genes assigned per module is next to the module name.

**Supplementary Table 1 Gene Ontology Analysis of genes in eigengene modules(Full results in Full_Metascape_Output.zip)**

Gene ontology analysis using MetaScape of the genes that belong to the different modules identified by WGCNA

| Category | CategoryID | GO | Description | PARENT_GO |
|---|---|---|---|---|
| GO Biological Processes | 19 | GO:0006325 | chromatin organization | 19_GO:0009987 cellular process |
| Reactome Gene Sets | 6 | R-HSA-5617833 | Cilium Assembly | |
| Reactome Gene Sets | 6 | R-HSA-1852241 | Organelle biogenesis and maintenance | |
| WikiPathways | 27 | WP3651 | Pathways affected in adenoid cystic carcinoma | |
| GO Biological Processes | 19 | GO:0006351 | DNA-templated transcription | 19_GO:0008152 metabolic process |
| GO Biological Processes | 19 | GO:0097659 | nucleic acid-templated transcription | 19_GO:0008152 metabolic process |
| Reactome Gene Sets | 6 | R-HSA-5620912 | Anchoring of the basal body to the plasma membrane | |
| GO Biological Processes | 19 | GO:0032774 | RNA biosynthetic process | 19_GO:0008152 metabolic process |
| GO Biological Processes | 19 | GO:0006338 | chromatin remodeling | 19_GO:0009987 cellular process |
| Reactome Gene Sets | 6 | R-HSA-2565942 | Regulation of PLK1 Activity at G2/M Transition | |
| GO Biological Processes | 19 | GO:0008380 | RNA splicing | 19_GO:0008152 metabolic process |
| GO Biological Processes | 19 | GO:0099111 | microtubule-based transport | 19_GO:0051179 localization |
| GO Biological Processes | 19 | GO:0010970 | transport along microtubule | 19_GO:0051179 localization |
| Reactome Gene Sets | 6 | R-HSA-380284 | Loss of proteins required for interphase microtubule organization from the centrosome | |
| Reactome Gene Sets | 6 | R-HSA-380259 | Loss of Nlp from mitotic centrosomes | |
| Reactome Gene Sets | 6 | R-HSA-8854518 | AURKA Activation by TPX2 | |
| Reactome Gene Sets | 6 | R-HSA-380270 | Recruitment of mitotic centrosome proteins and complexes | |
| Reactome Gene Sets | 6 | R-HSA-380287 | Centrosome maturation | |
| Reactome Gene Sets | 6 | R-HSA-68877 | Mitotic Prometaphase | |
| GO Biological Processes | 19 | GO:0030705 | cytoskeleton-dependent intracellular transport | 19_GO:0051179 localization |
| GO Biological Processes | 19 | GO:0000226 | microtubule cytoskeleton organization | 19_GO:0009987 cellular process |
| Reactome Gene Sets | 6 | R-HSA-380320 | Recruitment of NuMA to mitotic centrosomes | |
| GO Biological Processes | 19 | GO:0031023 | microtubule organizing center organization | 19_GO:0009987 cellular process |
| GO Biological Processes | 19 | GO:0006366 | transcription by RNA polymerase II | 19_GO:0008152 metabolic process |

**Supplementary figure 4 Mixed non-linear modeling of circadian genes**

**A**

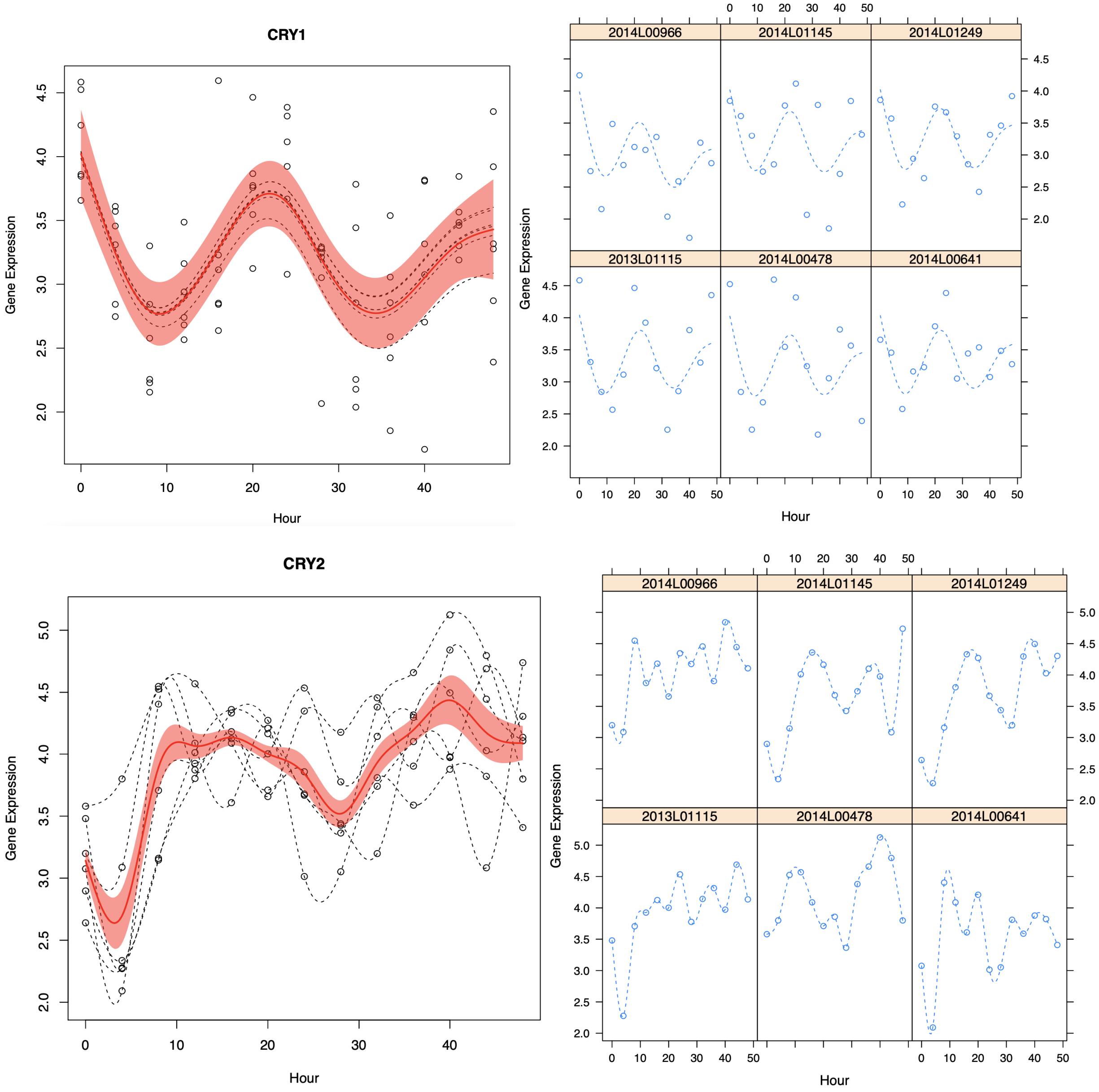
CRY1
Gene Expression
Hour
2014L00966
2014L01145
2014L01249
2013L01115
2014L00478
2014L00641
Gene Expression
Hour
CRY2
Gene Expression
Hour
2014L00966
2014L01145
2014L01249
2013L01115
2014L00478
2014L00641
Gene Expression
Hour

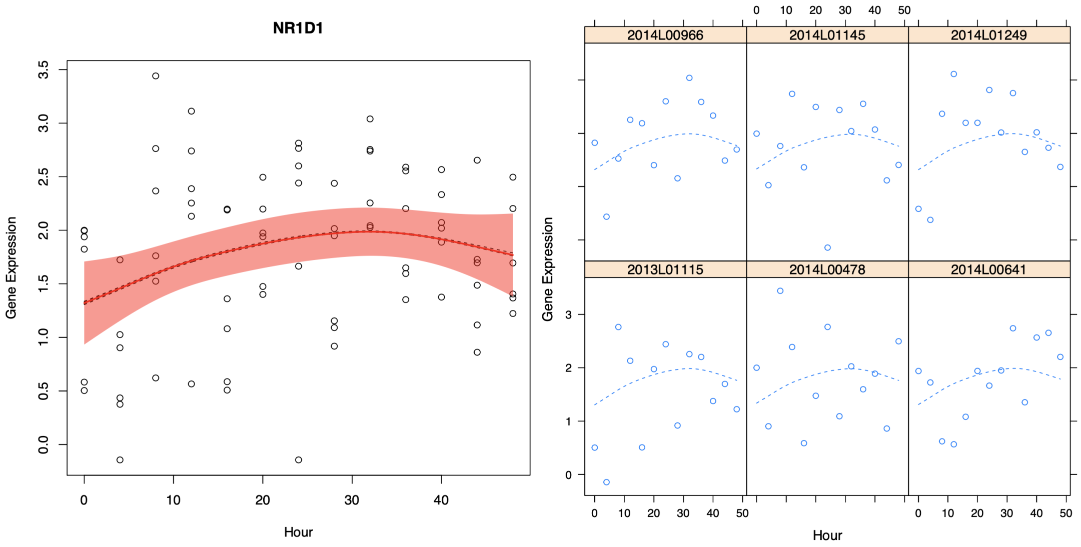
NR1D1
Gene Expression
Hour
2014L00966
2014L01145
2014L01249
2013L01115
2014L00478
2014L00641
Gene Expression
Hour

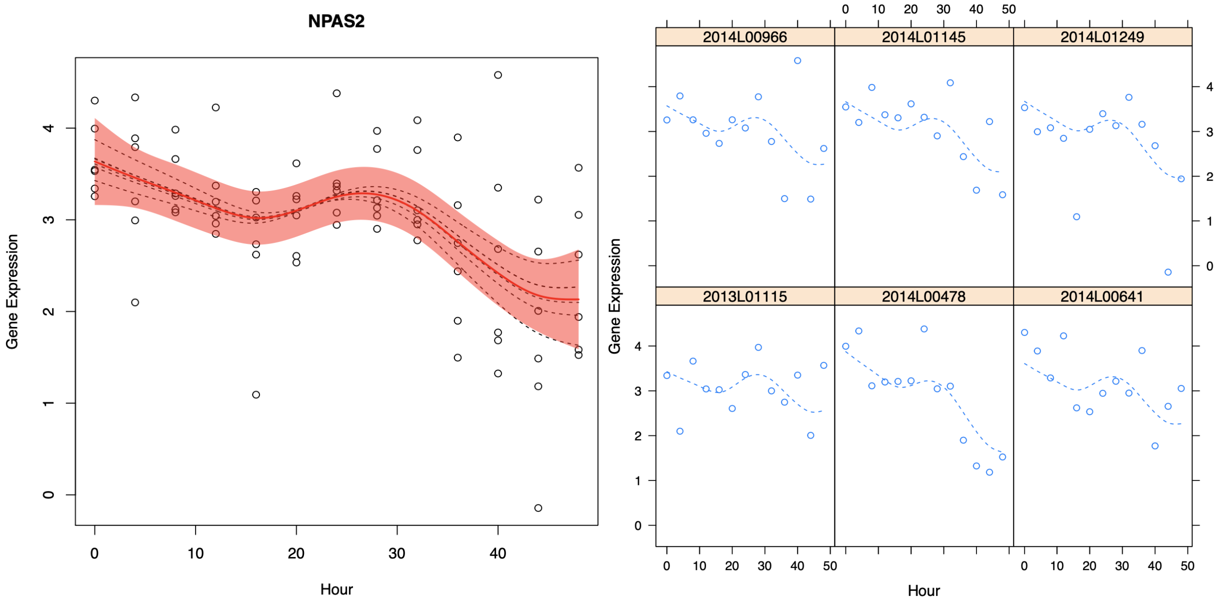
NPAS2
Gene Expression
Hour
2014L00966
2014L01145
2014L01249
2013L01115
2014L00478
2014L00641
Gene Expression
Hour

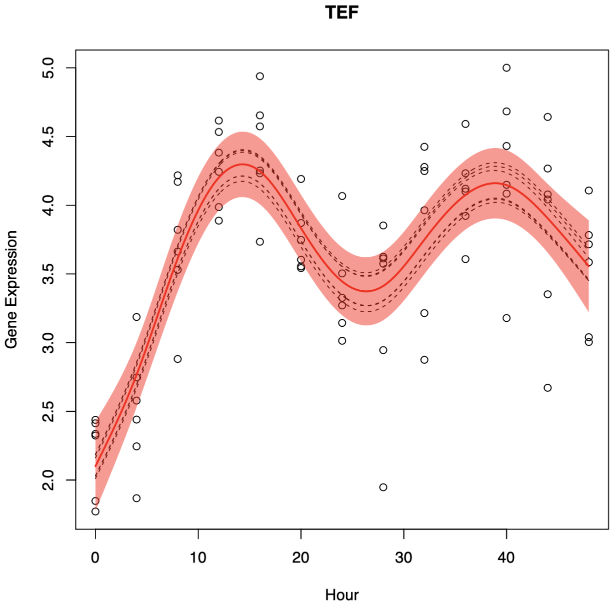
TEF
Gene Expression
Hour

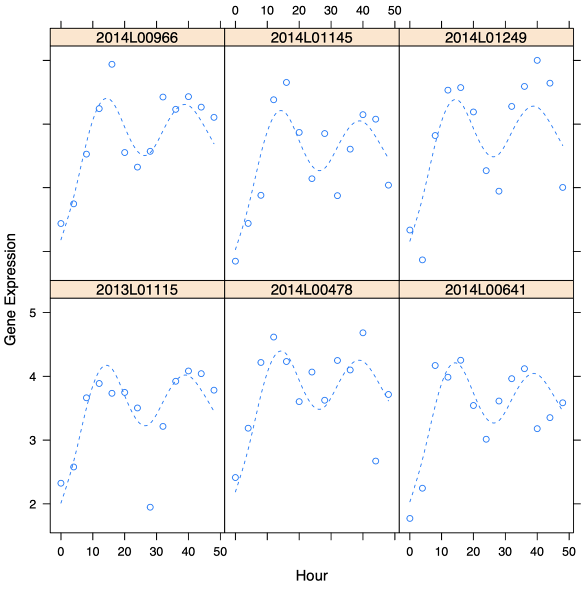
2014L00966
2014L01145
2014L01249
2013L01115
2014L00478
2014L00641
Gene Expression
Hour

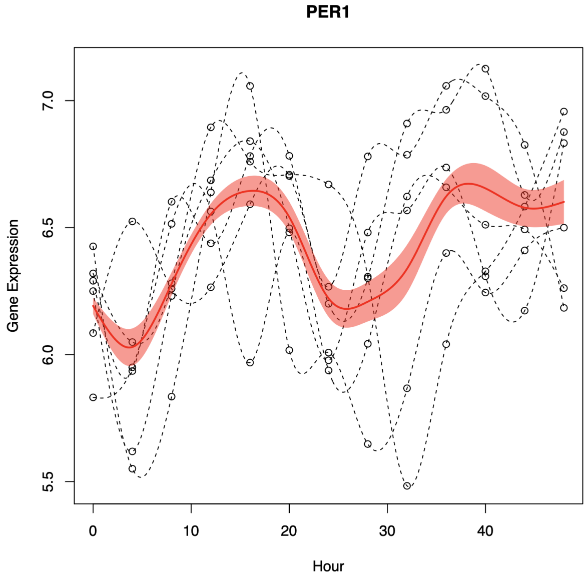
PER1
Gene Expression
Hour

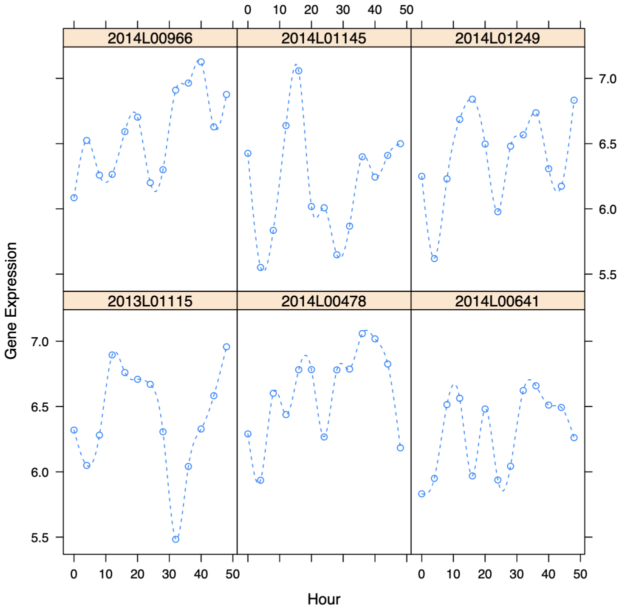
2014L00966
2014L01145
2014L01249
2013L01115
2014L00478
2014L00641
Gene Expression
Hour

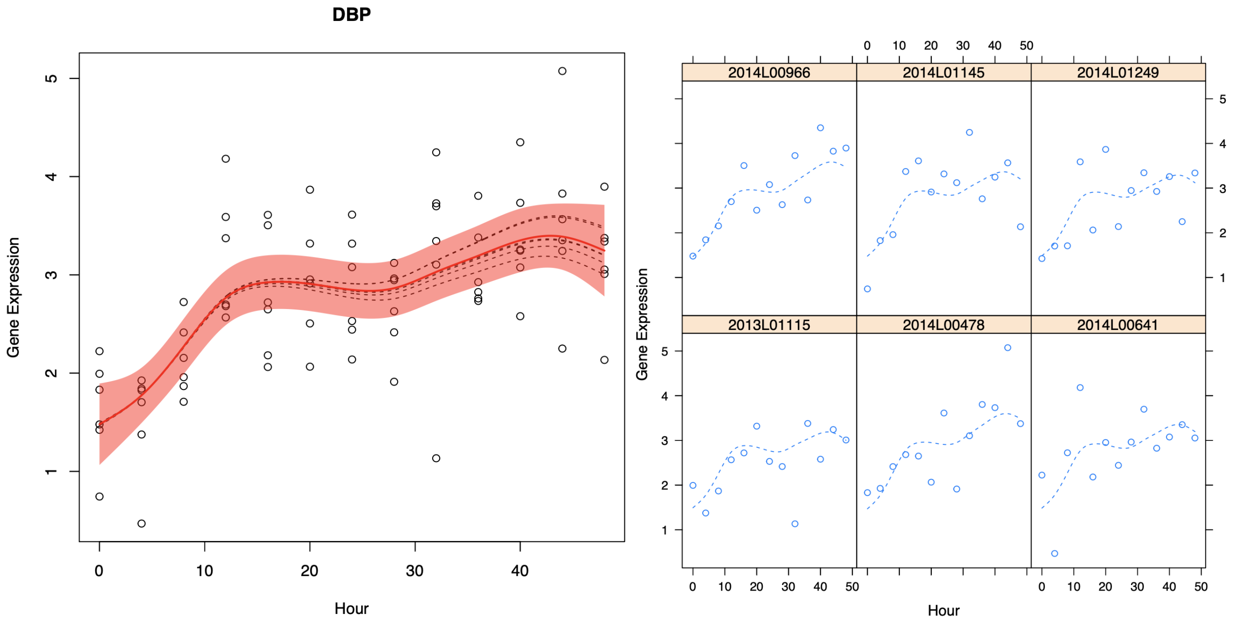
DBP
Gene Expression
Hour
2014L00966
2014L01145
2014L01249
2013L01115
2014L00478
2014L00641

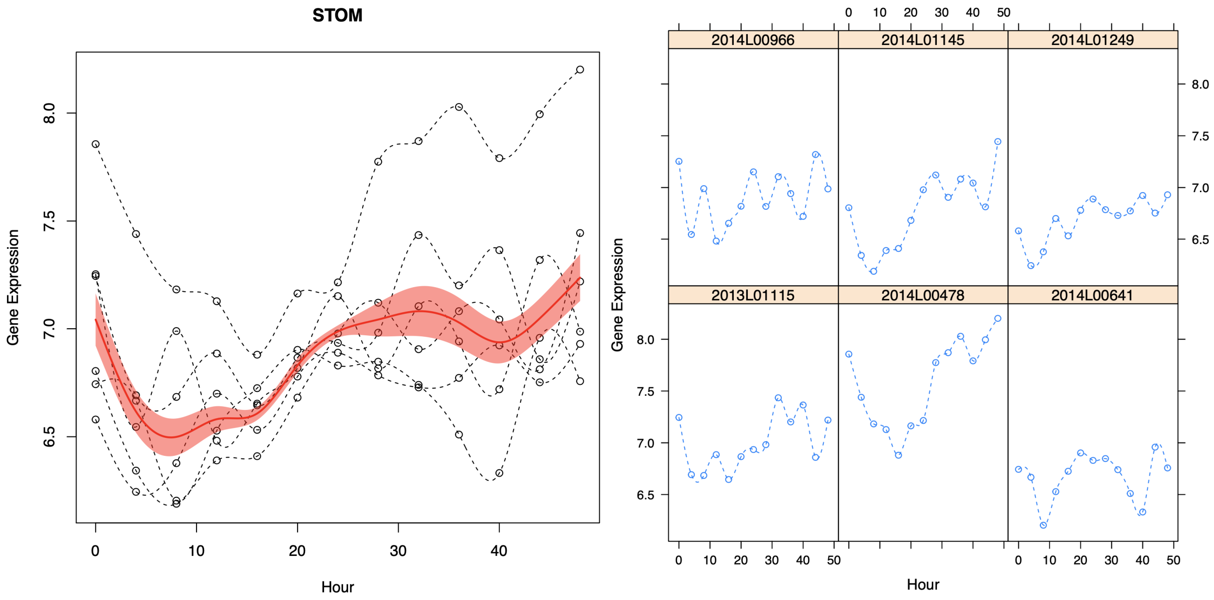
STOM
Gene Expression
Hour
2014L00966
2014L01145
2014L01249
2013L01115
2014L00478
2014L00641

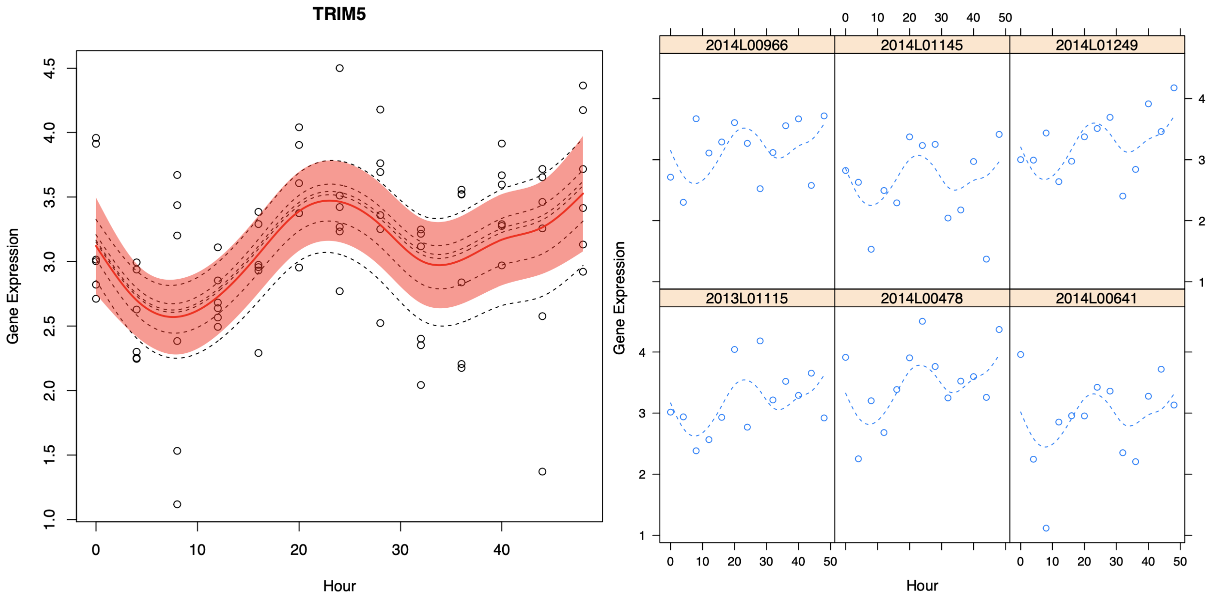
TRIM5
Gene Expression
Hour
2014L00966
2014L01145
2014L01249
2013L01115
2014L00478
2014L00641

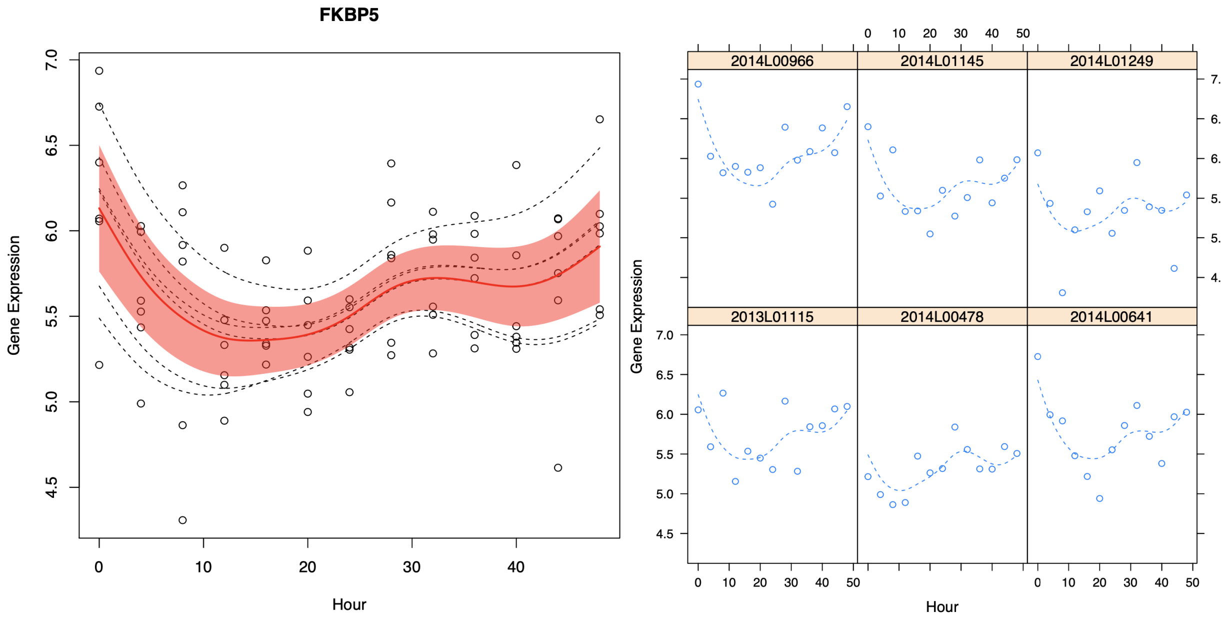
FKBP5
Gene Expression
Hour
2014L00966
2014L01145
2014L01249
2013L01115
2014L00478
2014L00641

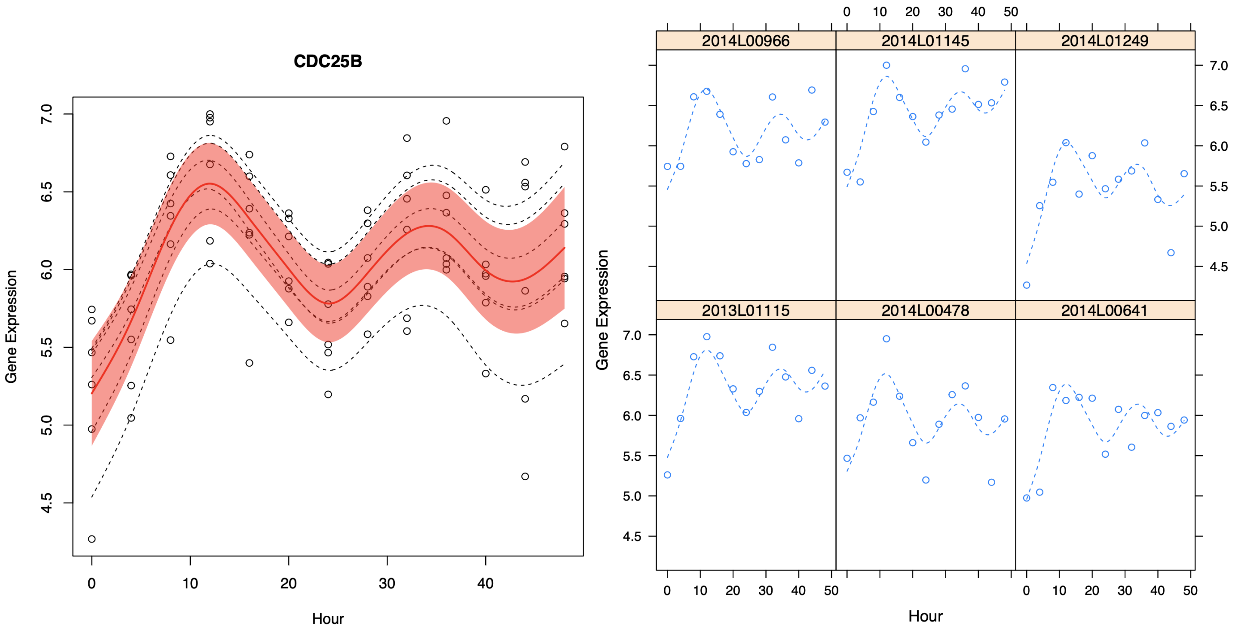
CDC25B
Gene Expression
Hour
2014L00966
2014L01145
2014L01249
2013L01115
2014L00478
2014L00641
Gene Expression
Hour

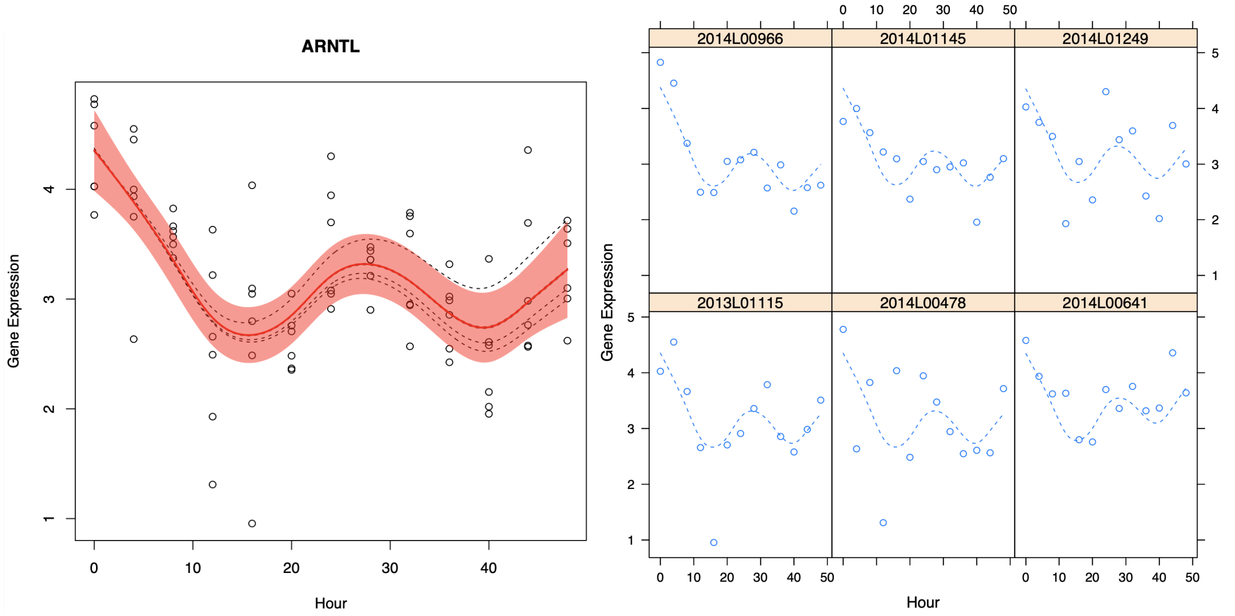
ARNTL
Gene Expression
Hour
2014L00966
2014L01145
2014L01249
2013L01115
2014L00478
2014L00641
Gene Expression
Hour

**NFIL3**

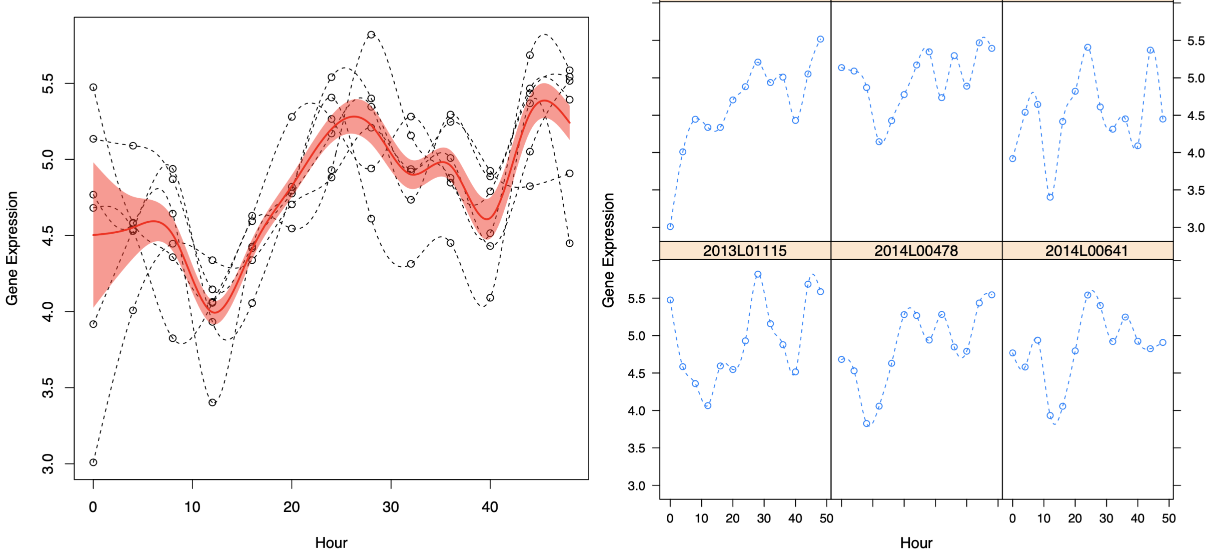
Gene Expression
Hour
2014L00966
2014L01145
2014L01249
2013L01115
2014L00478
2014L00641

**PER3**

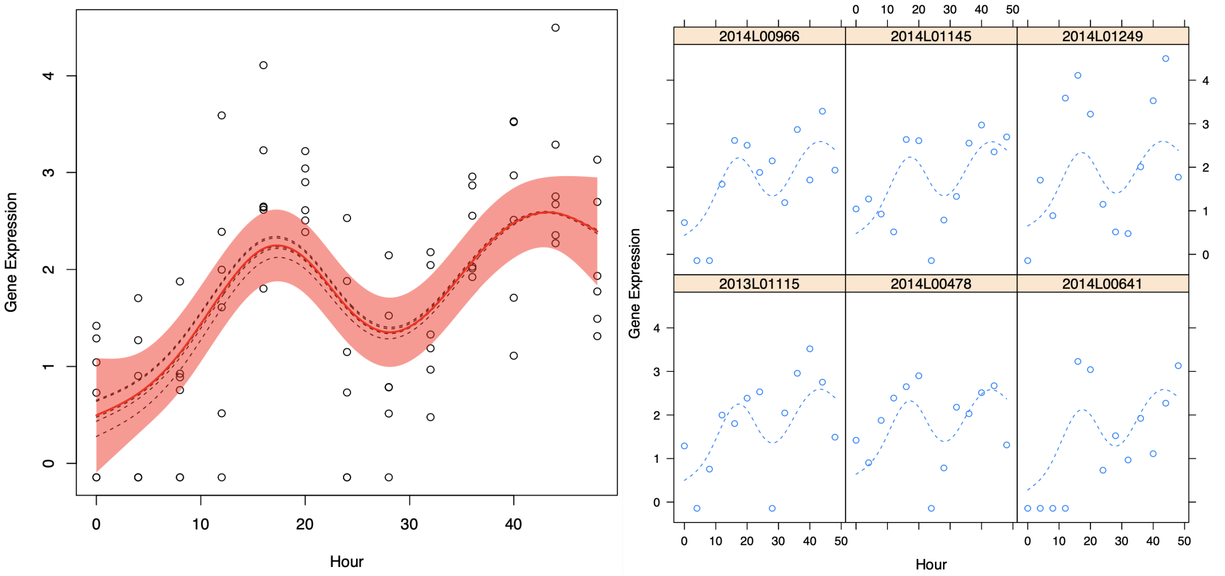
Gene Expression
Hour
2014L00966
2014L01145
2014L01249
2013L01115
2014L00478
2014L00641

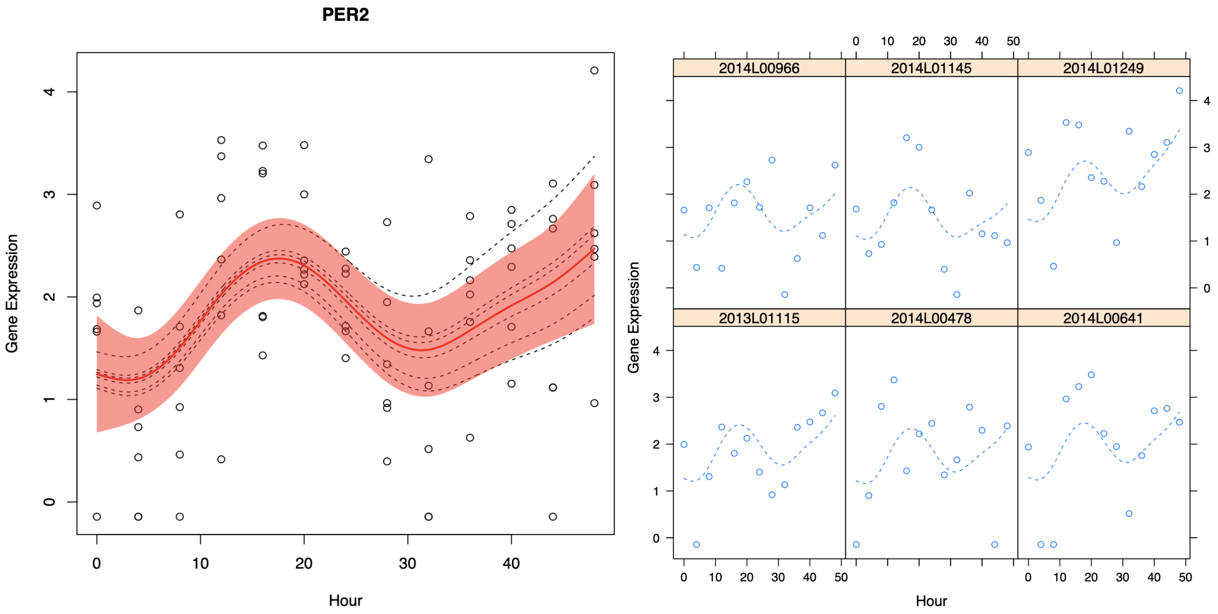


**Supplementary figure 4 description:** A, Smoothing-splines mixed-effect models of circadian gene expression across previously reported circadian genes in skin. Red line indicates the average fitted model across cell lines, with the red area representing a 95% confidence interval.

**Supplemental figure 5** Known interactions between circadian genes present identified in this dataset

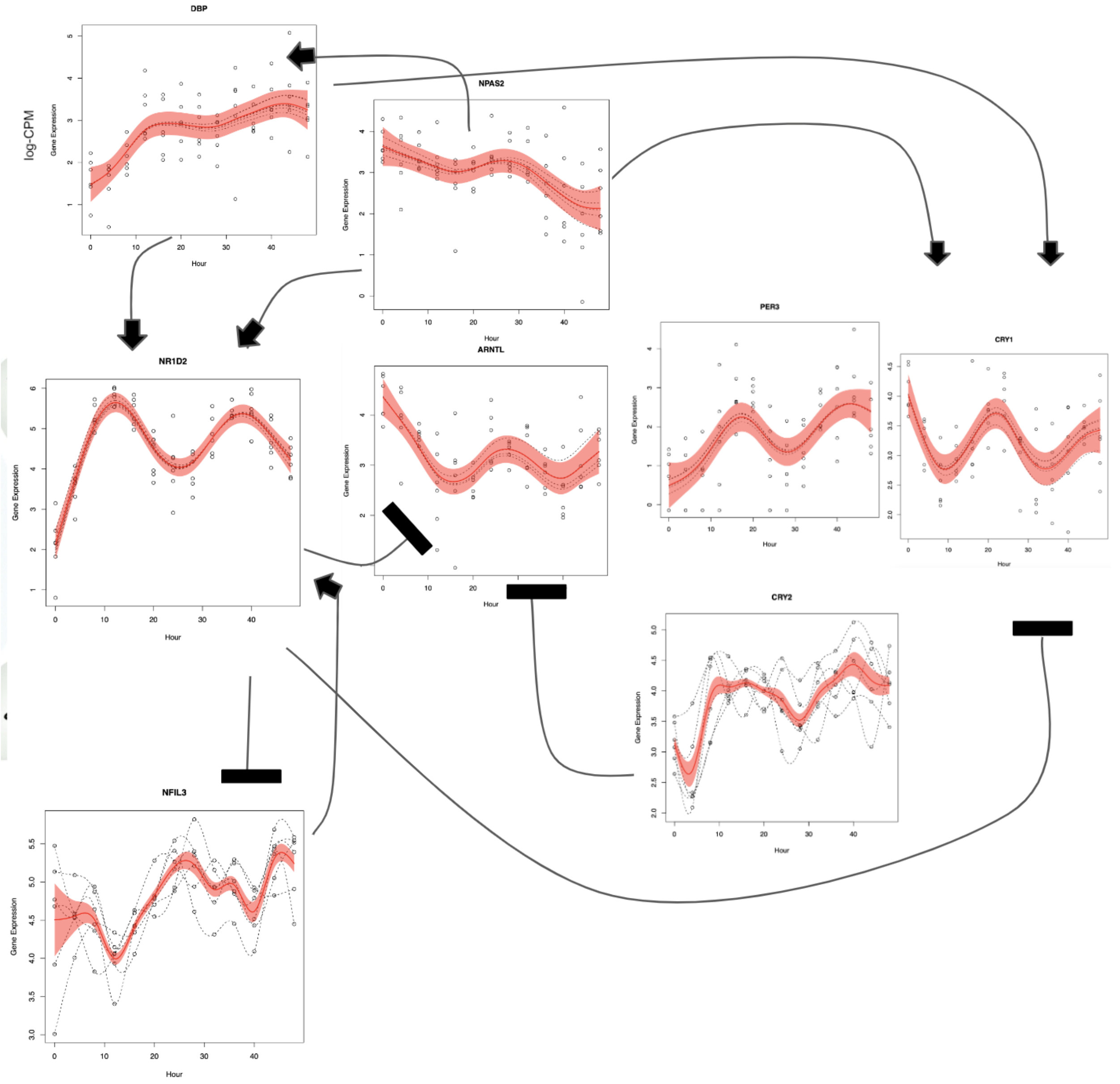


**Supplementary figure 5 description**: Known gene expression relationships of core circadian genes. Arrows indicate a gene inducing in the expression of another gene. The black bar indicates a gene repressing the expression of another gene.

**Supplementary table 2:** Results for circadian analysis tools ARSER, JTK, LS and metacycle

ARSER (Full results in MetaCycle Results folder)

| CycID | meta3d_Pvalue | meta3d_BH.Q | meta3d_Period | meta3d_Phase | meta3d_Base | meta3d_AMP | meta3d_rAMP |
|---|---|---|---|---|---|---|---|
| NR1D2 | 1.213429e-11 | 1.335258e-07 | 24.77884 | 12.3604684 | 4.587153 | 1.1940252 | 0.26131517 |
| TEF | 1.715038e-09 | 9.436137e-06 | 23.77328 | 13.5969101 | 3.699635 | 0.8258222 | 0.22327689 |
| CRY1 | 2.518402e-07 | 9.237500e-04 | 23.53269 | 21.9876554 | 3.224590 | 0.7364189 | 0.22766198 |
| TECPR1 | 8.314857e-07 | 2.287417e-03 | 23.10611 | 13.2858023 | 4.907502 | 0.4514711 | 0.09187609 |
| CDC25B | 2.614718e-06 | 5.754472e-03 | 21.75986 | 12.1292808 | 6.138671 | 0.4665738 | 0.07624382 |
| HNRNPU | 4.643564e-06 | 8.516295e-03 | 23.89180 | 22.2422183 | 8.048284 | 0.1775919 | 0.02208631 |
| COG8 | 8.449688e-06 | 1.109890e-02 | 23.42245 | 2.8576798 | 1.871940 | 0.9216777 | 0.51115138 |
| YBEY | 9.077615e-06 | 1.109890e-02 | 23.81287 | 20.9649802 | 3.579168 | 0.4444663 | 0.12770498 |
| ANGPTL2 | 8.758998e-06 | 1.109890e-02 | 21.93092 | 18.0668207 | 6.473876 | 0.4057508 | 0.06259919 |
| PER3 | 1.238981e-05 | 1.239431e-02 | 24.56212 | 16.2925841 | 1.839917 | 1.2330494 | 0.65617035 |
| KIAA1244 | 1.221822e-05 | 1.239431e-02 | 23.66213 | 12.9676442 | 2.154037 | 0.7363603 | 0.38380775 |
| RGS4 | 1.420238e-05 | 1.302358e-02 | 21.36553 | 20.9752510 | 3.487784 | 0.6029858 | 0.18236976 |
| CCDC22 | 1.565106e-05 | 1.324802e-02 | 24.96972 | 22.8001950 | 3.048887 | 0.6679597 | 0.21892731 |
| RASSF4 | 2.297644e-05 | 1.805948e-02 | 22.03420 | 10.8342720 | 3.589962 | 0.5734768 | 0.16243627 |
| LRP12 | 2.669229e-05 | 1.958146e-02 | 22.66092 | 21.0085194 | 5.286893 | 0.3908946 | 0.07430898 |
| MRPS25 | 2.907814e-05 | 1.999849e-02 | 23.27449 | 1.8402432 | 6.042704 | 0.2289662 | 0.03786866 |
| TRIT1 | 5.205274e-05 | 3.017278e-02 | 24.63071 | 22.7026574 | 3.092441 | 0.7523076 | 0.24757035 |
| TRAF3IP3 | 5.483965e-05 | 3.017278e-02 | 24.18468 | 8.6541727 | 1.986866 | 0.5518161 | 0.28633139 |
| NEAT1 | 5.003333e-05 | 3.017278e-02 | 25.64616 | 9.2529887 | 8.313655 | 0.2999881 | 0.03629947 |
| CPM | 5.357480e-05 | 3.017278e-02 | 21.85298 | 9.7703230 | 7.692930 | 0.3817325 | 0.04931089 |
| DFFB | 6.756078e-05 | 3.128110e-02 | 24.20344 | 19.7974604 | 2.767652 | 0.6863324 | 0.25005403 |

JTK

| CycID | meta3d_Pvalue | meta3d_BH.Q | meta3d_Period | meta3d_Phase | meta3d_Base | meta3d_AMP | meta3d_rAMP |
|---|---|---|---|---|---|---|---|
| NR1D2 | 9.595880e-12 | 1.055931e-07 | 27.68920 | 1.334773e+01 | 4.4620012 | 1.18082732 | 0.265828928 |
| TEF | 2.655185e-05 | 1.460883e-01 | 25.84020 | 1.566192e+01 | 3.6157010 | 0.77420363 | 0.213709662 |
| LOC102723897 | 9.999611e-01 | 1.000000e+00 | 20.00000 | 1.400000e+01 | 2.7681907 | 0.44691374 | 0.161446154 |
| LINC01128 | 9.570960e-01 | 1.000000e+00 | 23.04465 | 6.284749e+00 | 2.5484198 | 0.50800380 | 0.201301191 |
| NOC2L | 7.292068e-01 | 1.000000e+00 | 26.89478 | 1.041780e+01 | 1.9934585 | 0.56383520 | 0.290242126 |

LS

| CycID | meta3d_Pvalue | meta3d_BH.Q | meta3d_Period | meta3d_Phase | meta3d_Base | meta3d_AMP | meta3d_rAMP |
|---|---|---|---|---|---|---|---|
| NR1D2 | 0.01633951 | 1 | 25.64740 | 12.87639324 | 4.5440208 | 1.2277886 | 0.27153134 |
| TEF | 0.05023642 | 1 | 24.95835 | 15.27871444 | 3.6363365 | 0.8011303 | 0.22052088 |
| PER3 | 0.12094754 | 1 | 25.61226 | 17.01255748 | 1.8317137 | 1.2665310 | 0.67639039 |
| ZNF772 | 0.20681331 | 1 | 25.37394 | 1.94345724 | 3.2468177 | 0.6224801 | 0.19051836 |
| COG8 | 0.21600155 | 1 | 23.46244 | 2.96438099 | 1.8183856 | 0.9070828 | 0.51457672 |
| CRY1 | 0.23380311 | 1 | 23.53425 | 21.96794674 | 3.2583237 | 0.7522083 | 0.23038240 |

### Supplemental Figure 6 Quality Control for ATAC-seq data

**A**

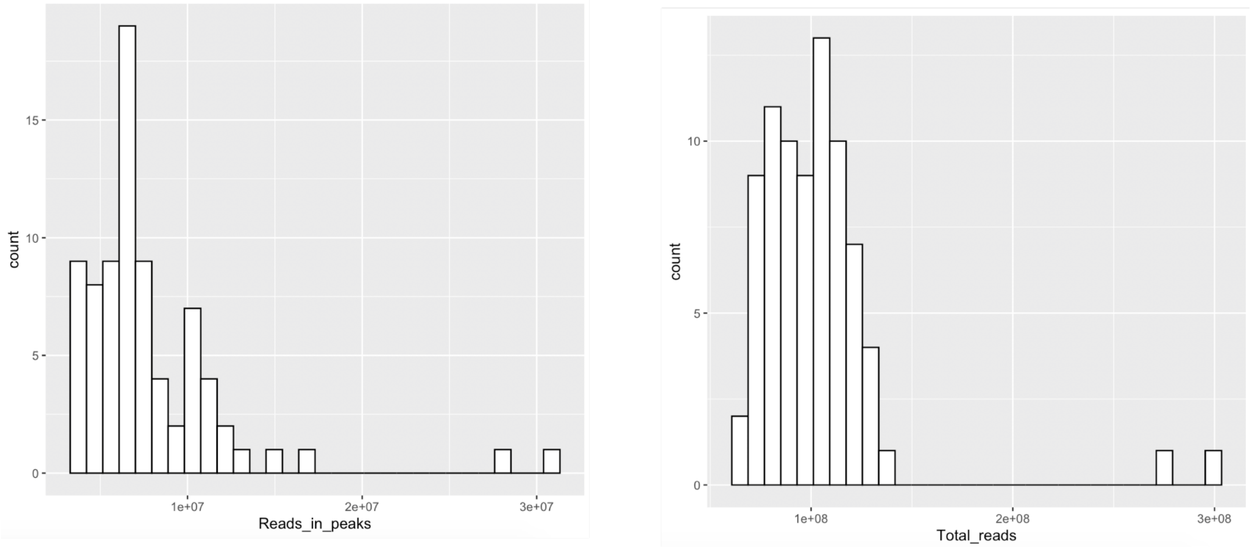


**B**

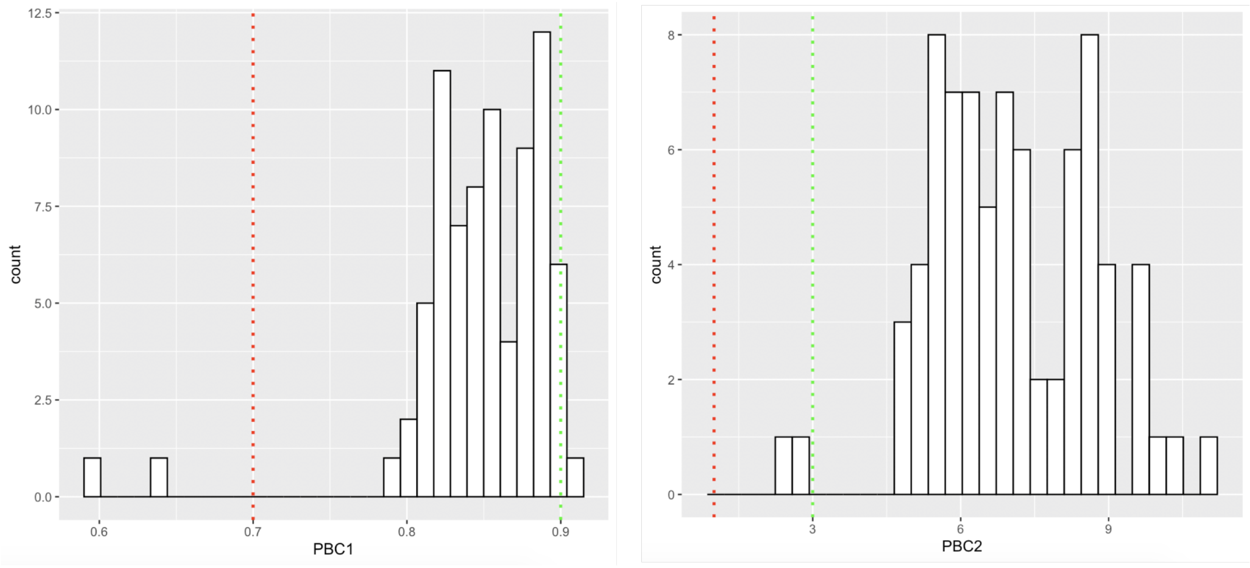

**C**

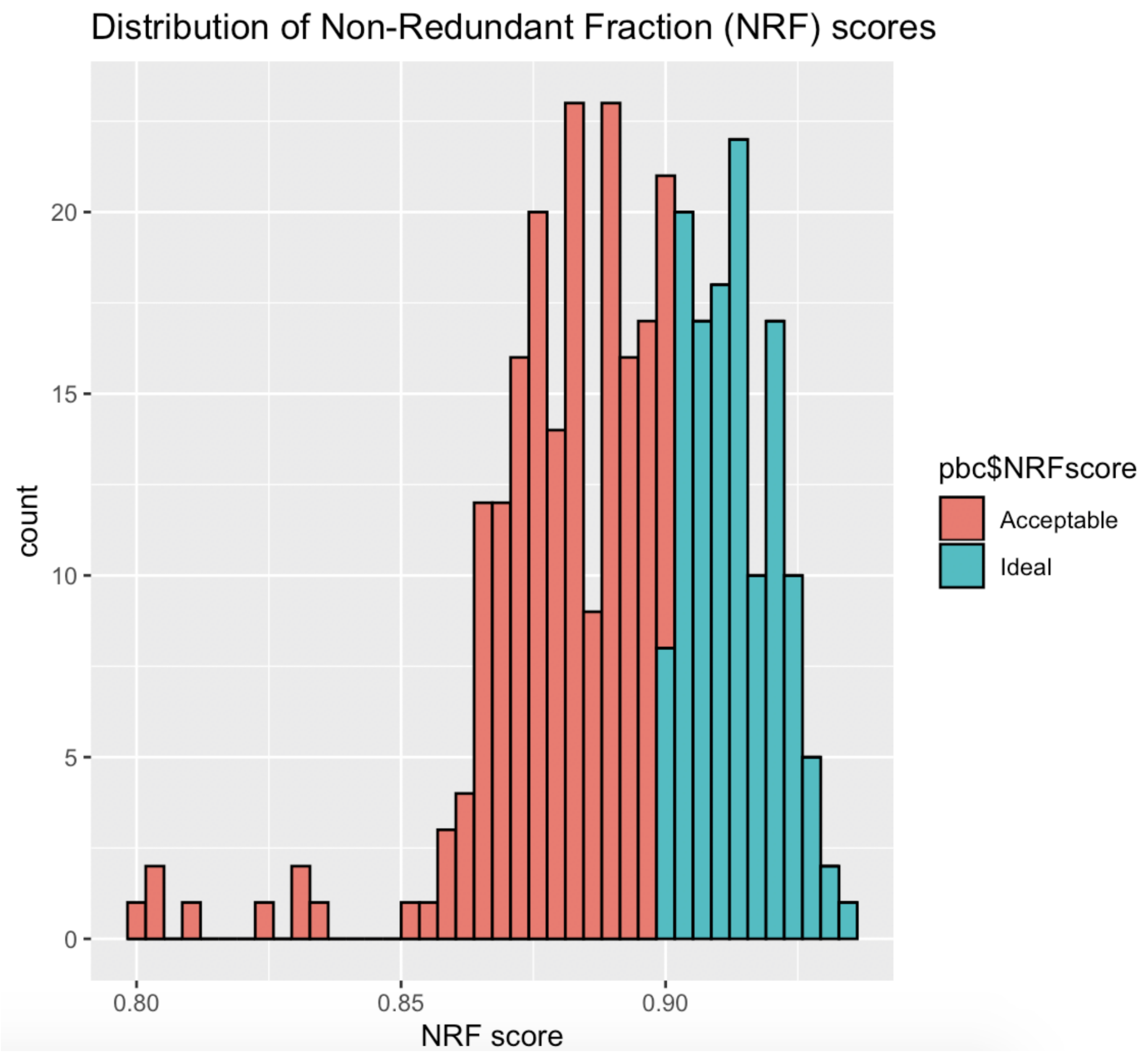


**Supplementary Figure 6 Description:** Quality control metrics, using the thresholds provided by the standard ENCODE Pipeline for the identification of open chromatin regions (OCRs) of the genome. A shows distribution of all reads present in the peaks identified in the ATAC-seq dataset, as well as the distribution of all the reads by sample. B shows library complexity metrics of PCR Bottlenecking Coefficients (PBC), measured as PBC1 = [# of positions with exactly 1 read mapped] / [# of positions with 1 or more reads mapped ] and PBC2 = [# of positions with exactly 1 read mapped] / [# of positions with 2 reads mapped ]. Red cutoff color indicates severe PCR bottlenecking threshold (<0.7 for PBC1, <1 for PBC2), green cutoff corresponds to no PCR bottleneck (>0.9 for PBC1, >3 for PBC2). C shows the distribution of Non-redundant Fraction(NRF) scores. Values within 0.7 and 0.9

are considered acceptable. The outlying samples across QC were from the 2014L00966 cell line, time points 20 and 0.

**Supplemental Figure 7 Eigengene values for ATAC-seq modules obtained from WGCNA**

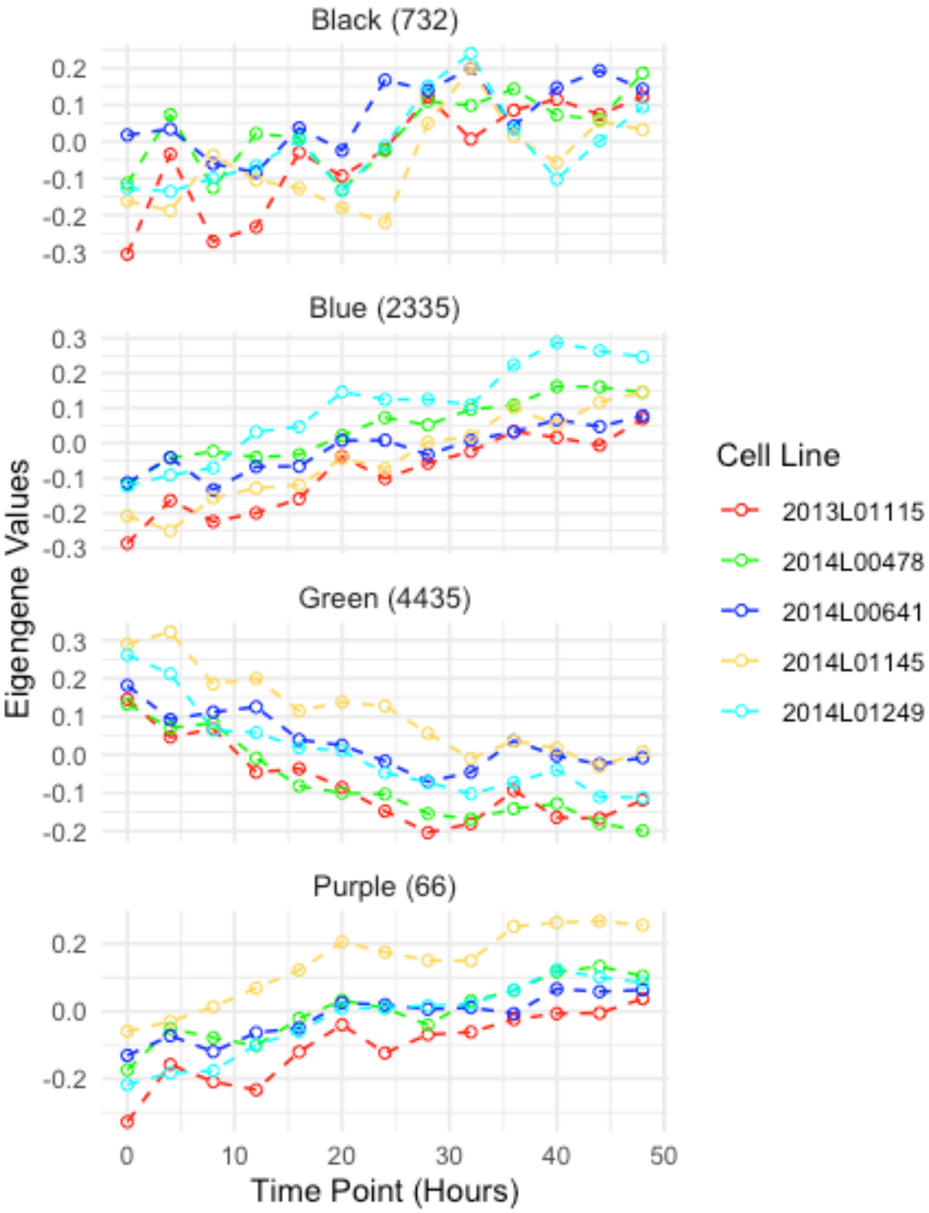


**Supplemental Figure 7 Description:** Eigengene modules from WGCNA of the longitudinal chromatin accessibility patterns of ATAC-seq data collected every 4 hours for a 48 hour period. Each color represents a fibroblast cell culture from a different individual. The number of chromatin accessible regions assigned per module is indicated next to the module name.

**Supplementary Figure 8 Genomic annotation of the consensus peak regions and selected time significant regions**

A. All consensus regions

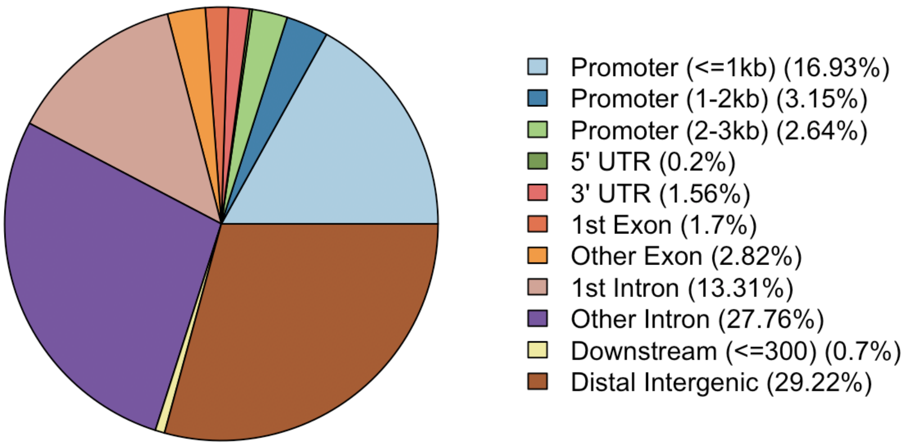


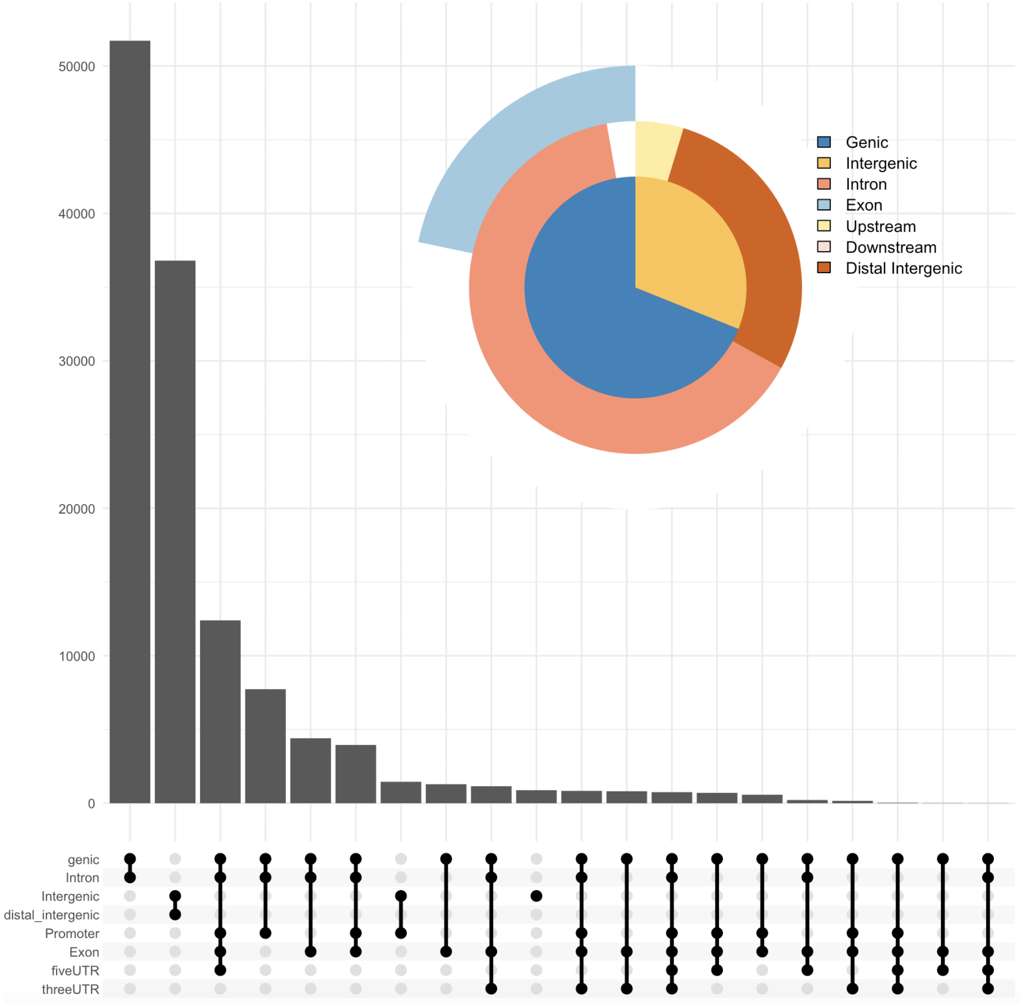

B. Time significant regions

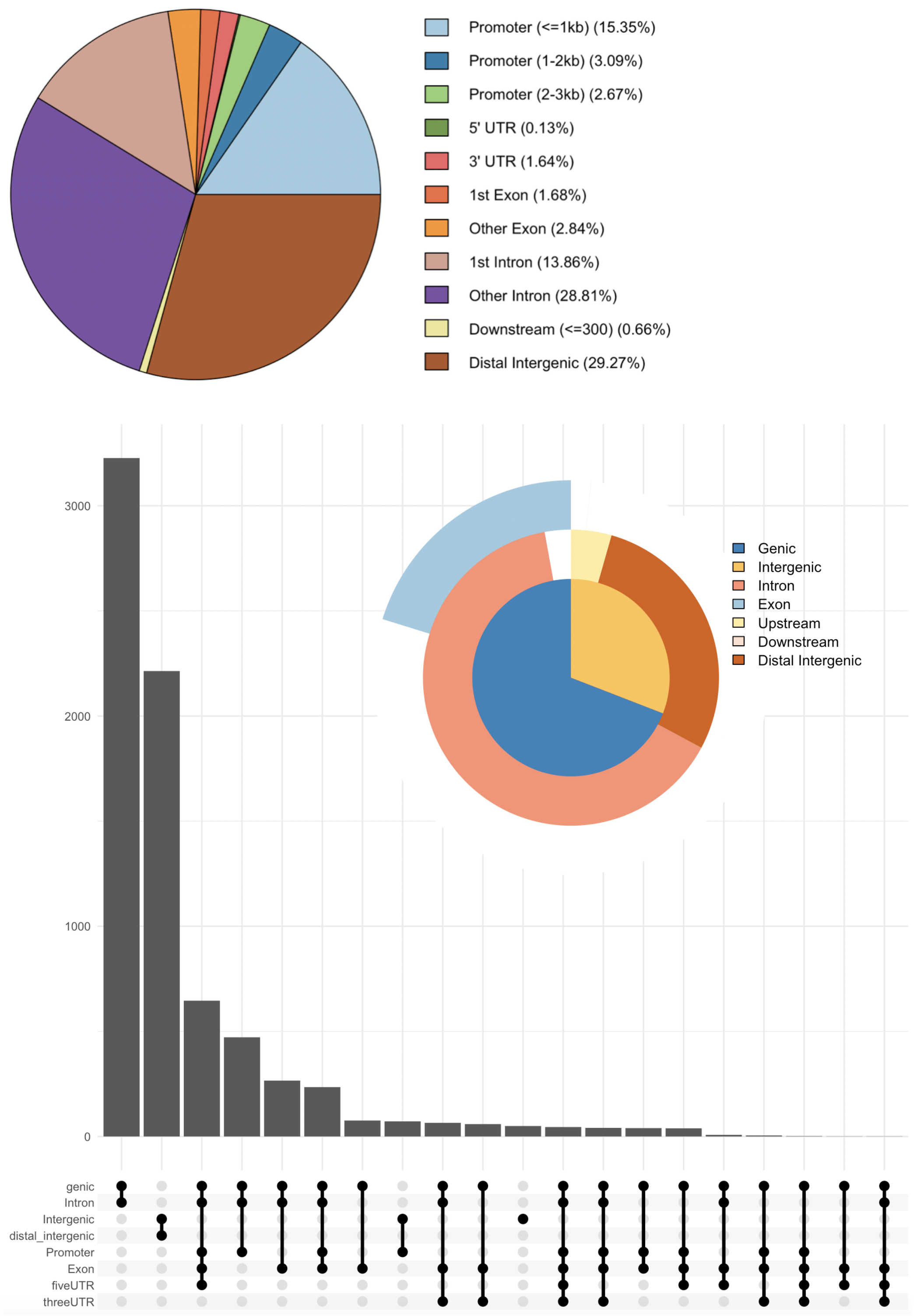


**Supplementary Figure 8 description:** Chipseeker annotations for peak regions. A. Genomic annotations for all n=126,057 consensus peak regions. B. Genomic annotations for the n=7,568 peaks that had a significant change over time in their accessibility.

**Supplementary figure 9 Schematic of synchronization and collection times**

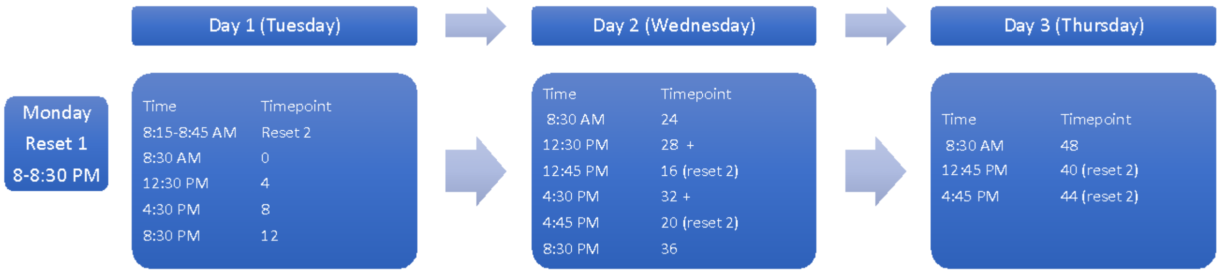


**Supplementary Figure 9 description:** Collection scheme for both RNA-seq and ATA-seq fibroblast cell culture samples. Cells were reset 12 hours before the first collection. In order to collect RNA or cells every 4 hours for 48 hours, cells were split into two batches, which were reset 12 hours apart.
