## Supplementary material for "Fibroblasts as an in vitro model of circadian genetic and genomic studies: A temporal analysis": TSS QC Summary.pdf

2013L01115-TP-0 **4.6**

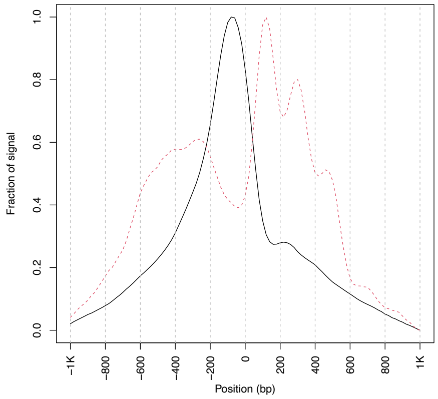


2013L01115-TP-4. **3.1**

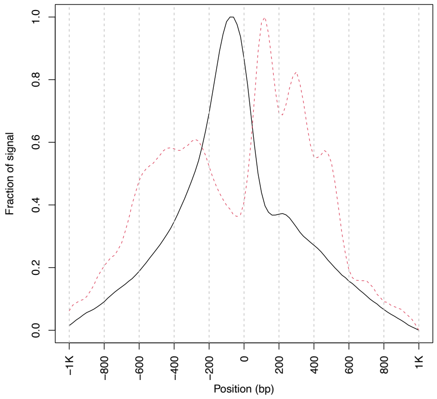


2013L01115-TP-8 **4.3**

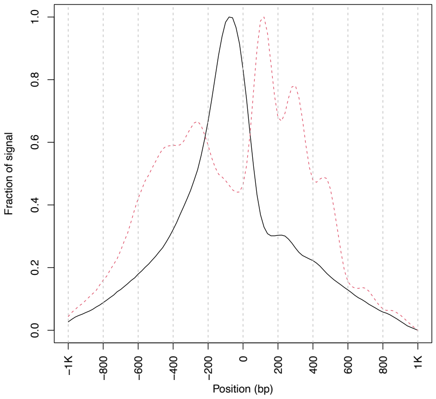


2013L01115-TP-12 **3.9**

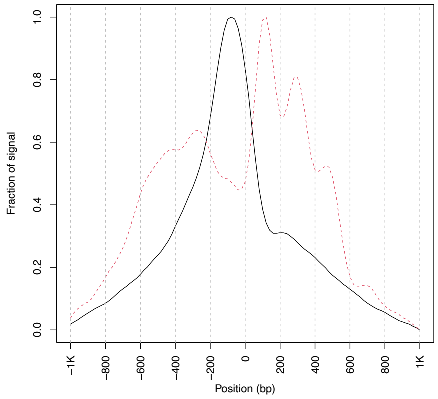


2013L01115-TP-16 **3.2**

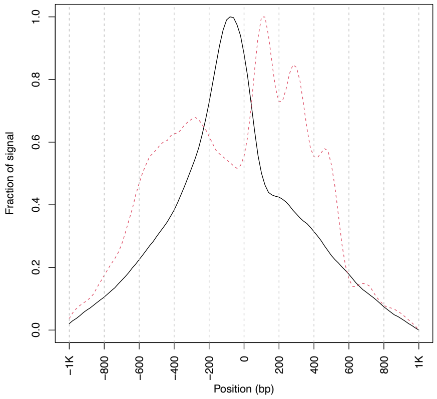


2013L01115-TP-20 **3.4**

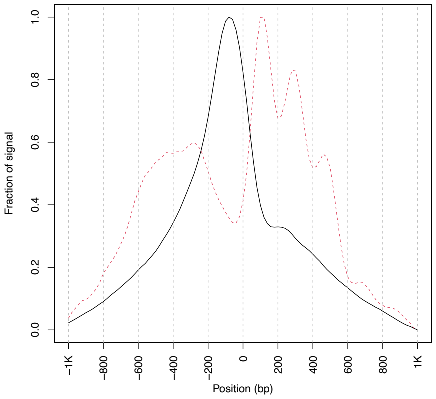


2013L01115-TP-24 **2.8**

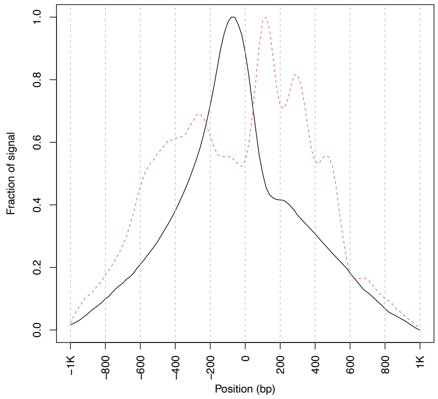


2013L01115-TP-28. **2.6**

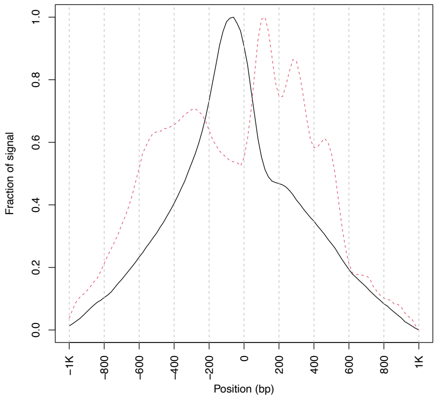


2013L01115-TP-32. **3.5**

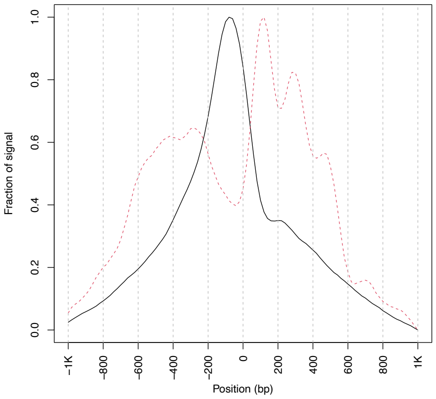

2013L01115-TP-36 **3.2**

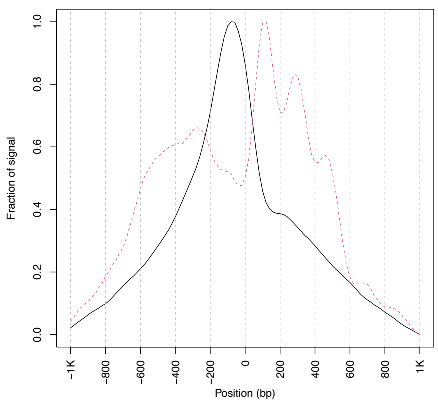


2013L01115-TP-40 **2.6**

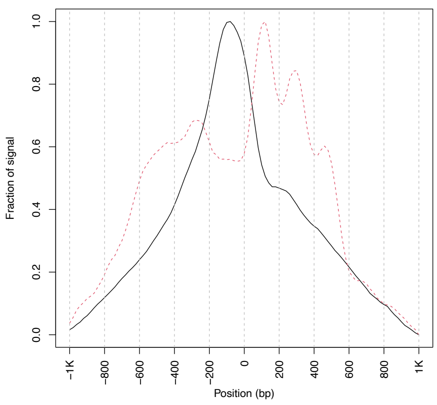


2013L01115-TP-44 **2.6**

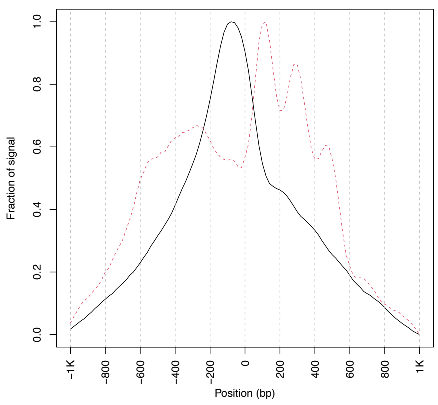


2013L01115-TP-48 **2.8**

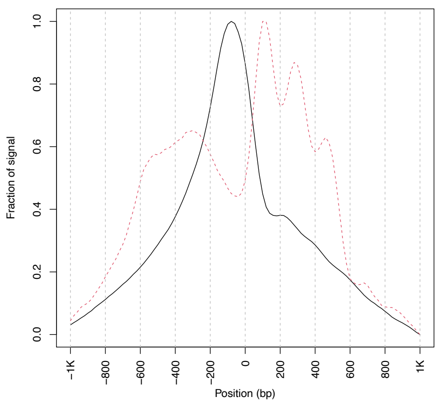

2014L00478-TP-0 **3.5**

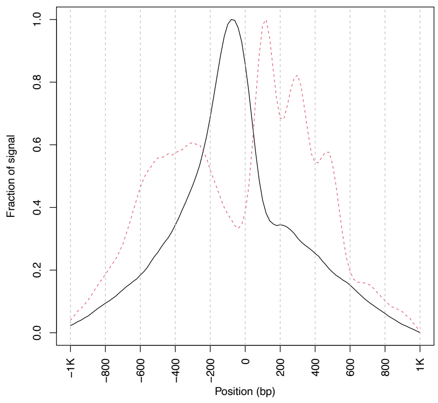


2014L00478-TP-4 **3.0**

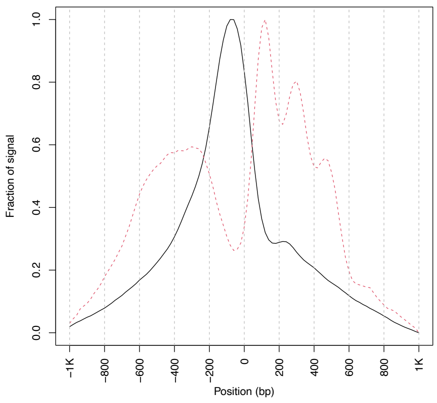


2014L00478-TP-8 **3.8**

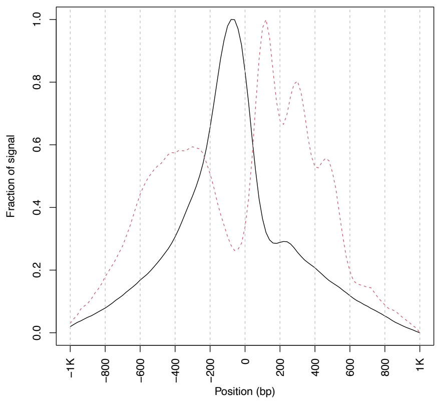


2014L00478-TP-12 **3.3**

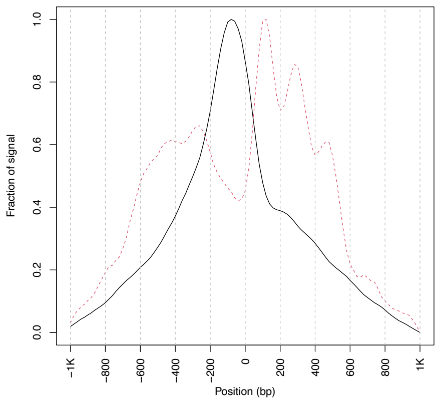


2014L00478-TP-16 **2.8**

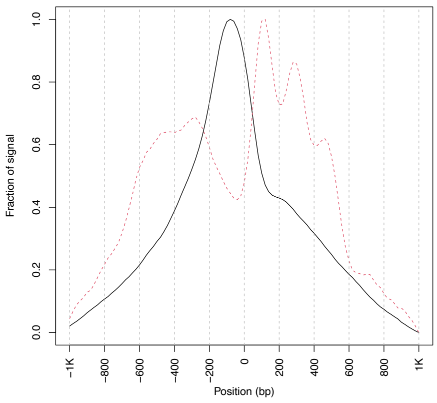


2014L00478-TP-20 **3.7**

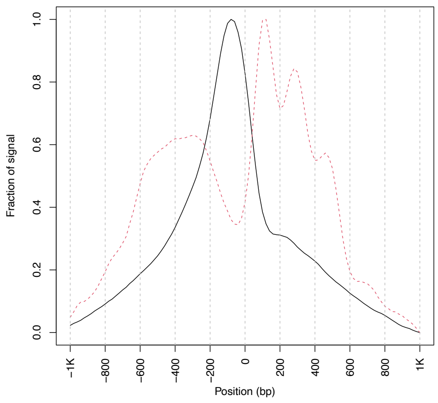


2014L00478-TP-24 **3.3**

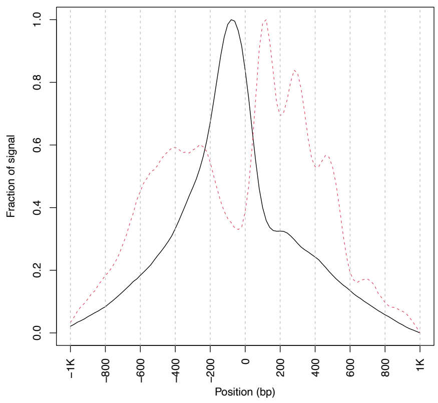


2014L00478-TP-28 **2.7**

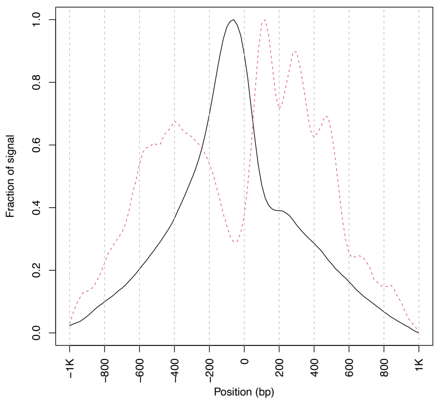


2014L00478-TP-32 **3.0**

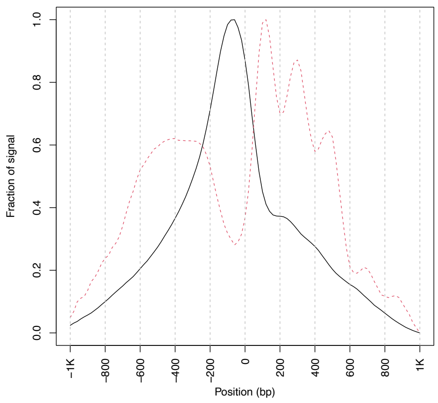

2014L00478-TP-36 **3.0**

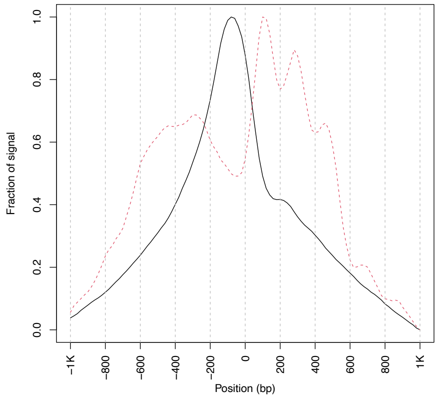


2014L00478-TP-40 **3.4**

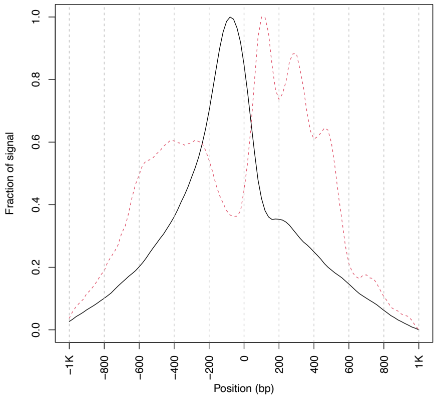


2014L00478-TP-44 **3.1**

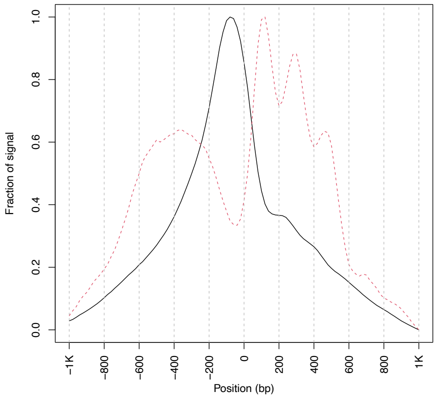


2014L00478-TP-48 **2.7**

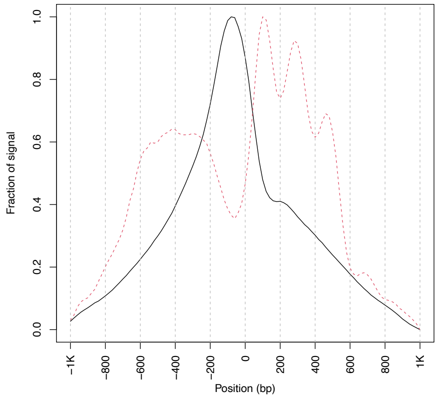

2014L00641-TP-0 **3.1**

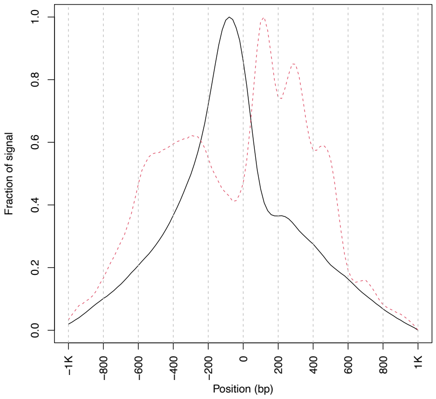


2014L00641-TP-4 **3.1**

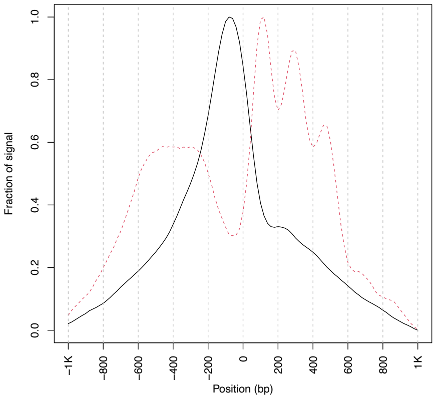


2014L00641-TP-8 **3.6**

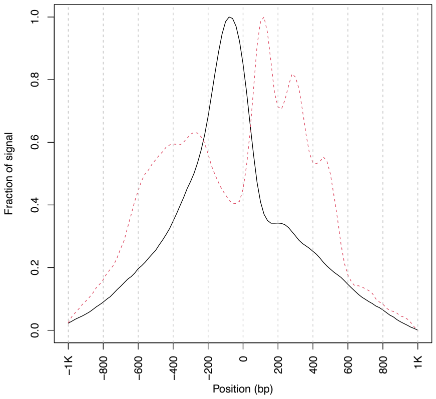


2014L00641-TP-12 **3.2**

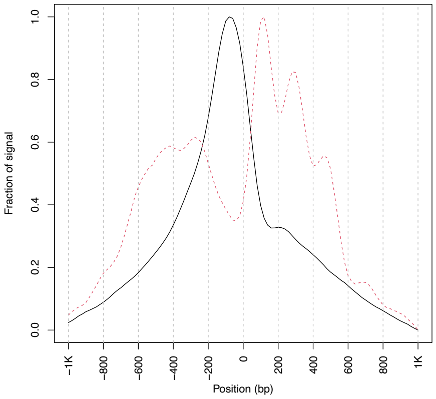


2014L00641-TP-16 **3.0**

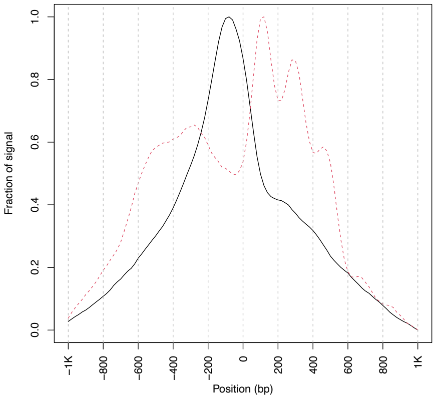


2014L00641-TP-20 **3.2**

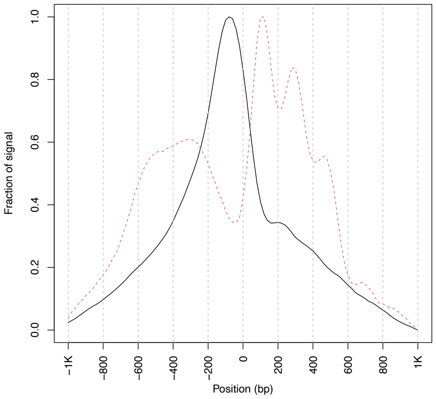


2014L00641-TP-24 **3.0**

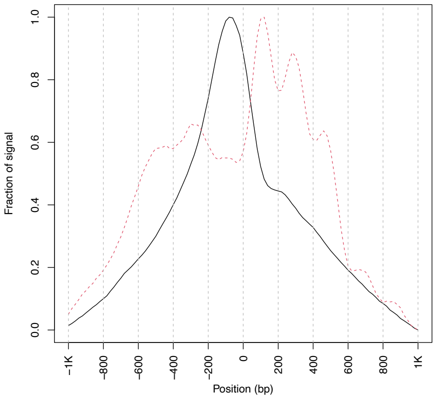


2014L00641-TP-28 **2.8**

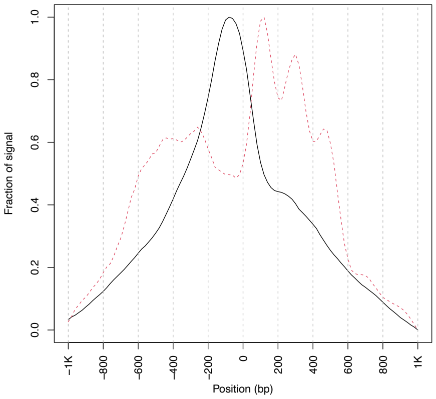


2014L00641-TP-32 **2.4**

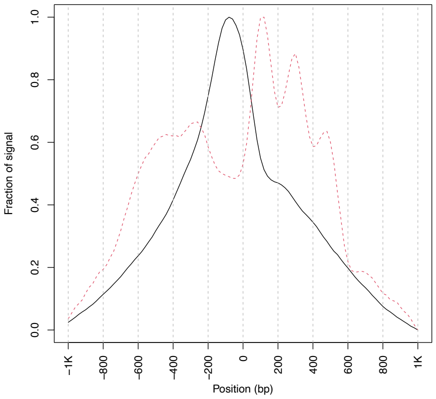

2014L00641-TP-36 **3.2**

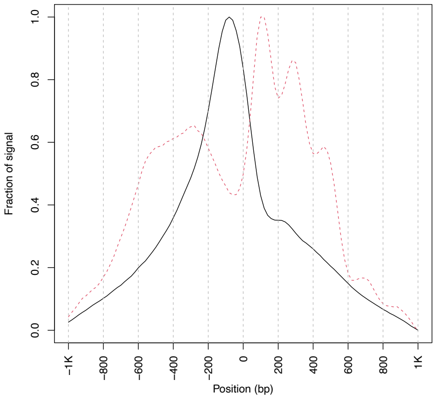


2014L00641-TP-40 **3.0**

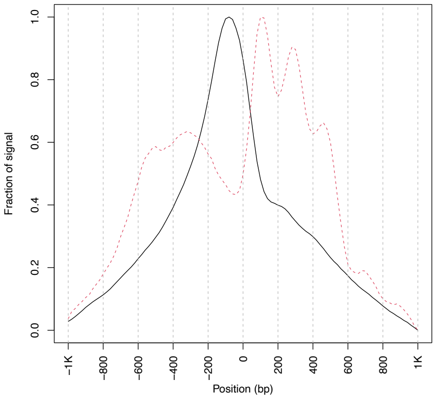


2014L00641-TP-44 **2.6**

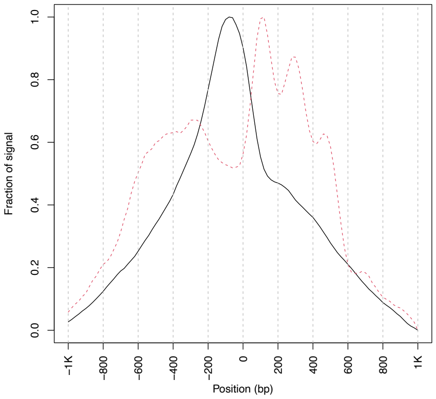


2014L00641-TP-48 **3.1**

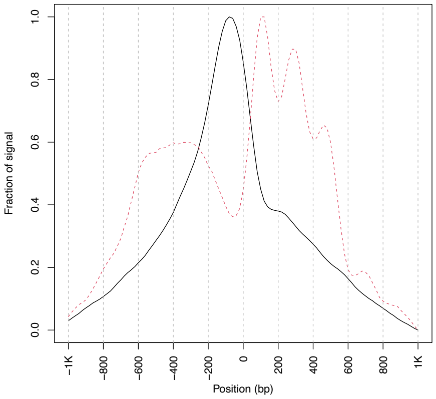

2014L00966-TP-4 **3.3**

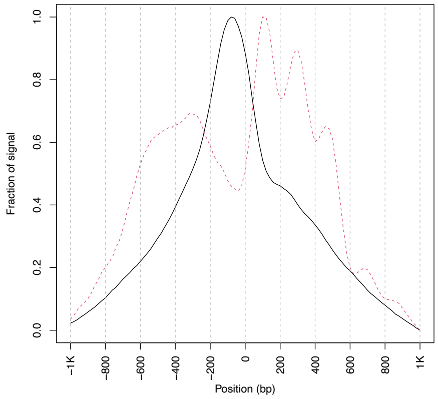


2014L00966-TP-8 **3.0**

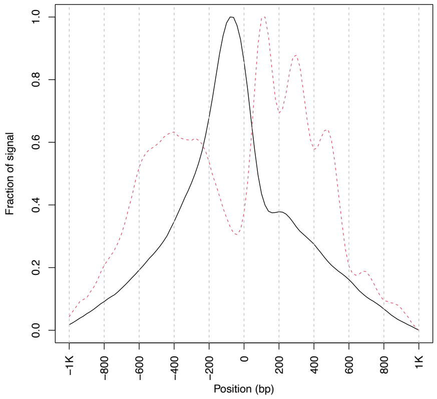


2014L00966-TP-12 **2.6**

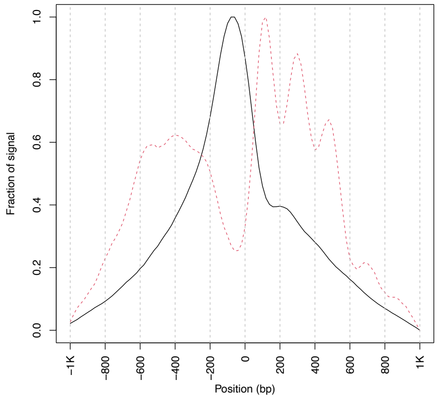


2014L00966-TP-16 **4.0**

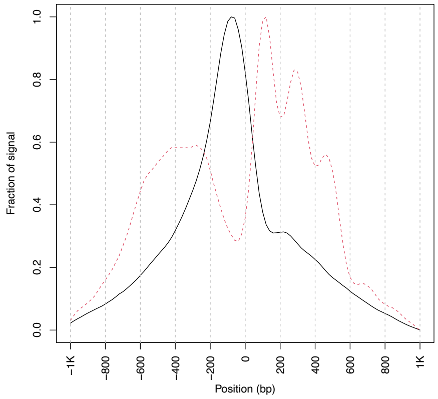


2014L00966-TP-24 **2.5**

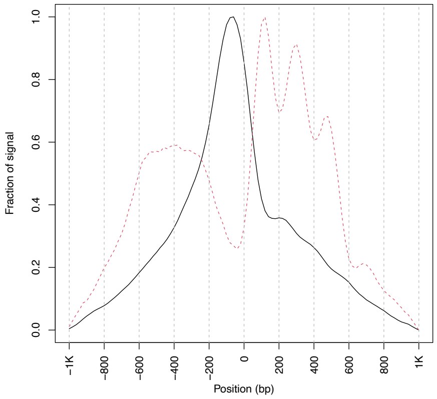


2014L00966-TP-28 **2.8**

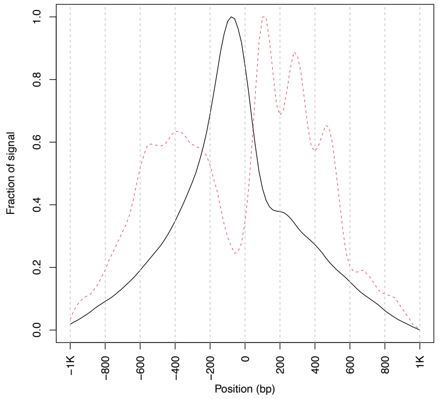


2014L00966-TP-32 **2.6**

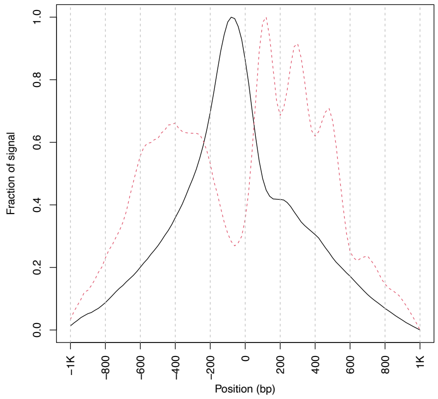


2014L00966-TP-36 **2.8**

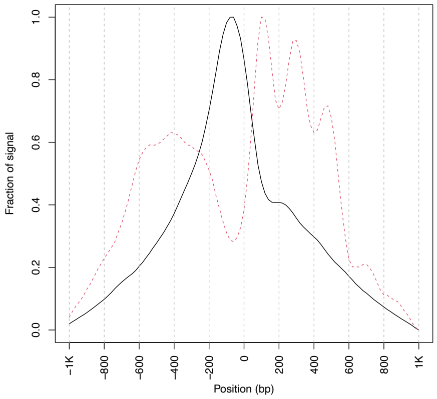


2014L00966-TP-40 **3.6**

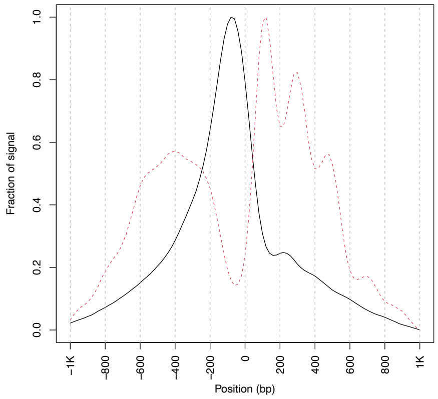

2014L00966-TP-44 **3.7**

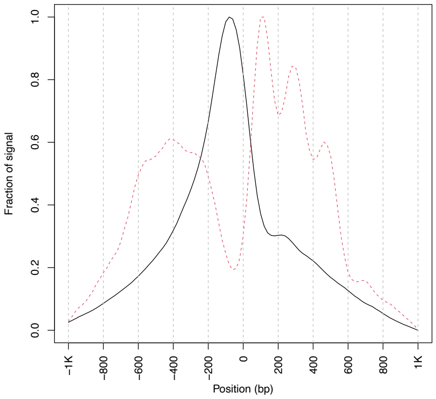


2014L00966-TP-48 **3.0**

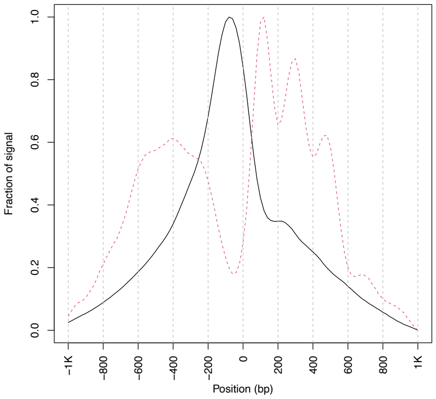

2014L01145-TP-0 **3.5**

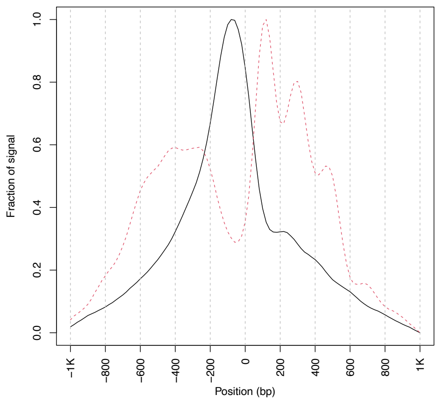


2014L01145-TP-4 **4.1**

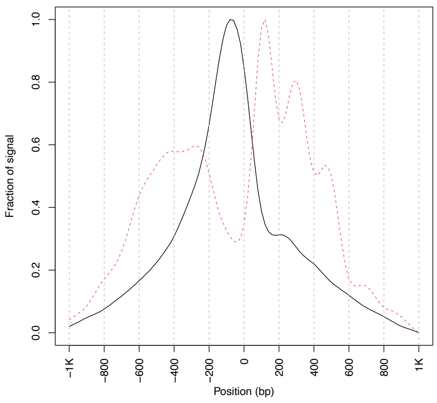


2014L01145-TP-8 **3.5**

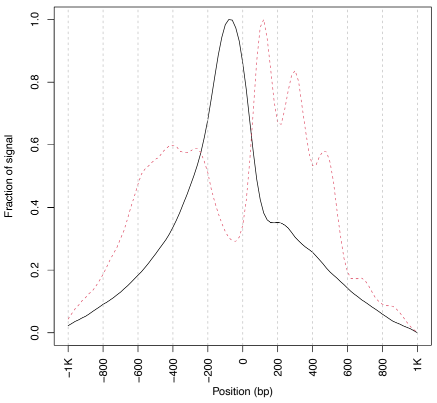


2014L01145-TP-12 **3.8**

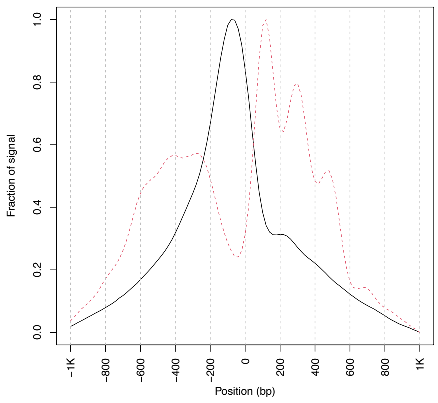


2014L01145-TP-16 **3.7**

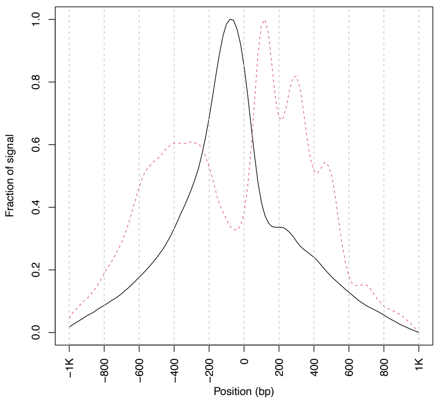


2014L01145-TP-20 **4.5**

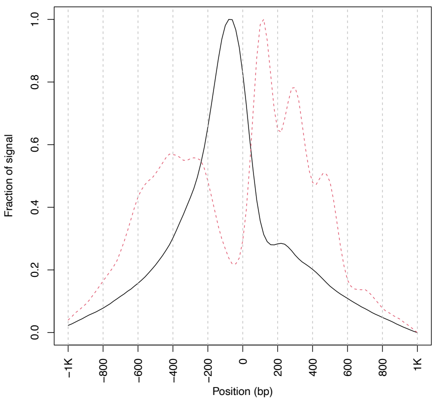


2014L01145-TP-24 **4.7**

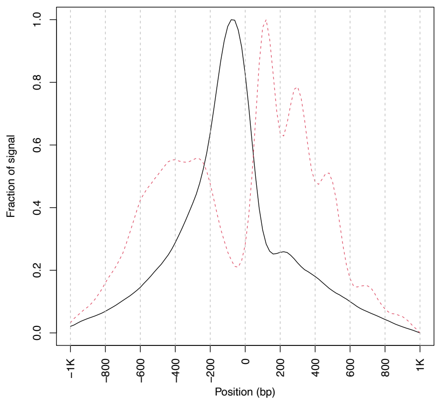


2014L01145-TP-28 **3.5**

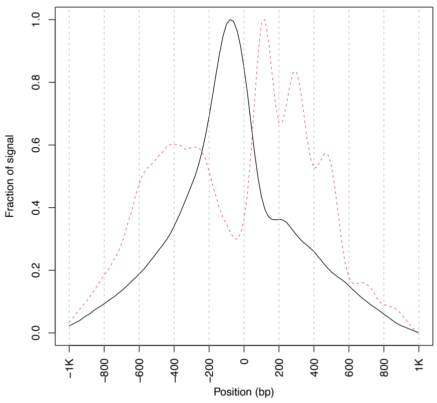


2014L01145-TP-32 **3.1**

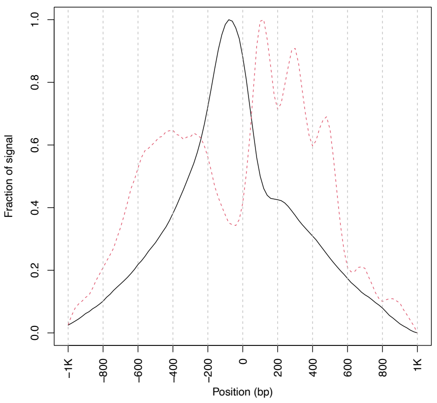

2014L01145-TP-36 **3.5**

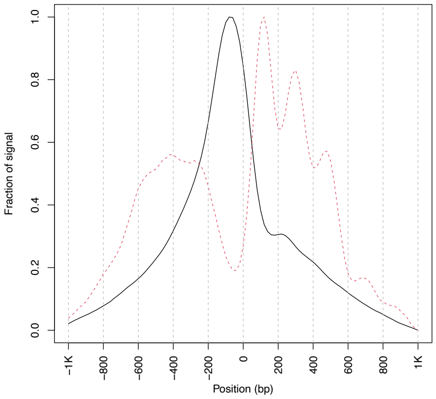


2014L01145-TP-40 **4.1**

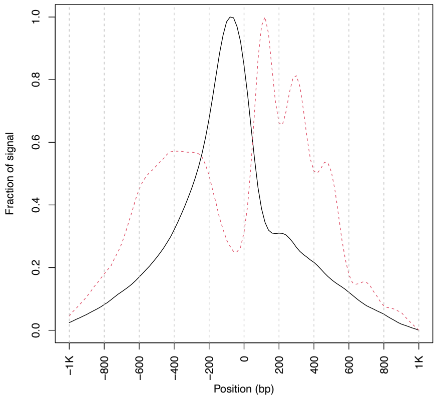


2014L01145-TP-44 **4.2**

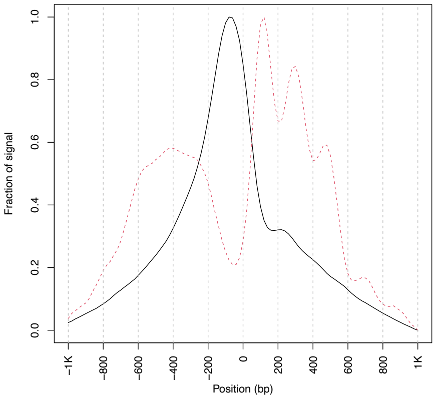


2014L01145-TP-48 **3.9**

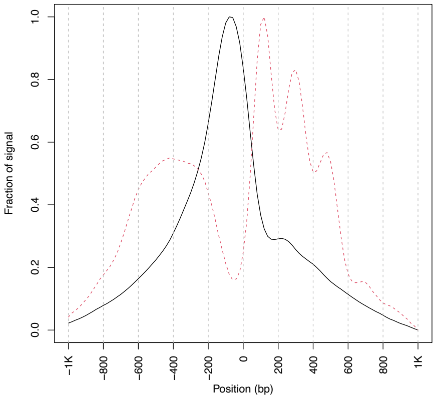

2014L01249-TP-0 **4.1**

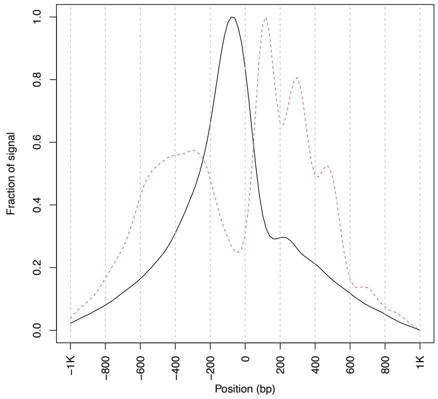


2014L01249-TP-4 **4.2**

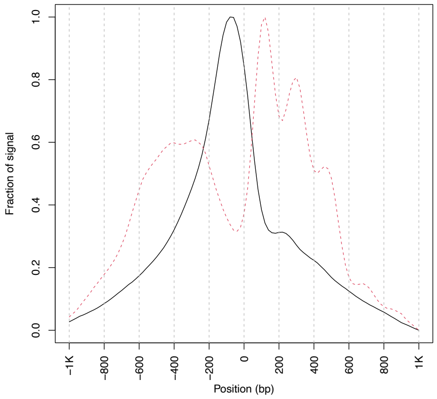


2014L01249-TP-8 **3.5**

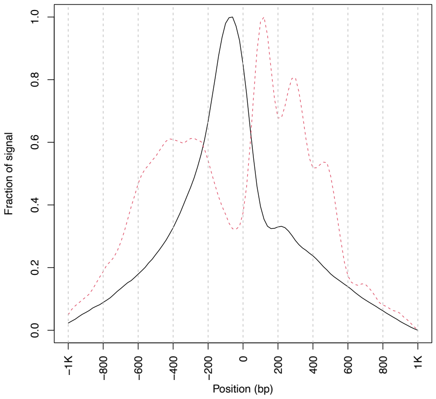


2014L01249-TP-12 **4.1**

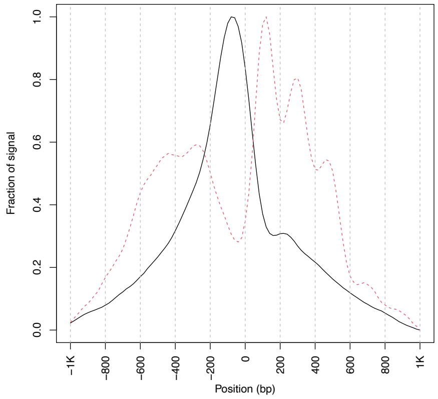


2014L01249-TP-16 **3.8**

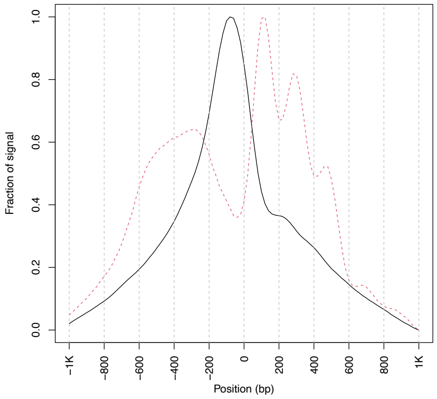


2014L01249-TP-20 **4.6**

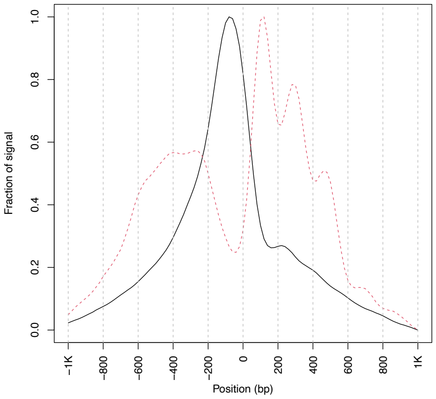


2014L01249-TP-24 **4.1**

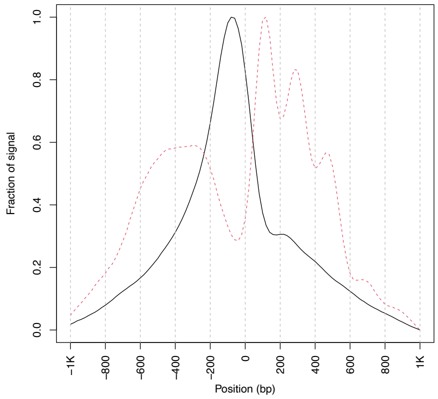


2014L01249-TP-28 **3.2**

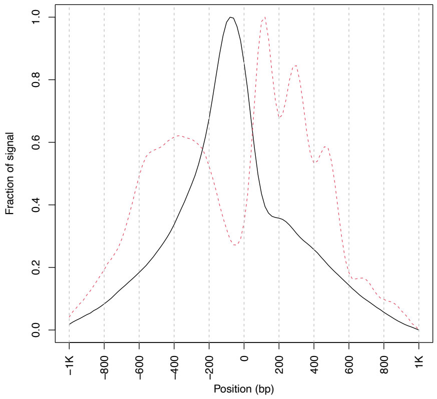


2014L01249-TP-32 **2.7**

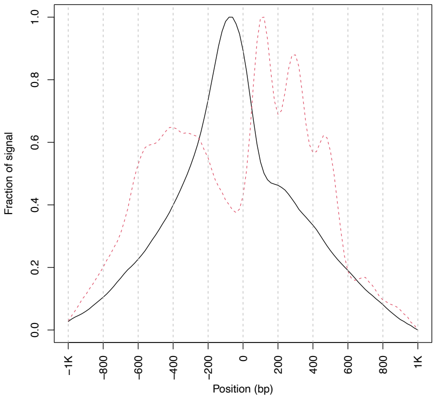

2014L01249-TP-36 **3.7**

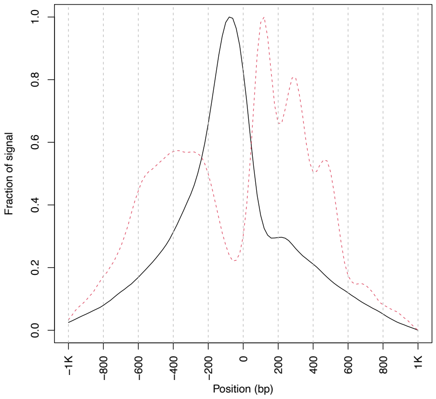


2014L01249-TP-40 **4.7**

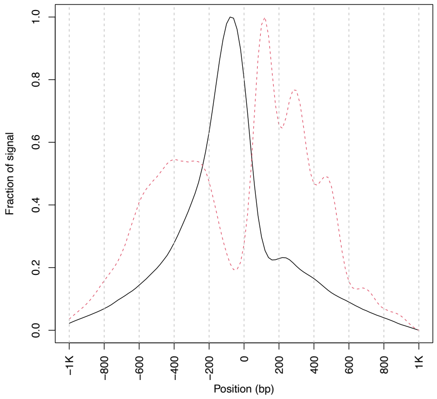


2014L01249-TP-44 **4.2**

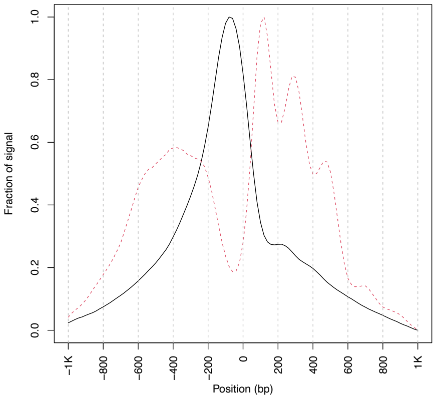


2014L01249-TP-48 **3.6**

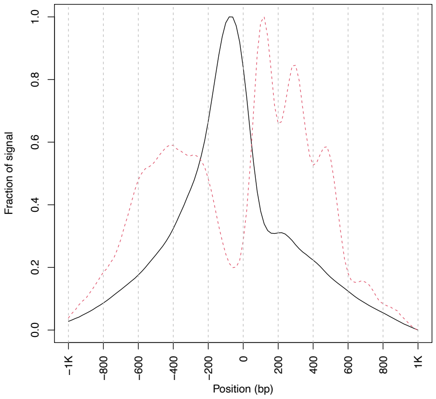
