## Supplementary figures and images for "Fibroblasts as an in vitro model of circadian genetic and genomic studies: A temporal analysis"

### heatmap_cluster 1 .png

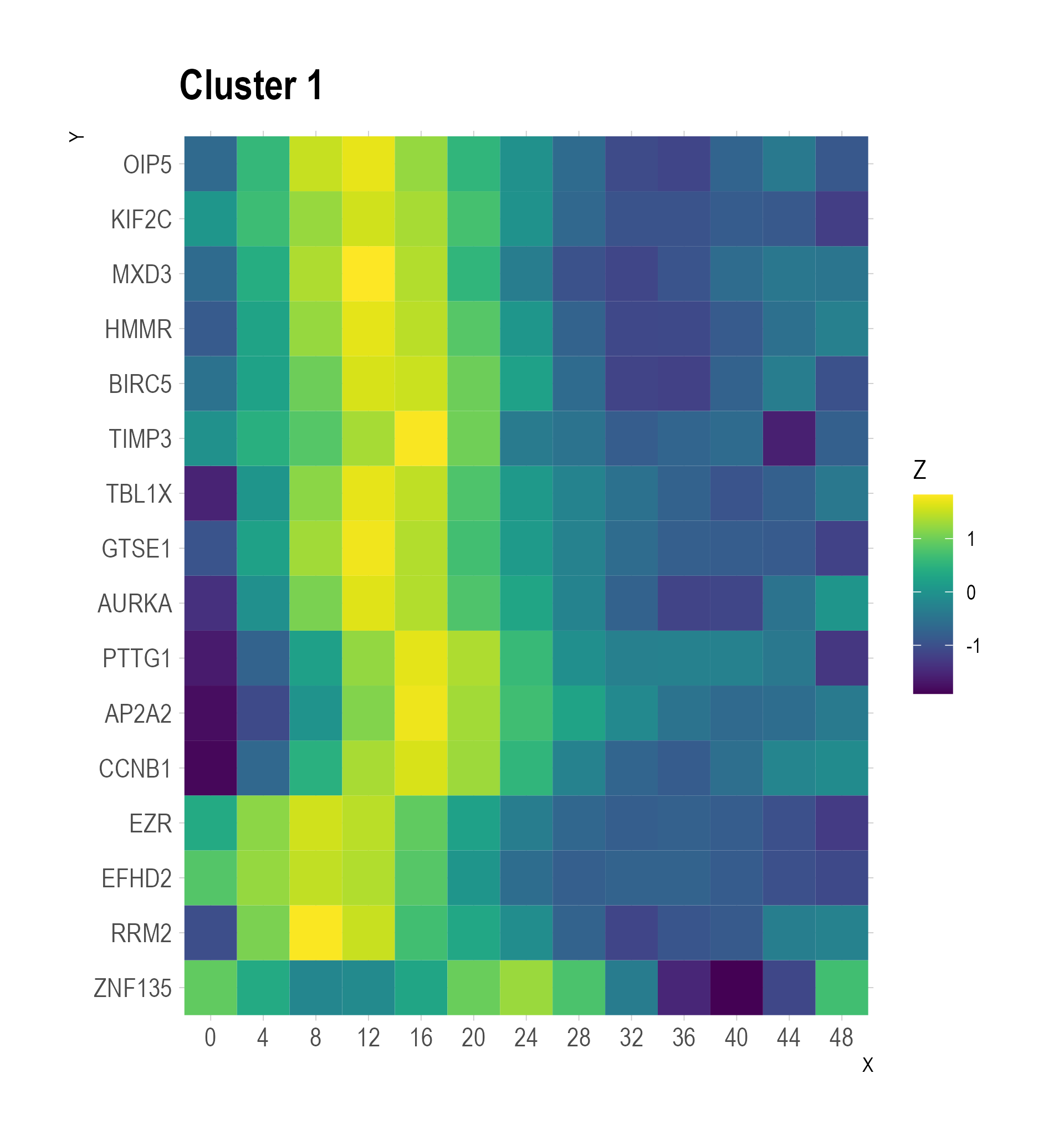

### heatmap_cluster 2 .png

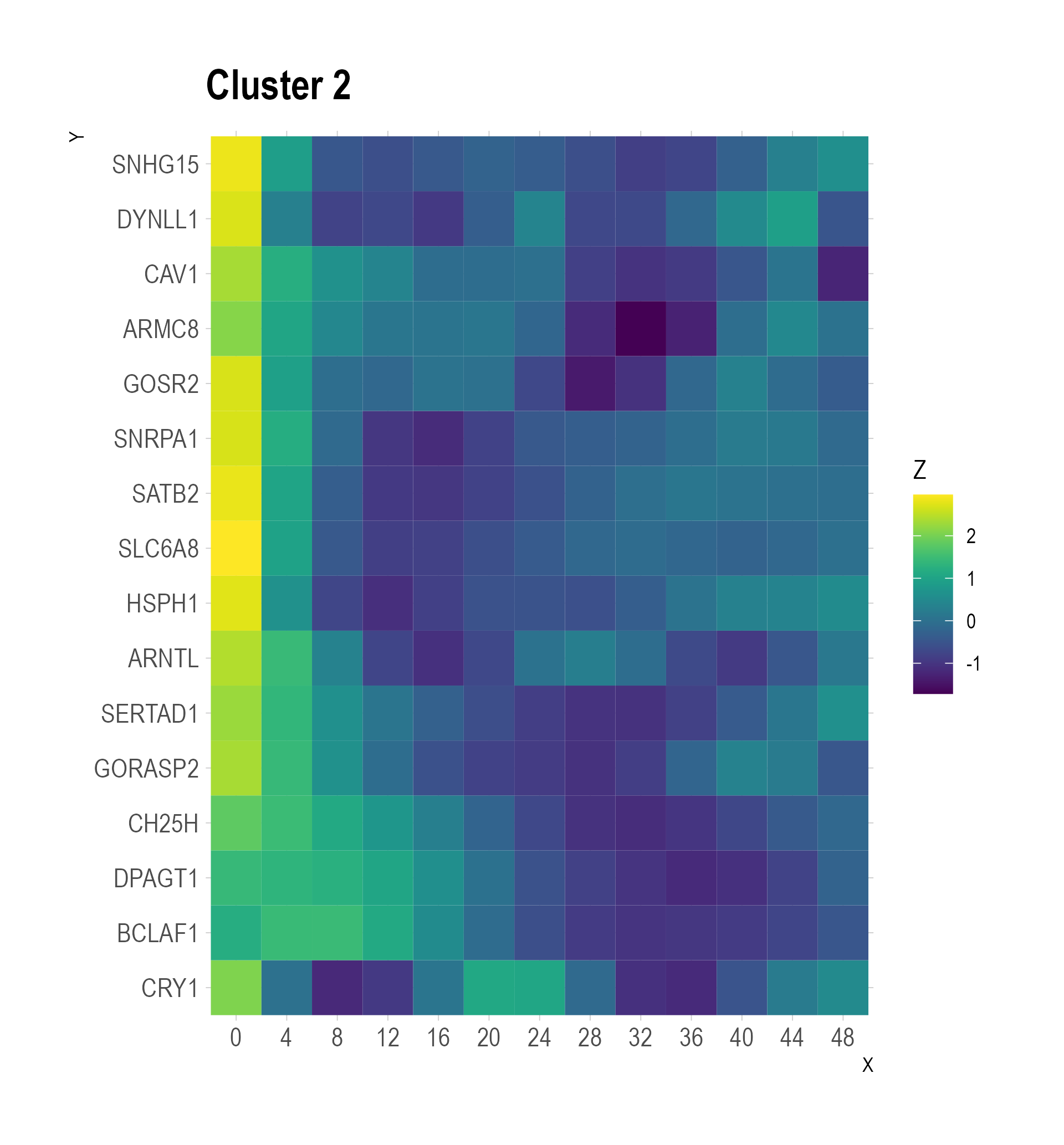

### heatmap_cluster 3 .png

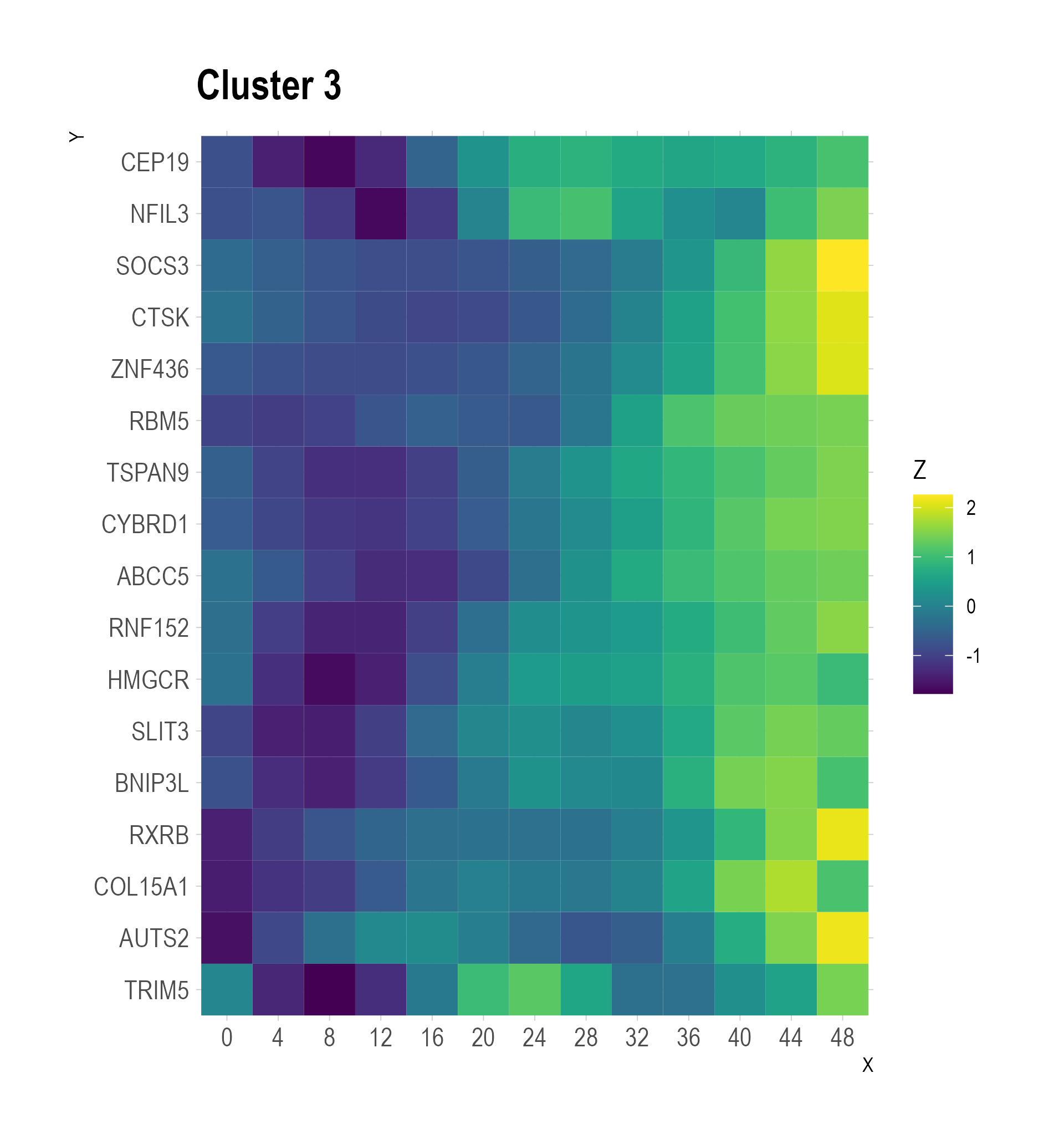

### heatmap_cluster 4 .png

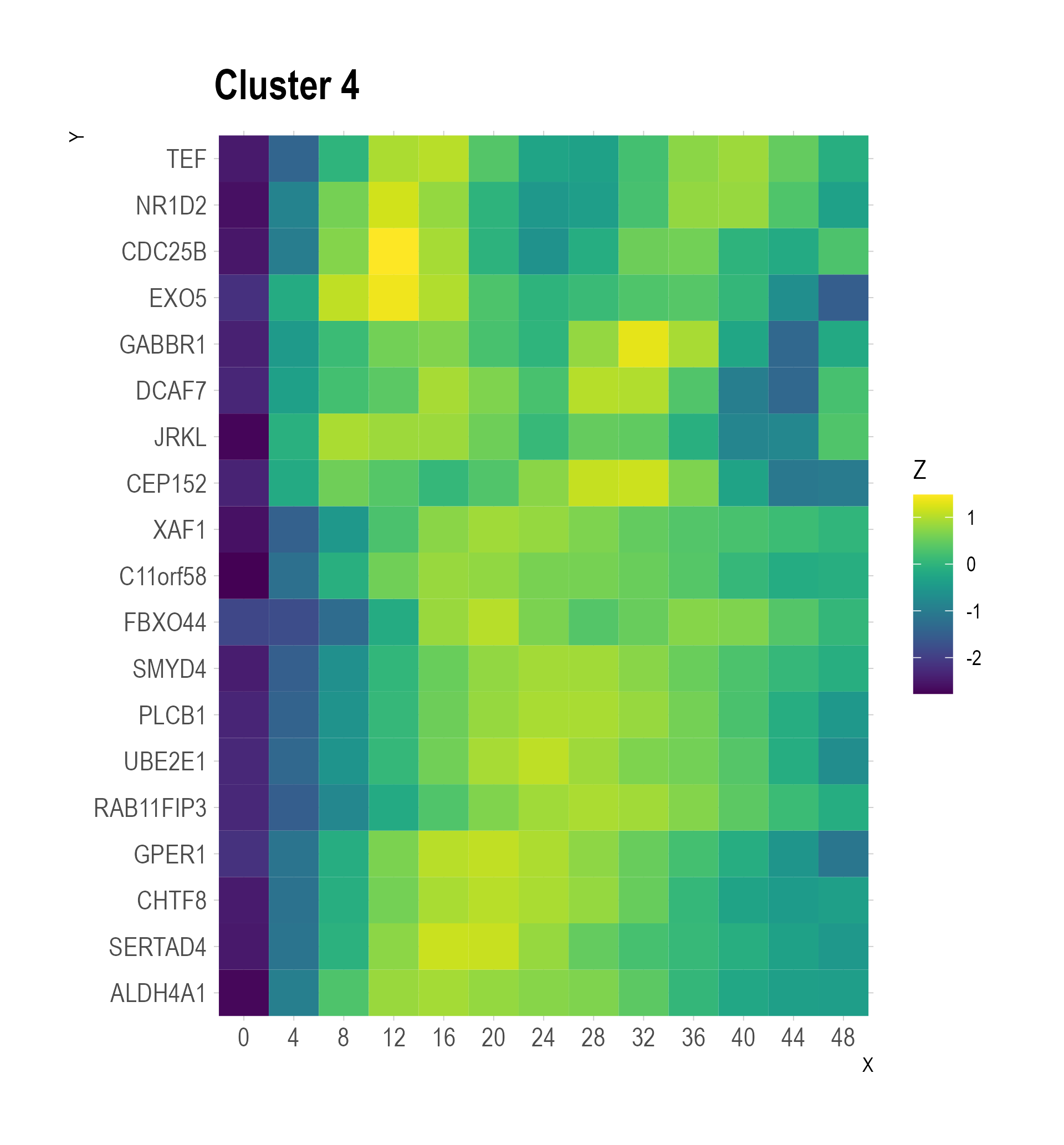

### heatmap_cluster 5 .png

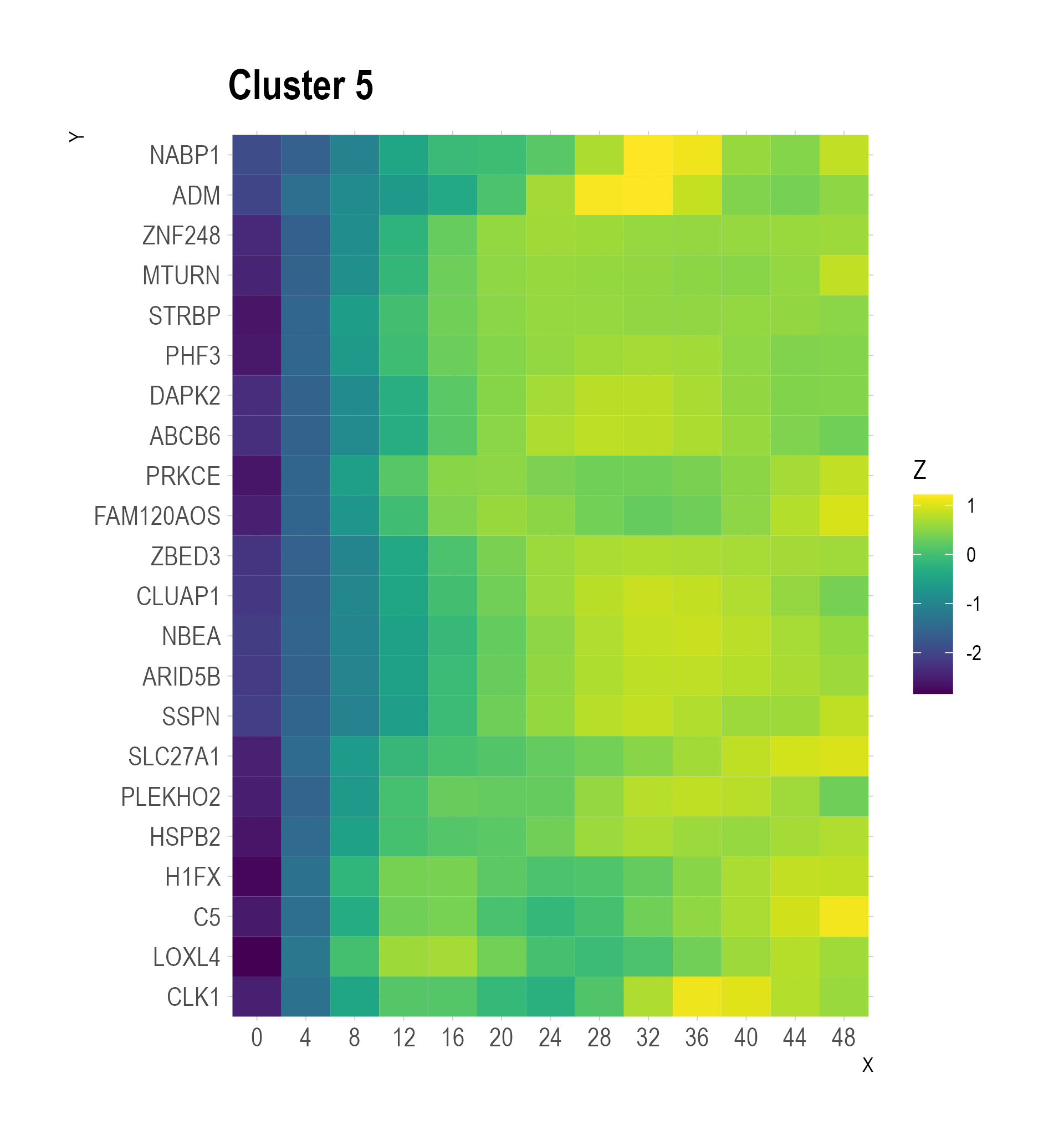

### heatmap_cluster 6 .png

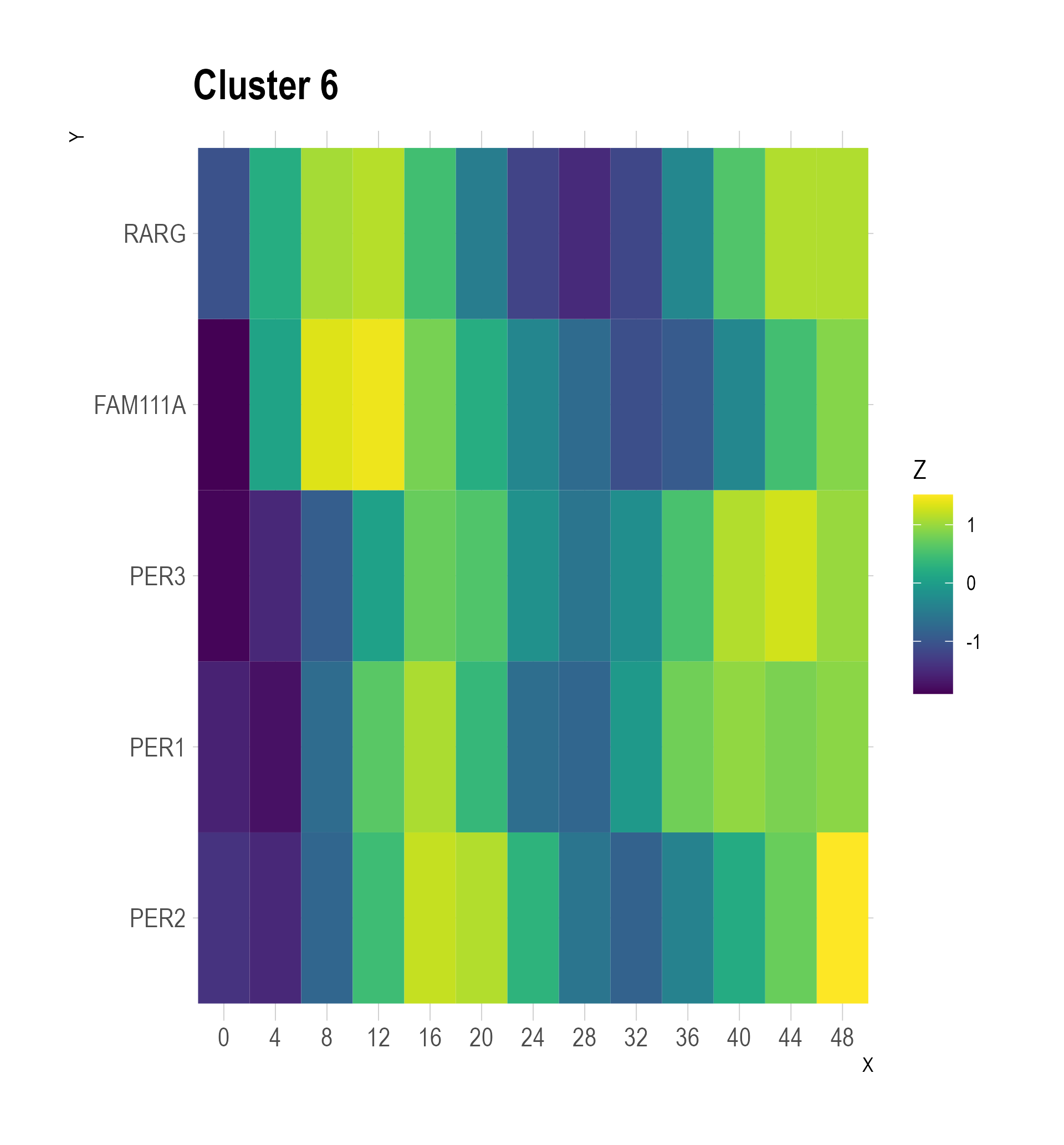

### heatmap_cluster 7 .png

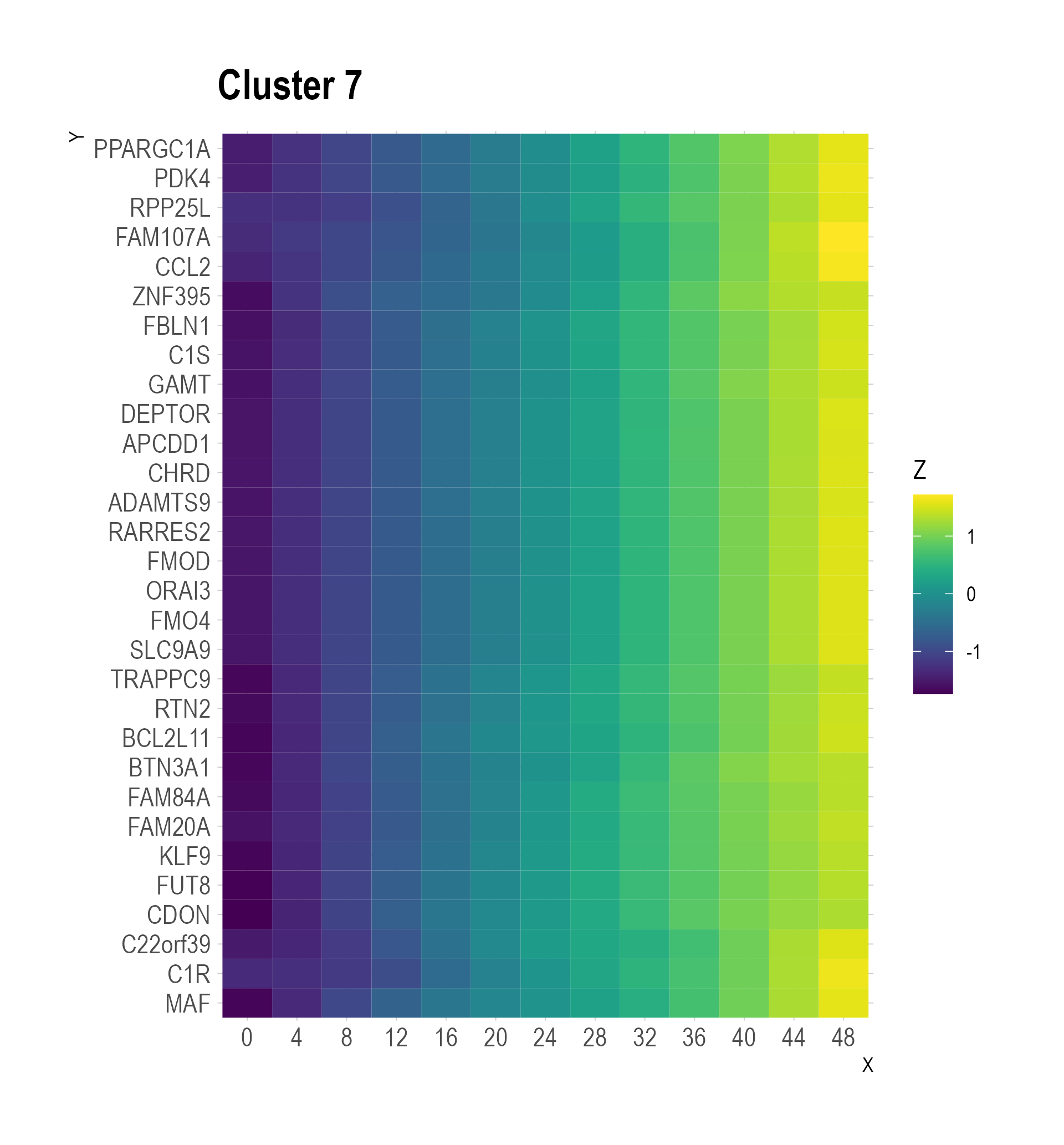

### heatmap_cluster 8 .png

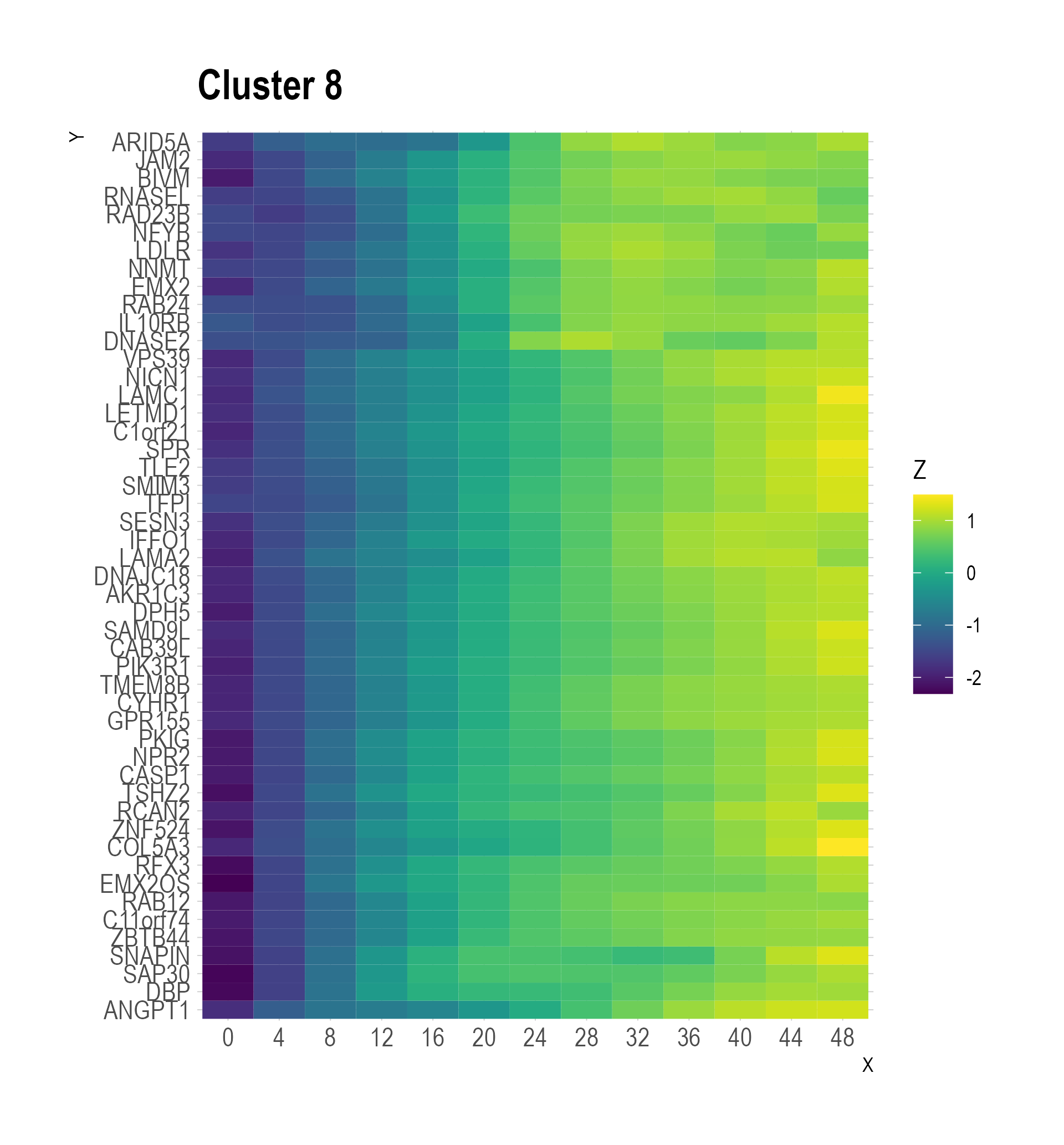

### heatmap_cluster 9 .png

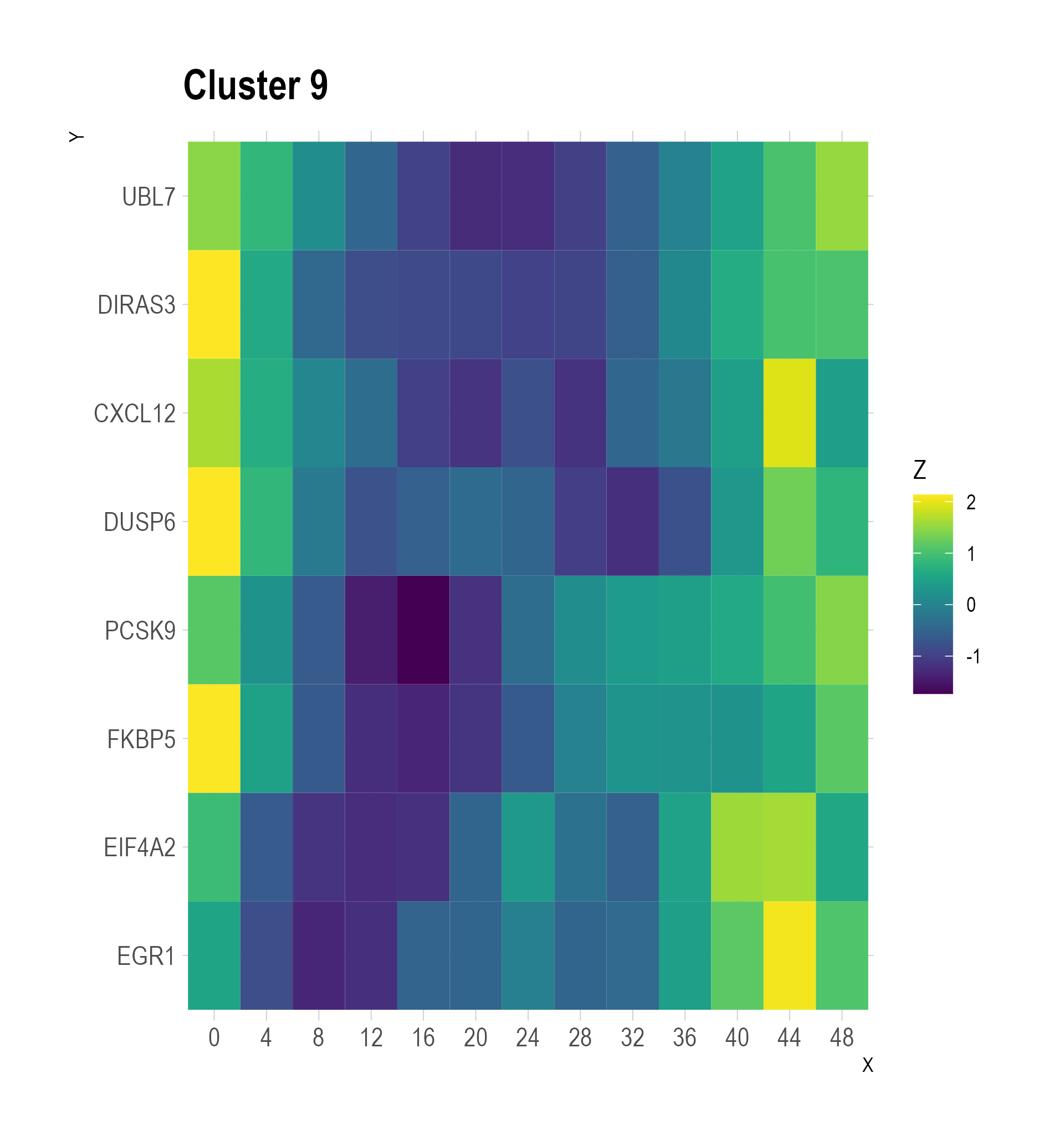

### heatmap_cluster 10 .png

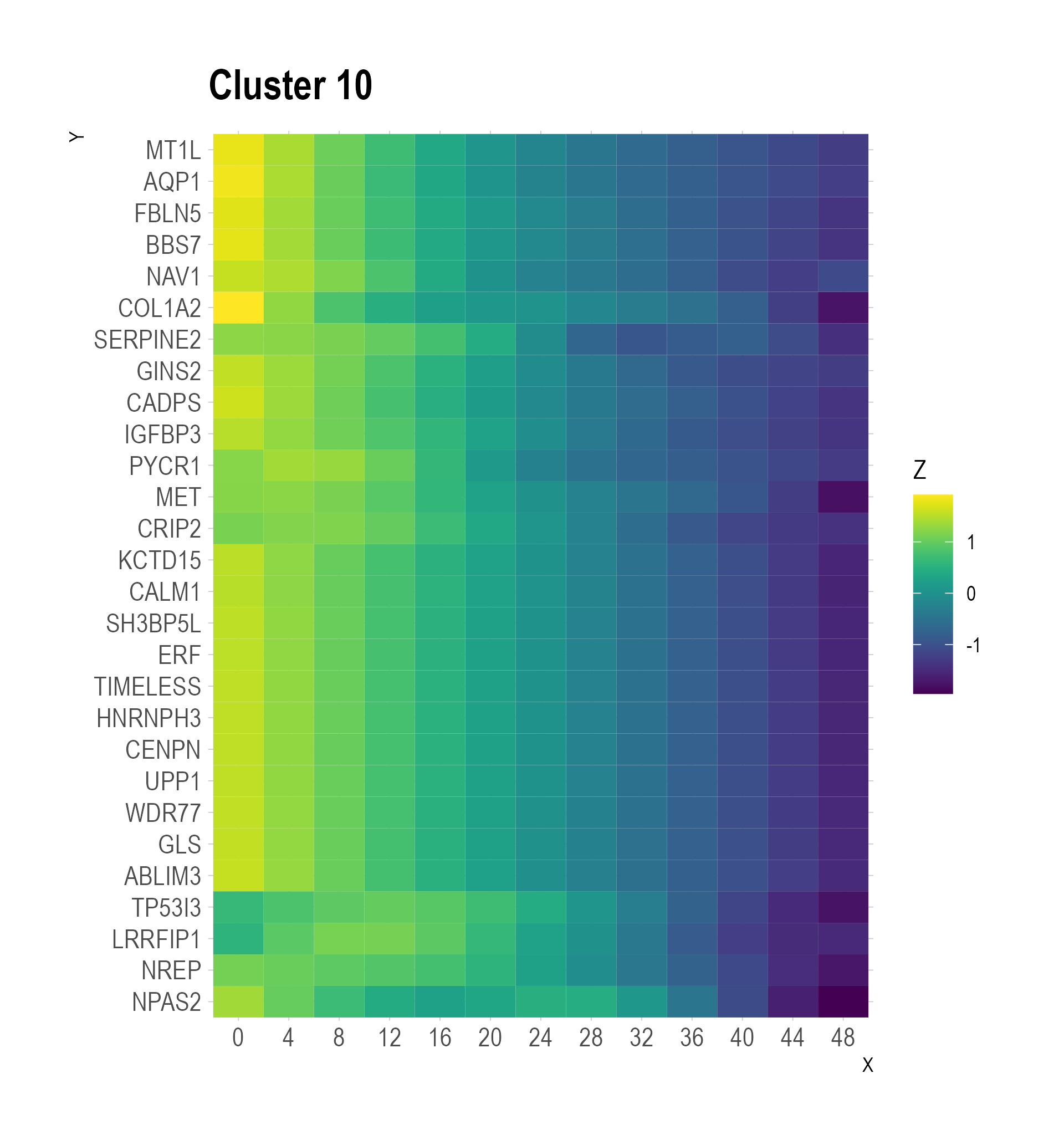

### heatmap_cluster 11 .png

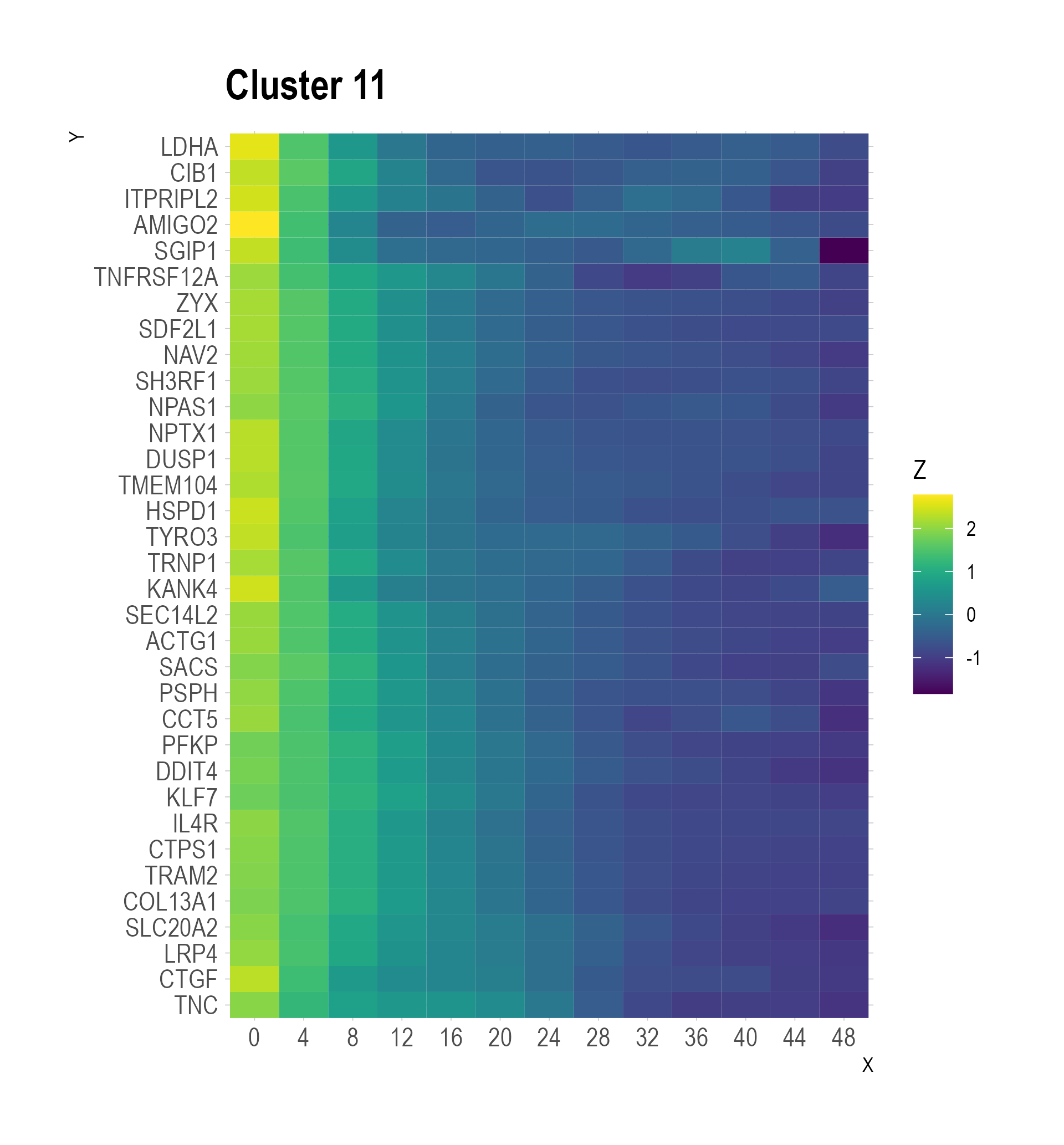

### heatmap_cluster 12 .png

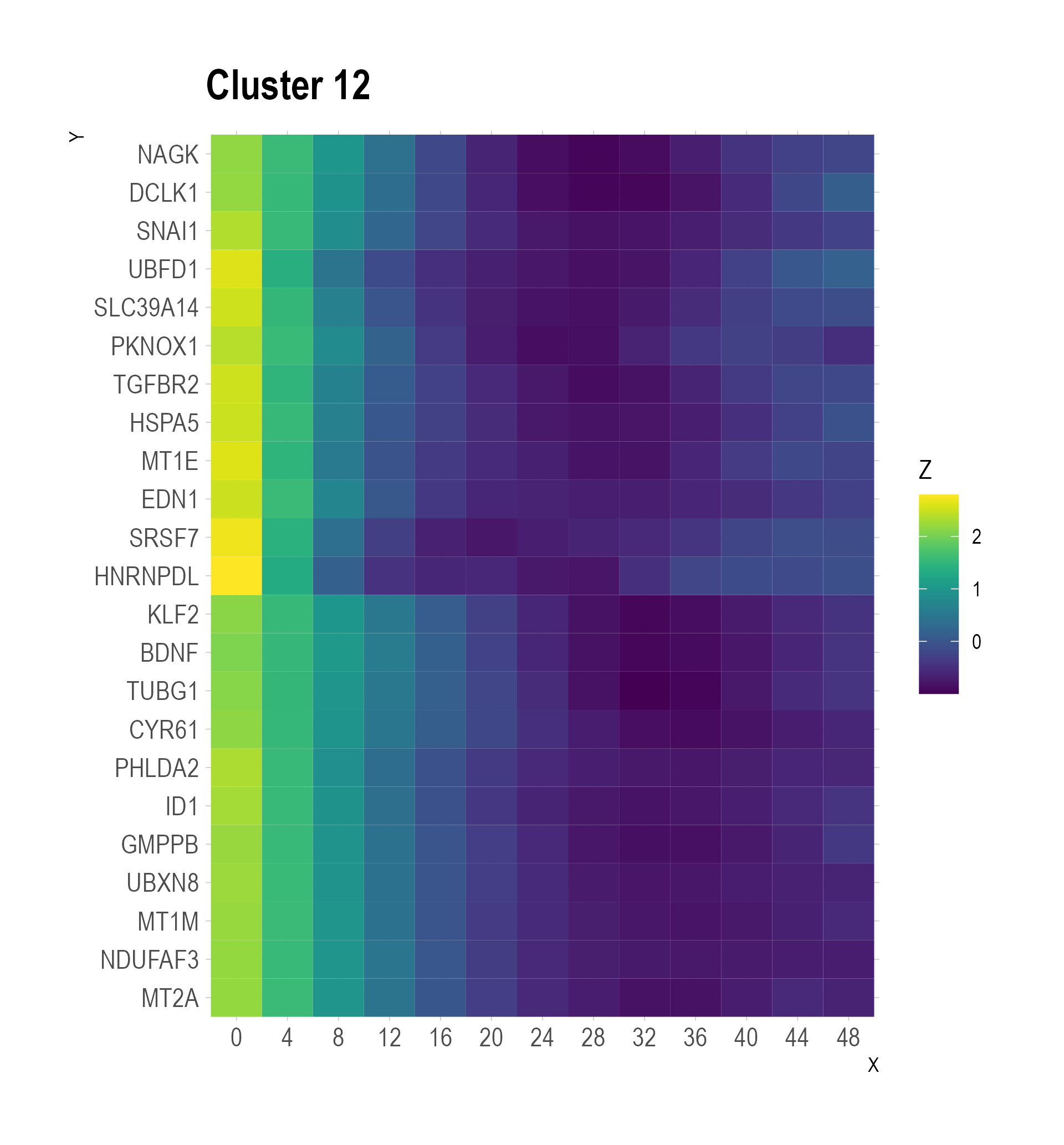
